# Origin and cross-species transmission of bat coronaviruses in China

**DOI:** 10.1101/2020.05.31.116061

**Authors:** Alice Latinne, Ben Hu, Kevin J. Olival, Guangjian Zhu, Libiao Zhang, Hongying Li, Aleksei A. Chmura, Hume E. Field, Carlos Zambrana-Torrelio, Jonathan H. Epstein, Bei Li, Wei Zhang, Lin-Fa Wang, Zheng-Li Shi, Peter Daszak

## Abstract

Bats are presumed reservoirs of diverse coronaviruses (CoVs) including progenitors of Severe Acute Respiratory Syndrome (SARS)-CoV and SARS-CoV-2, the causative agent of COVID-19. However, the evolution and diversification of these coronaviruses remains poorly understood. We used a Bayesian statistical framework and sequence data from all known bat-CoVs (including 630 novel CoV sequences) to study their macroevolution, cross-species transmission, and dispersal in China. We find that host-switching was more frequent and across more distantly related host taxa in alpha-than beta-CoVs, and more highly constrained by phylogenetic distance for beta-CoVs. We show that inter-family and -genus switching is most common in Rhinolophidae and the genus *Rhinolophus*. Our analyses identify the host taxa and geographic regions that define hotspots of CoV evolutionary diversity in China that could help target bat-CoV discovery for proactive zoonotic disease surveillance. Finally, we present a phylogenetic analysis suggesting a likely origin for SARS-CoV-2 in *Rhinolophus* spp. bats.

## Introduction

Coronaviruses (CoVs) are RNA viruses causing respiratory and enteric diseases with varying pathogenicity in humans and animals. All CoVs known to infect humans are zoonotic, or of animal origin, with many thought to originate in bat hosts^1,2^. Due to their large genome size (the largest non-segmented RNA viral genome), frequent recombination and high genomic plasticity, CoVs are prone to cross-species transmission and are able to rapidly adapt to new hosts^1,3^. This phenomenon is thought to have led to the emergence of a number of CoVs affecting livestock and human health^4–9^. Three of these causing significant outbreaks originated in China during the last two decades. Severe Acute Respiratory Syndrome (SARS)-CoV emerged first in humans in Guangdong province, southern China, in 2002 and spread globally, causing fatal respiratory infections in close to 800 people^10–12^. Subsequent investigations identified horseshoe bats (genus *Rhinolophus*) as the natural reservoirs of SARS-related CoVs and the likely origin of SARS-CoV^13–16^. In 2016, Swine Acute Diarrhea Syndrome (SADS)-CoV caused the death of over 25,000 pigs in farms within Guangdong province^17^. This virus appears to have originated within *Rhinolophus* spp. bats, and belongs to the HKU2-CoV clade previously detected in bats in the region^17–19^. In 2019, a novel coronavirus (SARS-CoV-2) caused an outbreak of respiratory illness (COVID-19) first detected in Wuhan, Hubei province, China, which has since become a pandemic. This emerging human virus is closely related to SARS-CoV, and also appears to have originated in horseshoe bats^20,21^ - with its full genome 96% similar to a viral sequence reported from *Rhinolophus affinis*^20^. Closely related sequences were also identified in Malayan pangolins^22,23^.

A growing body of research has identified bats as the evolutionary sources of SARS- and Middle East Respiratory Syndrome (MERS)-CoVs ^13,14,24–26^, and as the source of progenitors for the human CoVs, NL63 and 229E^27,28^. The emergence of SARS-CoV-2 further underscores the importance of bat-origin CoVs to global health, and understanding their origin and cross-species transmission is a high priority for pandemic preparedness^20,29^. Bats harbor the largest diversity of CoVs among mammals and two CoV genera, alpha- and beta-CoVs (α- and β-CoVs), have been widely detected in bats from most regions of the world^30,31^. Bat-CoV diversity seems to be correlated with host taxonomic diversity globally, the highest CoV diversity being found in areas with the highest bat species richness^32^. Host switching of viruses over evolutionary time is an important mechanism driving the evolution of bat coronaviruses in nature and appears to vary geographically^32,33^. However, detailed analyses of host-switching have been hampered by incomplete or opportunistic sampling, typically with relatively low numbers of viral sequences from any given region^34^.

China has a rich bat fauna, with more than 100 described bat species and several endemic species representing both the Palearctic and Indo-Malay regions^35^. Its situation at the crossroads of two zoogeographic regions heightens China’s potential to harbor a unique and distinctive CoV diversity. Since the emergence of SARS-CoV in 2002, China has been the focus of an intense viral surveillance and a large number of diverse bat-CoVs has been discovered in the region^36–44^. However, the macroevolution of CoVs in their bat hosts in China and their cross-species transmission dynamics remain poorly understood.

In this study, we analyze an extensive field-collected dataset of bat-CoV sequences from across China. We use a phylogeographic Bayesian statistical framework to reconstruct virus transmission history between different bat host species and virus spatial spread over evolutionary time. Our objectives were to compare the macroevolutionary patterns of α- and β-CoVs and identify the hosts and geographical regions that act as centers of evolutionary diversification for bat-CoVs in China. These analyses aim to improve our understanding of how CoVs evolve, diversify, circulate among, and transmit between bat families and genera to help identify bat hosts and regions where the risk of CoV spillover is the highest.

## Results

### Taxonomic and geographic sampling

We generated 630 partial sequences (440 nt) of the *RNA-dependent RNA polymerase* (*RdRp*) gene from bat rectal swabs collected in China and added 608 bat-CoV and eight pangolin CoV sequences from China available in GenBank or GISAID to our datasets (list of GenBank and GISAID accession numbers available in Supplementary Note 1). For each CoV genus, two datasets were created: one including all bat-CoV sequences with known host (host dataset) and one including all bat-CoV sequences with known sampling location at the province level (geographic dataset). To create a geographically discrete partitioning scheme that was more ecologically relevant than administrative borders for our phylogeographic reconstructions, we defined six zoogeographic regions within China by clustering provinces with similar mammalian diversity using hierarchical clustering^45^ (see Methods): South western region (SW), Northern region (NO), Central northern region (CN), Central region (CE), Southern region (SO) and Hainan island (HI) (Fig. 1 and Supplementary Fig. 1).

**Fig. 1.**
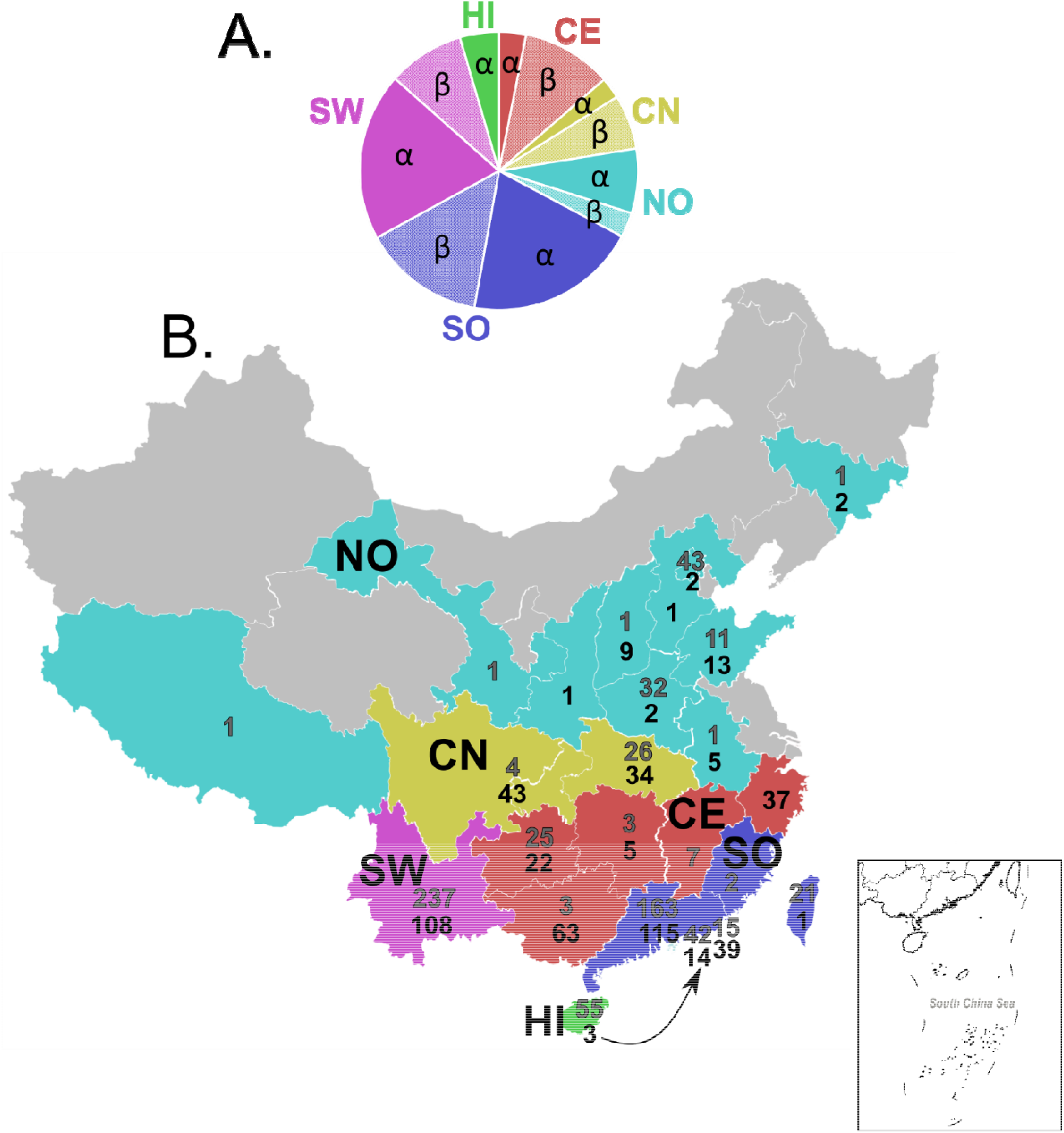
Geographic sampling. Pie chart (A) showing the number of sequences of each CoV genus (alpha-CoVs and beta-CoVs) available for each zoogeographic region and map of China provinces (B) showing the number of *RdRp* sequences available for each province, in bold grey for alpha-CoVs and black for beta-CoVs. Province colors correspond to the zoogeographic region to which they belong: NO, Northern region; CN, Central northern region; SW, South western region; CE, Central region; SO, Southern region; HI, Hainan island. The three beta-CoV sequences from HI were included in the SO region. Provinces colored in grey are those where CoV sequences are not available.

Our host datasets included 701 α-CoV sequences (353 new sequences, including 102 new SADSr-CoV sequences (*Rhinacovirus*) from 41 bat species (14 genera, five families) and 528 β-CoV sequences (273 new sequences, including 97 new SARSr-CoV sequences (*Sarbecovirus*) from 31 bat species (15 genera, four families) (Supplementary Table 1). Our geographic datasets included 677 α-CoV sequences from six zoogeographic regions (22 provinces) and 503 β-CoV sequences from five zoogeographic regions (21 provinces) (Fig. 1). As some regions or hosts were overrepresented in our datasets, we also created and ran our analyses using a more uniform subset of our sequence data that included ~30 randomly-selected sequences per host family or region to mitigate sampling and surveillance intensity bias.

### Ancestral hosts and cross-species transmission

We used a Bayesian discrete phylogeographic approach implemented in BEAST^46^ to reconstruct the ancestral host of each node in the phylogenetic tree using bat host family as a discrete character state. The phylogenetic reconstructions for α-CoVs in China suggest an evolutionary origin within rhinolophid and vespertilionid bats (Fig. 2A). The first α-CoV lineage to diverge historically corresponds to the subgenus *Rhinacovirus* (L1), originating within rhinolophid bats, and includes sequences related to HKU2-CoV and SADS-CoV (Supplementary Fig. 2). Then several lineages, labelled L2 to L7, emerged from vespertilionid bats (Fig. 2A). The subgenus *Decacovirus* (L2) includes sequences mostly associated with the Rhinolophidae and Hipposideridae and related to HKU10-CoV (Supplementary Fig. 3), while the subgenera *Myotacovirus* (L3) and *Pedacovirus* (L5) as well as an unidentified lineage (L4) include CoVs mainly from vespertilionid bats and related to HKU6-, HKU10-, and 512-CoVs (Supplementary Fig. 4-5). Finally, a well-supported node comprises the subgenera *Nyctacovirus* (L6) from vespertilionid bats and *Minunacovirus* (L7) from miniopterid bats, and includes HKU7-, HKU8-, 1A-, and 1B-CoVs (Supplementary Fig. 6). These seven α-CoV lineages are mostly associated with a single host family but each also included several sequences identified from other bat families (Fig. 2A, Supplementary Fig. 2-6 and Supplementary Table 1), suggesting frequent cross-species transmission events have occurred among bats. Ancestral host reconstructions based on the random data subset, to normalize sampling effort, gave very similar results with rhinolophids and vespertilionids being the most likely ancestral hosts of most α-CoV lineages too (Supplementary Fig. 7A). However, the topology of the tree based on the random subset was slightly different as the lineage L5 was paraphyletic.

**Fig. 2.**
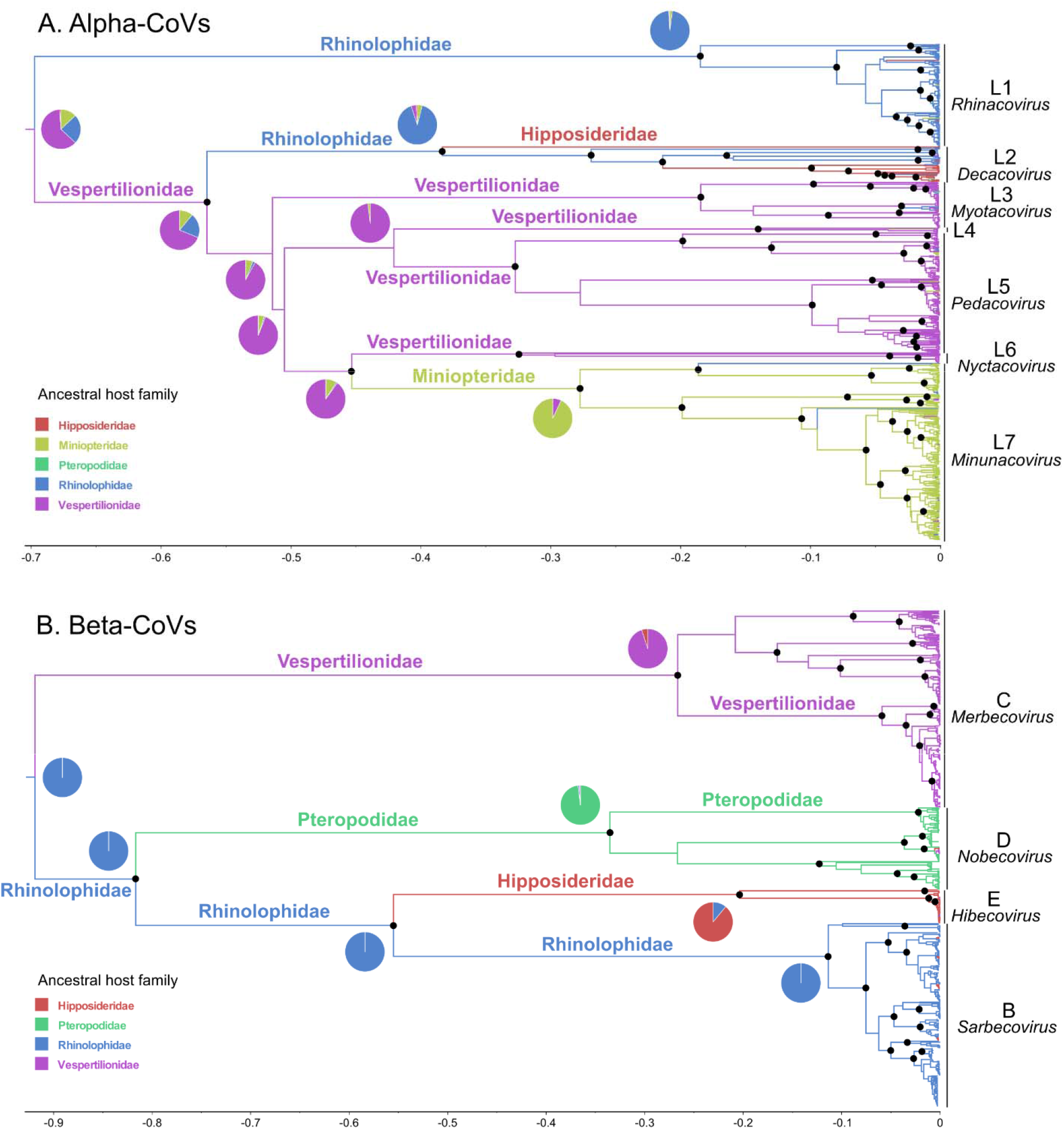
Phylogenetic trees and ancestral host reconstructions. Alpha-CoV (A) and beta-CoV (B) maximum clade credibility annotated trees using complete datasets of *RdRp* sequences and bat host family as discrete character state. Pie charts located at the root and close to the deepest nodes show the state posterior probabilities for each bat family. Branch colors correspond to the inferred ancestral family with the highest probability. Branch lengths are scaled according to relative time units (clock rate = 1.0). Well-supported nodes (posterior probability > 0.95) are indicated with a black dot. The ICTV approved CoV subgenera were highlighted: *Rhinacovirus* (L1), *Decacovirus* (L2), *Myotacovirus* (L3), *Pedacovirus* (L5), *Nyctacovirus* (L6), *Minunacovirus* (L7) and an unidentified lineage (L4) for alpha-CoVs; and *Merbecovirus* (Lineage C), *Nobecovirus* (lineage D), *Hibecovirus* (lineage E) and *Sarbecovirus* (Lineage B) for beta-CoVs.

Chinese β-CoVs likely originated from vespertilionid and rhinolophid bats (Fig. 2B). The MCC tree was clearly structured into four main lineages: *Merbecovirus* (Lineage C), including MERS-related (MERSr-) CoVs, HKU4- and HKU5-CoVs and strictly restricted to vespertilionid bats (Supplementary Fig. 8); *Nobecovirus* (lineage D), originating from pteropodid bats and corresponding to HKU9-CoV (Supplementary Fig. 9); *Hibecovirus* (lineage E) comprising sequences isolated in hipposiderid bats (Supplementary Fig. 10) and *Sarbecovirus* (Lineage B) including sequences related to HKU3- and SARS-related (SARSr-) CoVs originating in rhinolophid bats (Supplementary Fig. 11). We show that SARS-CoV-2 forms a divergent clade within *Sarbecovirus* and is most closely related to viruses sampled from *Rhinolophus malayanus* and *R. affinis* and from Malayan pangolins (*Manis javanica*) (Fig. 3). Similar tree topology and ancestral host inference were obtained with the random subset (Supplementary Fig. 7B).

**Fig. 3.**
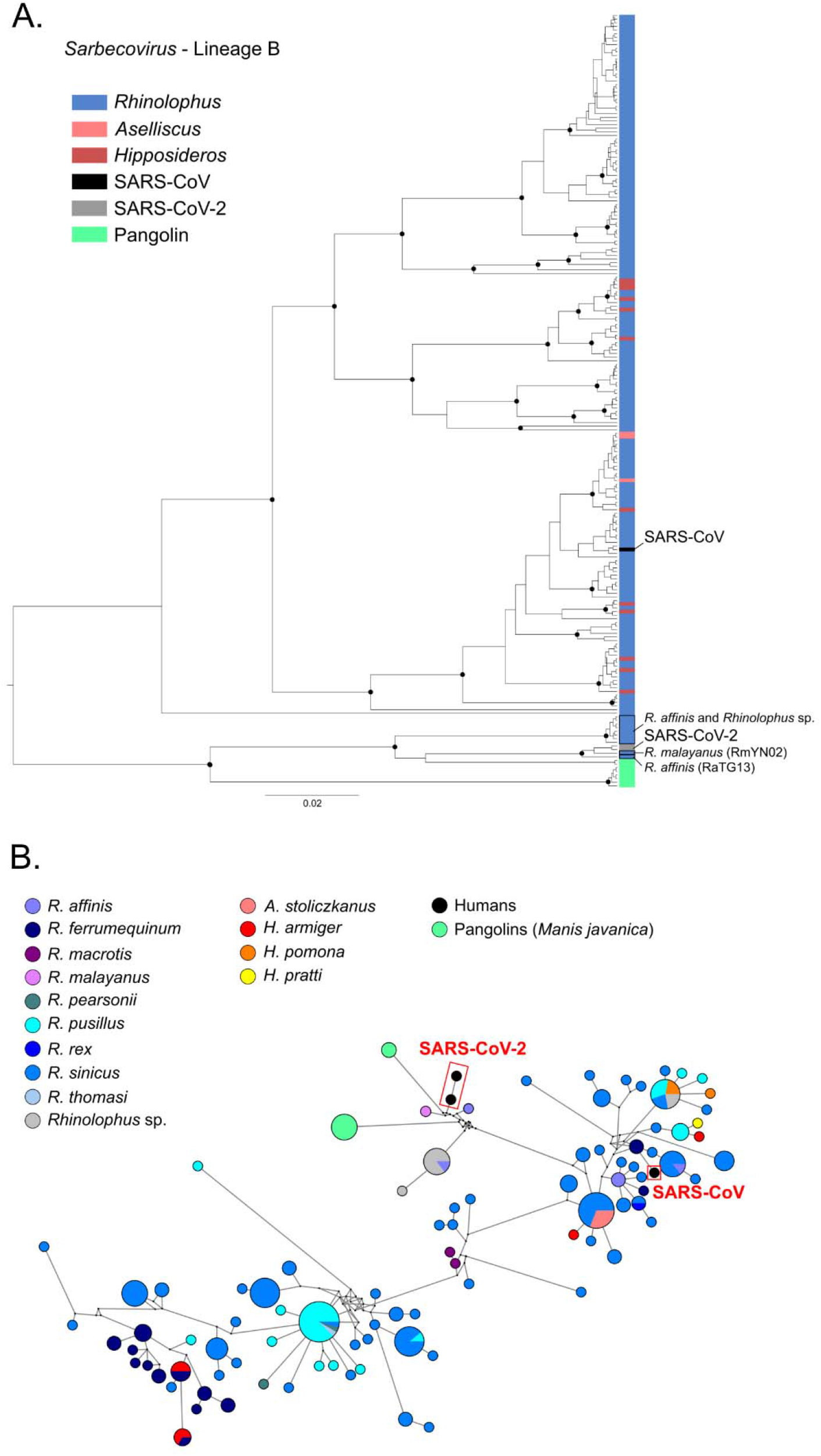
Phylogenetic relationships within the *Sarbecovirus* subgenus (beta-CoVs). Maximum clade credibility tree (A) including 202 *RdRp* sequences from the *Sarbecovirus* subgenus isolated in bats, two sequences of SARS-CoV-2 and one sequence of SARS-CoV isolated in humans and eight sequences isolated in Malayan pangolins (*Manis javanica*). Well-supported nodes (posterior probability > 0.95) are indicated with a black dot. Tip colors correspond to the host genus, SARS-CoV-2 sequences and SARS-CoV sequence are highlighted in grey and black, respectively. Median-joining network (B) including 202 *RdRp* sequences from the *Sarbecovirus* lineage isolated in bats, two sequences of SARS-CoV-2 and one sequence of SARS-CoV isolated in humans and eight sequences isolated in Malayan pangolins (*Manis javanica*). Colored circles correspond to distinct CoV sequences, circle size is proportional to the number of identical sequences in the data set. Small black circles represent median vectors (ancestral or unsampled intermediate sequences). Branch length is proportional to the number of mutational steps between haplotypes.

We used a Bayesian Stochastic Search Variable Selection (BSSVS) procedure^47^ to identify viral host switches (transmission over evolutionary time) between bat families and genera that occurred along the branches of the MCC annotated tree and calculated Bayesian Factor (BF) to estimate the significance of these switches (Fig. 4). We identified nine highly supported (BF > 10) inter-family host switches for α-CoVs and three for β-CoVs (Fig. 4A and 4B). These results are robust over a range of sample sizes, with seven of these nine switches for α-CoVs and the exact same three host switches for β-CoVs having strong BF support (BF > 10) when analyzing our random subset (Supplementary Tables 2 and 3). To quantify the magnitude of these host switches, we estimated the number of host switching events (Markov jumps)^48,49^ along the significant inter-family switches (Fig. 4C and 4D) and estimated the rate of inter-family host switching events per unit of time for each CoV genus. The rate of inter-family host switching events was five times higher in the evolutionary history of α- (0.010 host switches/unit time) than β-CoVs (0.002 host switches/unit time) in China. For α-CoVs, host switching events from the Rhinolophidae and the Miniopteridae were greater than from other bat families while rhinolophids were the highest donor family for β-CoVs. The Rhinolophidae and the Vespertilionidae for α-CoVs and the Hipposideridae for β-CoVs received the highest numbers of switching events (Fig. 4C and 4D). When using the random dataset, similar results were obtained for β-CoVs while rhinolophids were the highest donor family for α-CoVs (Supplementary Tables 4 and 5).

**Fig. 4.**
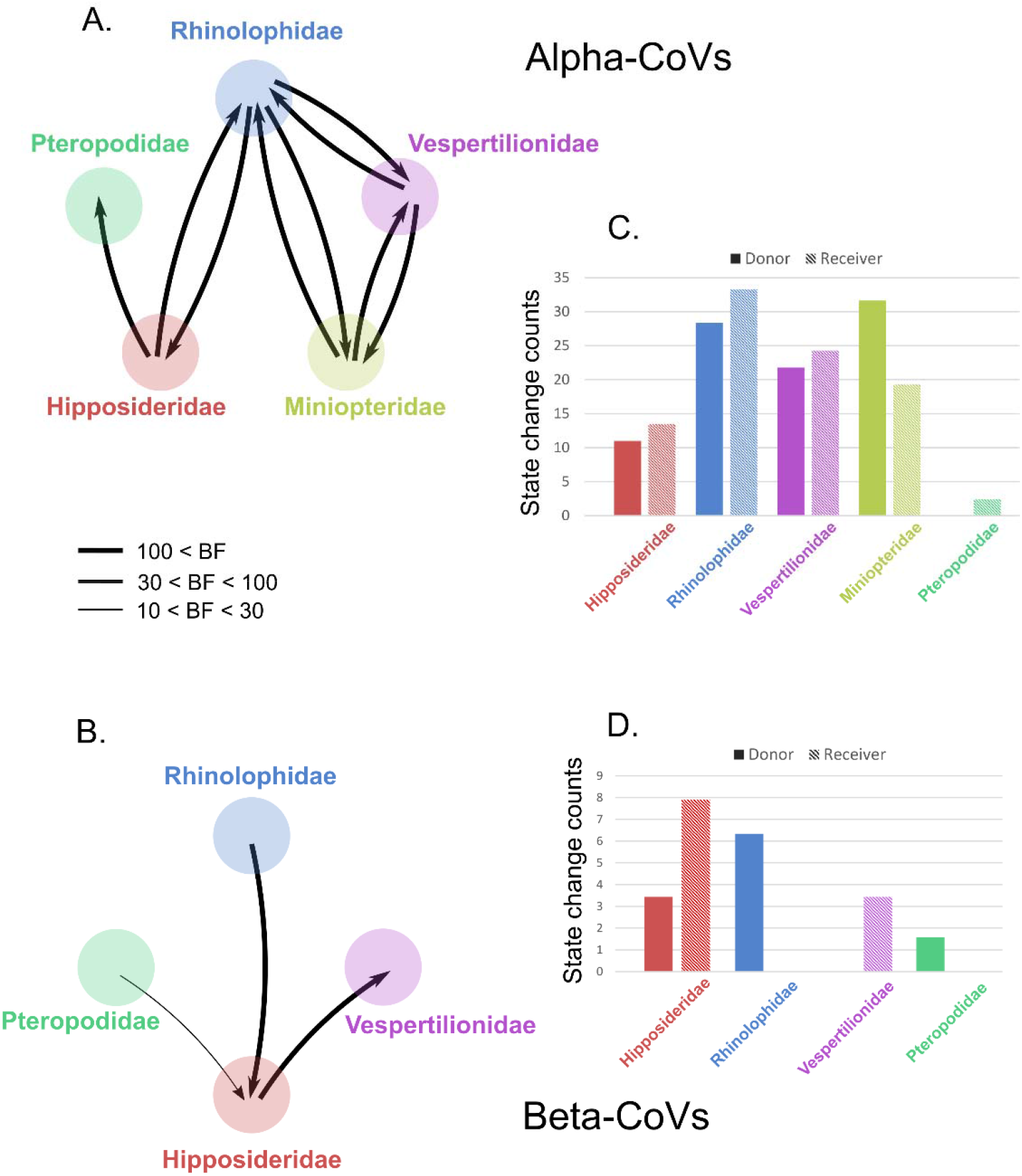
Inter-family host switches. Strongly supported host switches between bat families for alpha- (A) and beta-CoVs (B). Arrows indicate the direction of the switch; arrow thickness is proportional to the switch significance level, only host switches supported by strong Bayes factor (BF) > 10 are shown. Histograms of total number of host switching events (state changes counts using Markov jumps) from/to each bat family along the significant inter-family switches for alpha- (C) and beta-CoVs (D).

At the genus level, we identified 20 highly supported inter-genus host switches for α-CoVs, 17 of them were also highly significant using the random subset (Fig. 5A and Supplementary Table 6). Sixteen highly supported inter-genus switches were identified for β-CoVs (Fig. 5B). Similar results were obtained for the random β-CoV subset (Supplementary Table 7). Most of the significant cross-genus CoV switches for α-CoVs, 15 of 20 (75%), were between genera in different bat families, while this proportion was only 6 of 16 (37.5%) for β-CoVs. The estimated rate of inter-genus host switching events (Markov jumps) was similar for α- (0.014 host switches/unit time) and β-CoVs (0.014 host switches/unit time). For α-CoVs, *Rhinolophus* and *Miniopterus* were the greatest donor genera and *Rhinolophus* was the greatest receiver (Supplementary Table 8). For β-CoVs, *Rousettus* was the greatest donor and *Eonycteris* the greatest receiver genus (Supplementary Table 9).

**Fig. 5.**
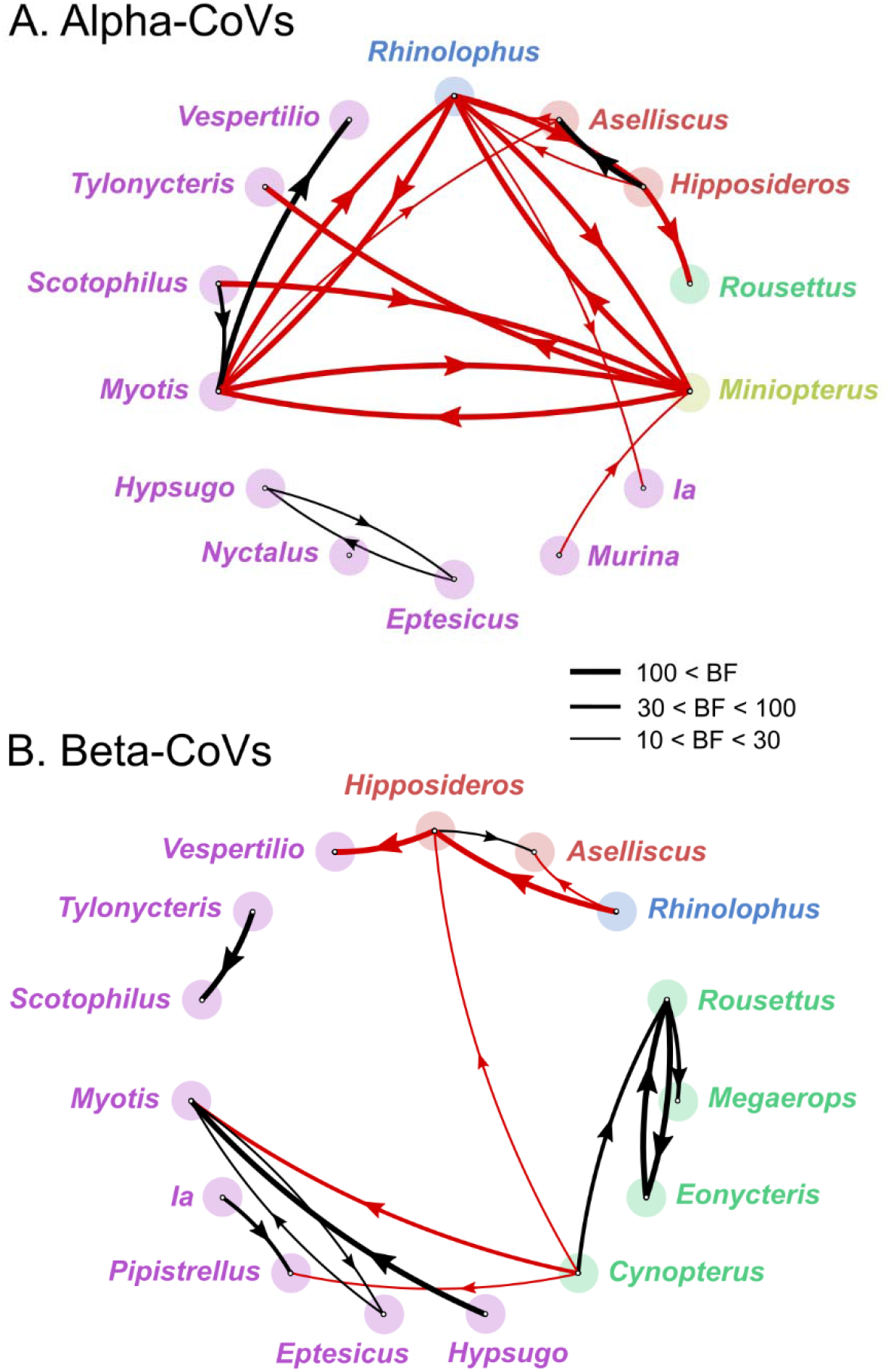
Inter-genus host switches. Strongly supported host switches between bat genera for alpha- (A) and beta-CoVs (B) and their significance level (Bayes factor, BF). Only host switches supported by strong BF values > 10 are shown. Line thickness is proportional to the switch significance level. Red lines correspond to host switches among bat genera belonging to different families, black lines correspond to host switches among bat genera from the same family. Arrows indicate the direction of the switch. Genus names are colored according to the family they belong to using the same colors as in Fig. 2 and 3.

### CoV spatiotemporal dispersal in China

We used our Bayesian discrete phylogeographic model with zoogeographic regions as character states to reconstruct the spatiotemporal dynamics of CoV dispersal in China. Eleven and seven highly significant (BF > 10) dispersal routes within China were identified for α- and β-CoVs, respectively (Fig. 6). Seven and five of these dispersal routes, respectively, remained significant when using our random subsets (Supplementary Tables 10 and 11). The *Rhinacovirus* lineage (L1) that includes HKU2- and SADS-CoV likely originated in the SO region while all other α-CoV lineages historically arose in SW China and spread to other regions before several dispersal events from SO and NO in all directions (Fig. 6A and Supplementary Fig. 12). A roughly similar pattern of α-CoV dispersal was obtained using the random subset (Supplementary Tables 10 and 12).

**Fig. 6.**
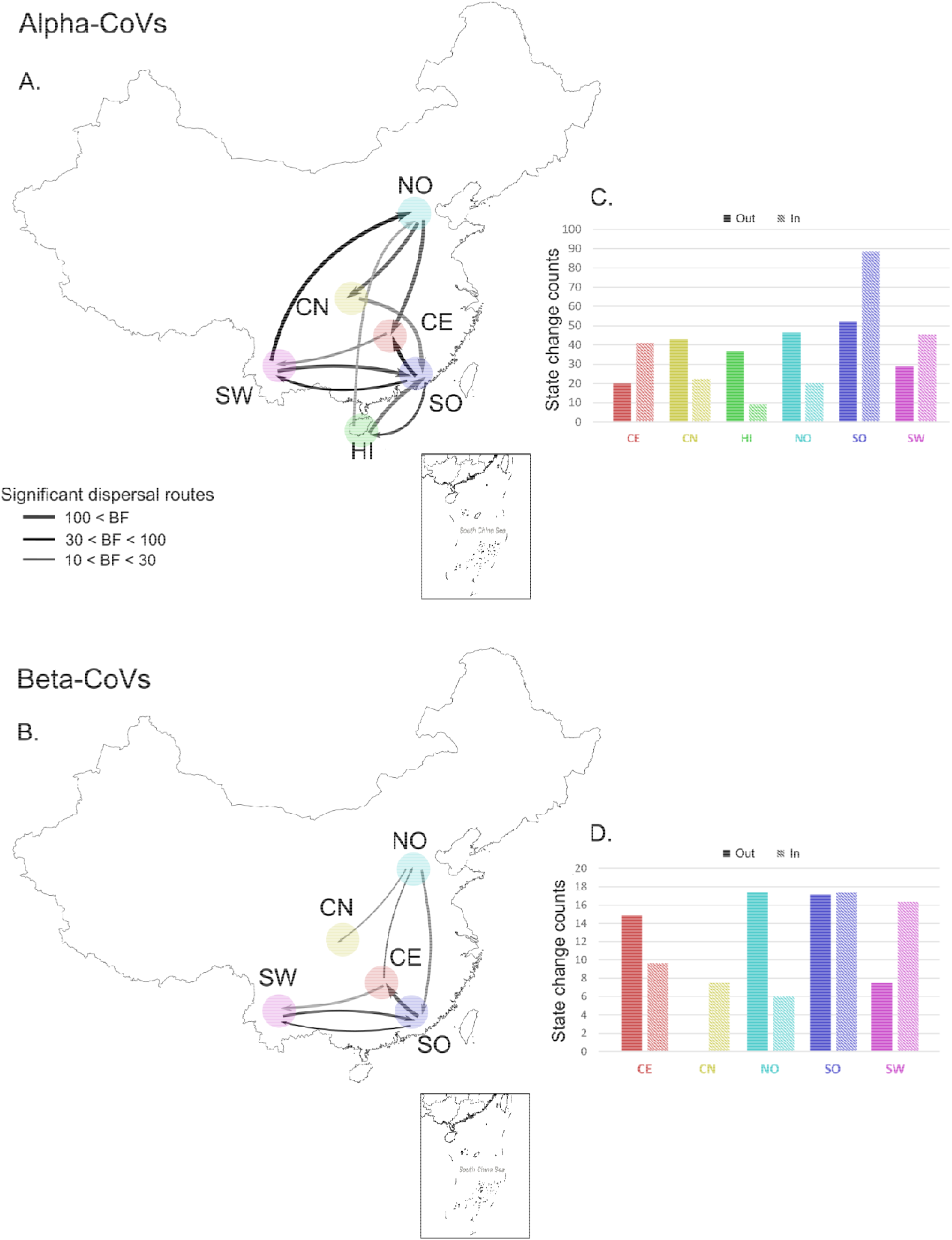
CoV spatiotemporal dispersal in China. Strongly supported dispersal routes (Bayes factor, BF > 10) over recent evolutionary history among China zoogeographic regions for alpha- (A) and beta-CoVs (B). Arrows indicate the direction of the dispersal route; arrow thickness is proportional to the dispersal route significance level. Darker arrow colors indicate older dispersal events. Histograms of total number of dispersal events (Markov jumps) from/to each region along the significant dispersal routes for alpha- (C) and beta-CoVs (D). NO, Northern region; CN, Central northern region; SW, South western region; CE, Central region; SO, Southern region; HI, Hainan island.

The oldest inferred dispersal movements for β-CoVs occurred among the SO and SW regions (Fig. 6B). The SO region was the likely origin of *Merbecovirus* (Lineage C, including HKU4- and HKU5-CoV) and *Sarbecovirus* subgenera (Lineage B, including HKU3- and SARSr-CoVs) while the *Nobecovirus* (lineage D, including HKU9-CoV) and *Hibecovirus* (lineage E) subgenera originated in SW China (Supplementary Fig. 12). Then several dispersal movements likely originated from SO and CE (Fig. 6B). More recent southward dispersal from NO was observed. Similar spatiotemporal dispersal patterns were observed using the random subset of β-CoVs (Supplementary Tables 11 and 13).

The estimated rate of migration events per unit of time along these significant dispersal routes was more than two times higher for α- (0.026 host switches/unit time) than β-CoVs (0.011 host switches/unit time) and SO was the region involved in the greatest total number of migration events for both α- and β-CoVs. SO had the highest number of outbound and inbound migration events for α-CoVs (Fig. 6C and Supplementary Table 12). For β-CoVs, the highest number of outbound migration events was estimated to be from NO and SO while SO and SW had the highest numbers of inbound migration events (Fig. 6D and Supplementary Table 13).

### Phylogenetic diversity

In order to identify the hotspots of CoV phylogenetic diversity in China and evaluate phylogenetic clustering of CoVs, we calculated the Mean Phylogenetic Distance (MPD) and the Mean Nearest Taxon Distance (MNTD) statistics^50^ and their standardized effect size (SES).

We found significant and negative SES MPD values, indicating significant phylogenetic clustering, within all bat families and genera for both α- and β-CoVs, except within the *Aselliscus* and *Tylonycteris* for α-CoVs (Fig. 7A and 7B). Negative and mostly significant SES MNTD values, reflecting phylogenetic structure closer to the tips, were also observed within most bat families and genera for α- and β-CoVs but we found non-significant positive SES MNTD value for vespertilionid bats, and particularly for those in the *Pipistrellus* genus, for β-CoVs (Fig. 7A and 7B). In general, we observed lower phylogenetic diversity for β- than α-CoVs within all bat families and most genera when looking at SES MPD, but the difference in the level of diversity between α- and β-CoVs is less important when looking at SES MNTD (Fig. 7). These results suggest stronger basal clustering (reflected by larger SES MPD values) for β-CoVs than α-CoVs, indicating stronger host structuring effect and phylogenetic conservatism for β-CoVs. Very similar results were obtained with the random subsets for both α- and β-CoVs (Supplementary Tables 14-21).

**Fig. 7.**
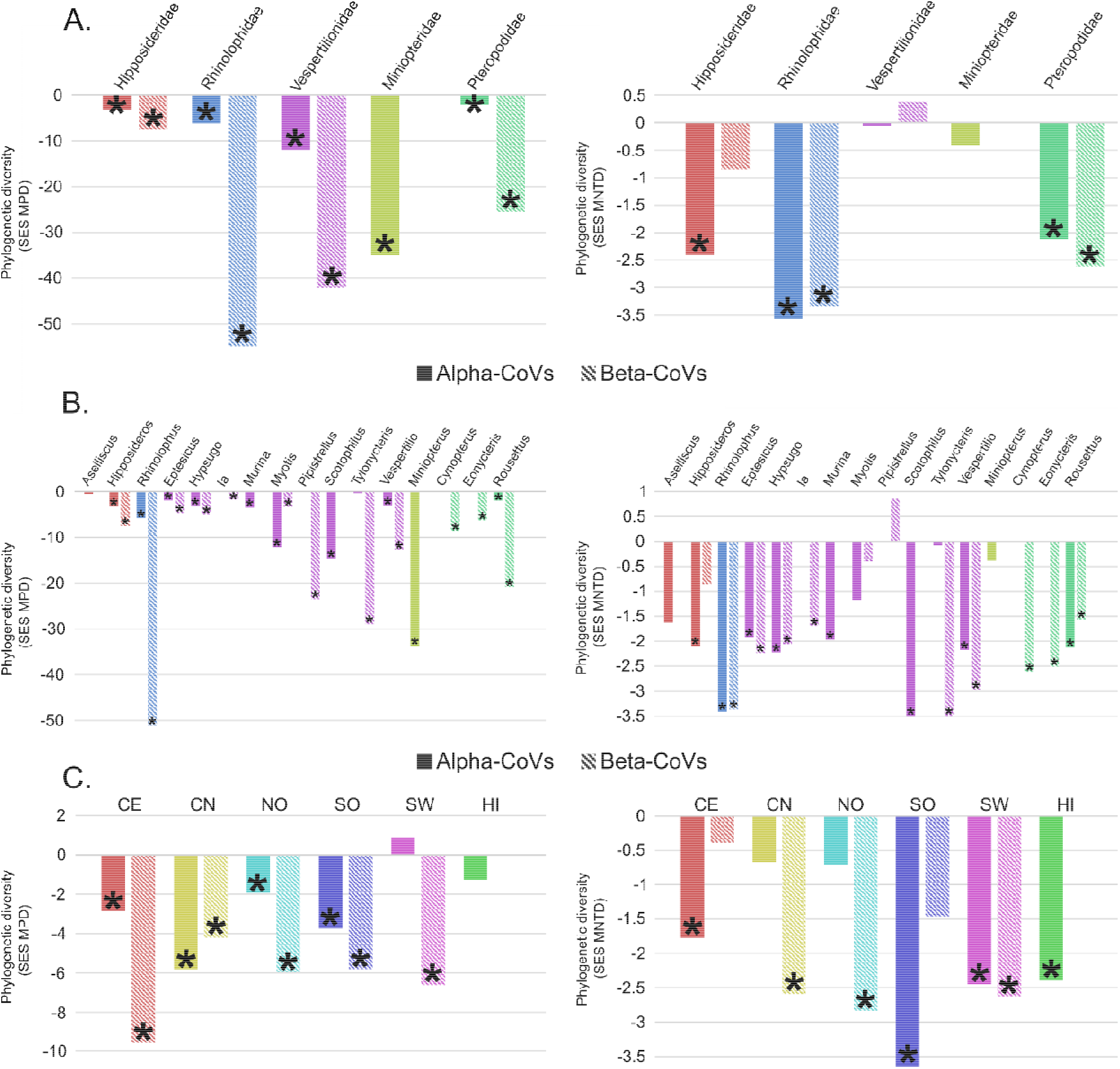
Phylogenetic diversity. Metrics of CoV phylogenetic diversity within each bat family (A), genus (B) and zoogeographic regions (C): standardized effect size of Mean Phylogenetic Distance (SES MPD), on the left panels; and standardized effect size of Mean Nearest Taxon Distance (SES MNTD), on the right panels. One-tailed p-values (quantiles) were calculated after randomly reshuffling tip labels 1000 times along the entire phylogeny. Values departing significantly from the null model (p-value < 0.05) are indicated with an asterisk,all exact p-values are available in Supplementary Tables 14-27. NO, Northern region; CN, Central northern region; SW, South western region; CE, Central region; SO, Southern region; HI, Hainan island.

We found negative and mostly significant values of MPD and MNTD (Fig. 7C and Supplementary Tables 22-25) indicating significant phylogenetic clustering of CoV lineages in bat communities within the same zoogeographic region. However, SES MPD values for α-CoVs in SW were positive (significant for the random subset) indicating a greater evolutionary diversity of CoVs in that region than others (Fig. 7 and Supplementary Tables 22-25). We used a linear regression analysis to assess the relationship between CoV phylogenetic diversity and bat species richness in China and determine if bat richness is a significant predictor of bat-CoV diversity and evolution. α-CoV phylogenetic diversity (MPD) was not significantly correlated to total bat species richness or sampled bat species richness in zoogeographic regions or provinces (Supplementary Table 26). Non-significant correlations between bat species richness and β-CoV phylogenetic diversity were also observed at the zoogeographic region level (Supplementary Table 27). However, a significant correlation was observed between sampled bat species richness and β-CoV phylogenetic diversity at the province level (Supplementary Table 27). Similar results were obtained when using the random subsets (Supplementary Tables 26 and 27). These findings suggest that bat host diversity is not the main driver of CoV diversity in China and that other ecological or biogeographic factors may influence this diversity. We observed higher CoV diversity than expected in several southern or central provinces (Hainan, Guangxi, Hunan) given their underlying total or sampled bat diversity (Supplementary Fig. 13 and 14).

We also assessed patterns of CoV phylogenetic turnover/differentiation among Chinese zoogeographic regions and bat host families by measuring the inter-region and inter-host values of MPD (equivalent to a measure of phylogenetic β-diversity) and their SES. We found positive inter-family SES MPD values, except between Pteropodidae and Hipposideridae for α-CoVs and between Rhinolophidae and Hipposideridae for β-CoVs (Fig. 8A and 8B and Supplementary Tables 28 and 29), suggesting higher phylogenetic differentiation of CoVs among most bat families than among random communities. Our phylo-ordination based on inter-family MPD values indicated that α-CoVs from vespertilionids and miniopterids, and from hipposiderids and pteropodids; as well as β-CoVs from rhinolophids and hipposiderids are phylogenetically closely related (Fig. 8A and 8B). We also observed strong phylogenetic turnover between α-CoV strains from rhinolophids and from miniopterids and all other bat families, and between β-CoV strains from vespertilionids and all other bat families (Supplementary Tables 28 and 29). Phylo-ordination among bat genera based on inter-genus MPD confirmed these results and indicated that CoV strains from genera belonging to the same bat family were mostly more closely related to each other than to genera from other families (Fig. 8C and 8D and Supplementary Tables 30 and 31).

**Fig. 8.**
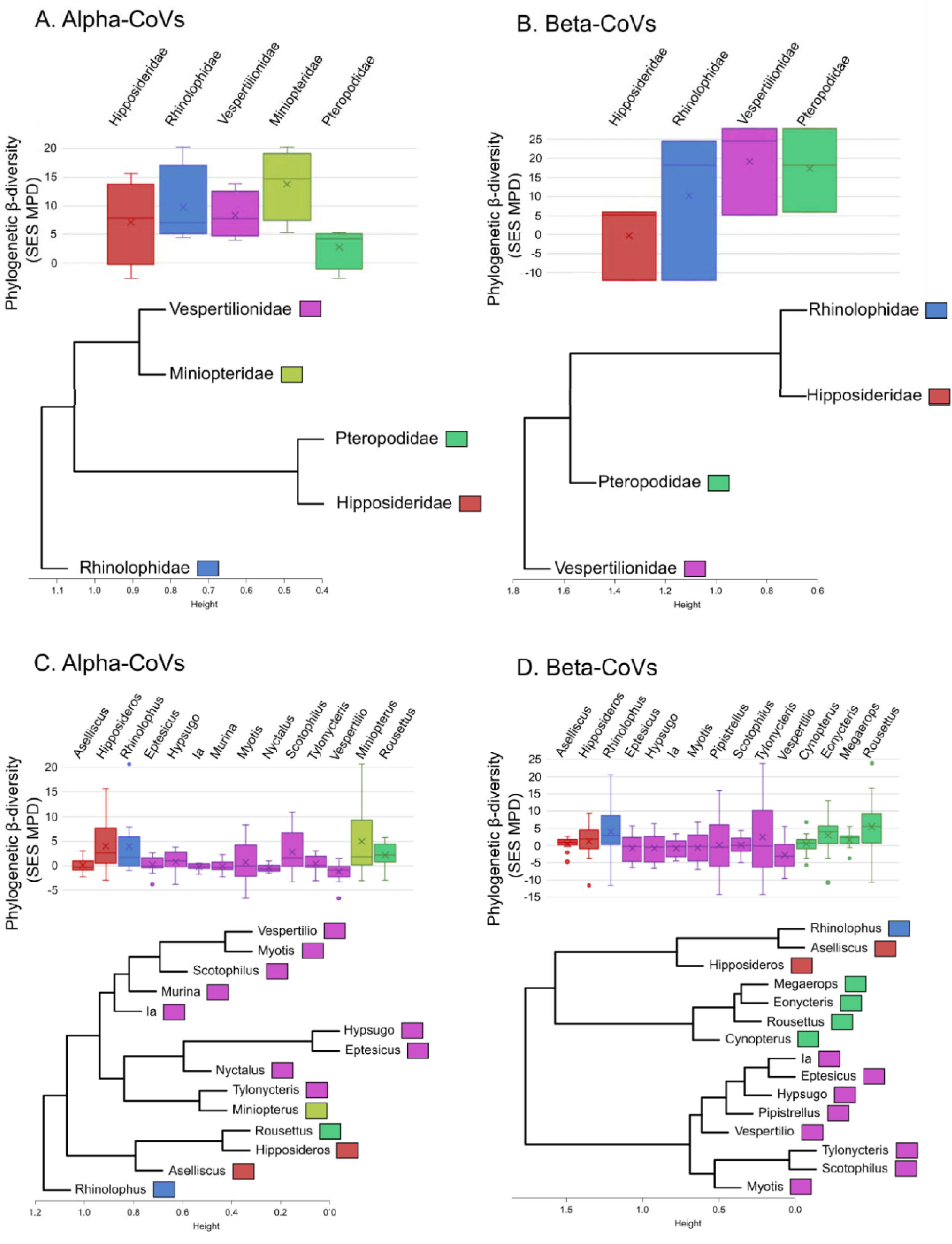
Phylogenetic diversity. Standardized effect size of Mean Phylogenetic Distance (SES MPD) and phylogenetic ordination among bat host families (A, B) and genera (C, D) for alpha- and beta-CoVs. Boxplots for each host family and genus show the mean (cross), median (dark line within the box), interquartile range (box), 95% confidence interval (whisker bars), and outliers (dots), calculated from all pairwise comparisons between bat families (n=10 for alpha-CoVs and n=6 for beta-CoVs) and genera (n=91 for alpha-CoVs and n=105 for beta-CoVs).

We observed high and positive inter-region SES MPD values between SW/HI and all other regions, suggesting that these two regions host higher endemic diversity (Fig. 9 and Supplementary Tables 32 and 31). Negative inter-region SES MPD values suggested that the phylogenetic turnover among other regions was less important than expected among random communities. Our phylo-ordination among zoogeographic regions also reflected the high phylogenetic turnover and deep evolutionary distinctiveness of both α- and β-CoVs from SW and HI regions (Fig. 9 and Supplementary Tables 32 and 33). Similar results were obtained using the random subset (Supplementary Tables 32 and 33).

**Fig. 9.**
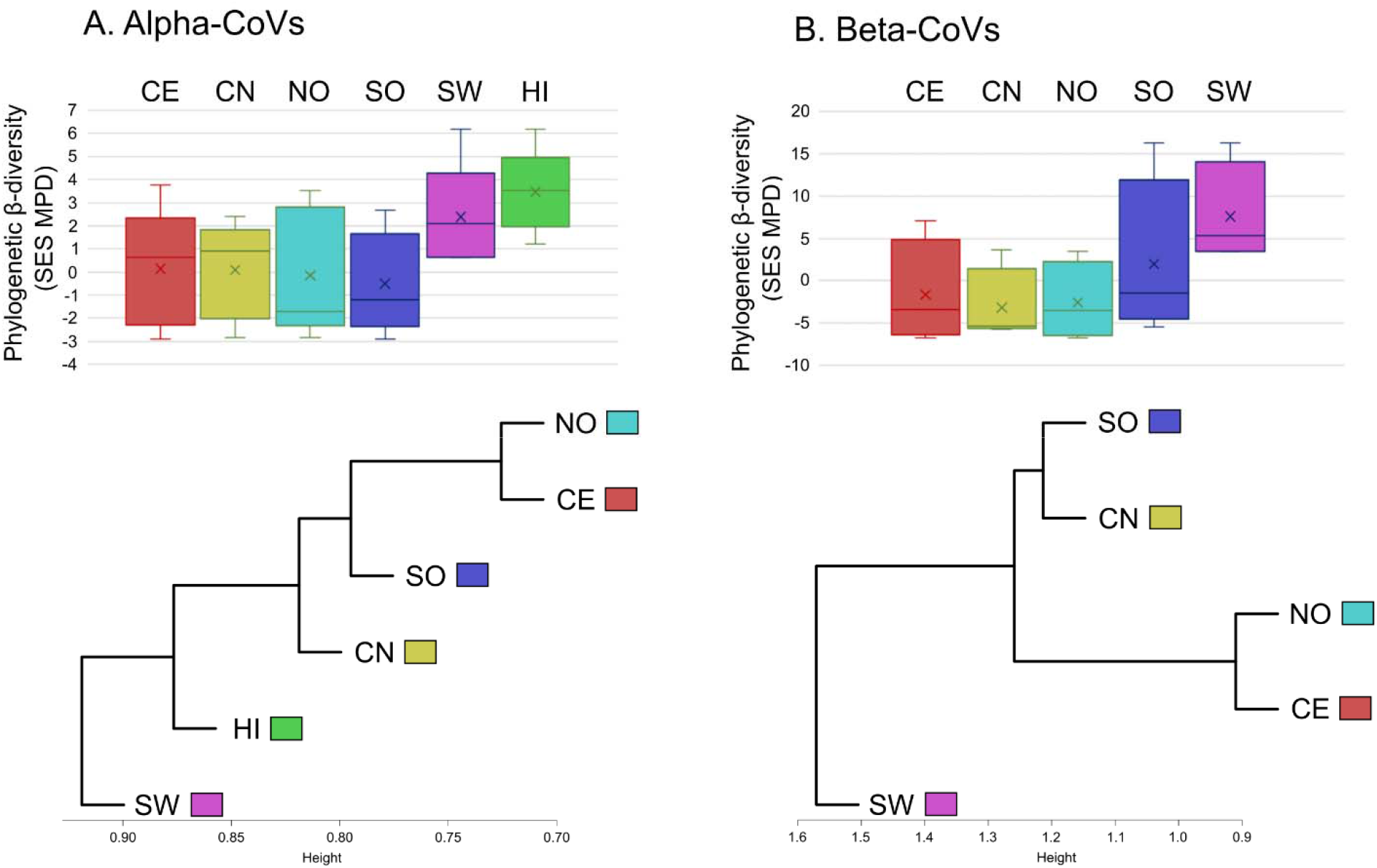
Phylogenetic diversity. Standardized effect size of Mean Phylogenetic Distance, SES MPD) and phylogenetic ordination among zoogeographic regions for alpha- (A) and beta-CoVs (B). Boxplots for each region show the mean (cross), median (dark line within the box), interquartile range (box), 95% confidence interval (whisker bars), and outliers (dots), calculated from all pairwise comparisons between regions (n=15 for alpha-CoVs and n=10 for beta-CoVs). NO, Northern region; CN, Central northern region; SW, South western region; CE, Central region; SO, Southern region; HI, Hainan island.

### Mantel tests

Mantel tests revealed a positive and significant correlation between CoV genetic differentiation (*F*_ST_) and geographic distance matrices, both with and without provinces including fewer than four viral sequences, for α- (*r* = 0.25, p = 0.0097; *r* = 0.32, p = 0.0196; respectively) and β-CoVs (*r* = 0.22, p = 0.0095; *r* = 0.23, p = 0.0336; respectively). We also detected a positive and highly significant correlation between CoV genetic differentiation (*F*_ST_) and their host phylogenetic distance matrices, both with and without genera including fewer than four viral sequences, for β-CoVs (*r* = 0.41, p = 0; *r* = 0.39, p = 0.0012; respectively) but not for α-CoVs (*r* = −0.13, p = 0.8413; *r* = 0.02, p = 0.5019; respectively).

## Discussion

Our phylogenetic analysis shows a high diversity of CoVs from bats sampled in China, with most bat genera included in this study (10/16) infected by both α- and β-CoVs. In our phylogenetic analysis that includes all known bat-CoVs from China, we find that SARS-CoV-2 is likely derived from a clade of viruses originating in horseshoe bats (*Rhinolophus* spp.). The geographic location of this origin appears to be Yunnan province. However, it is important to note that: 1) our study collected and analyzed samples solely from China; 2) many sampling sites were close to the borders of Myanmar and Lao PDR; and 3) most of the bats sampled in Yunnan also occur in these countries, including *R. affinis* and *R. malayanus*, the species harboring the CoVs with highest *RdRp* sequence identity to SARS-CoV-2^20,21^. For these reasons, we cannot rule out an origin for the clade of viruses that are progenitors of SARS-CoV-2 that is outside China, and within Myanmar, Lao PDR, Vietnam or another Southeast Asian country. Additionally, our analysis shows that the virus RmYN02 from *R. malayanus*, which is characterized by the insertion of multiple amino acids at the junction site of the S1 and S2 subunits of the Spike (S) protein, belongs to the same clade as both RaTG13 and SARS-CoV-2, providing further support for the natural origin of SARS-CoV-2 in *Rhinolophus* spp. bats in the region^20,21^. Finally, while our analysis shows that the RdRp sequences of coronaviruses from the Malayan pangolin are closely related to SARS-CoV-2 RdRp, analysis of full genomes of these viruses suggest that these terrestrial mammals are less likely to be the origin of SARS-CoV-2 than *Rhinolophus* spp. bats^22,23^.

This analysis also demonstrates that a significant amount of cross-species transmission has occurred among bat hosts over evolutionary time. Our Bayesian phylogeographic inference and analysis of host switching showed varying levels of viral connectivity among bat hosts and allowed us to identify significant host transitions that appear to have occurred during bat-CoV evolution in China.

We found that bats in the family Rhinolophidae (horseshoe bats) played a key role in the evolution and cross-species transmission history of α-CoVs. The family Rhinolophidae and the genus *Rhinolophus* were involved in more inter-family and inter-genus highly significant host switching of α-CoVs than any other family or genus. They were the greatest receivers of α-CoV host switching events and second greatest donors after Miniopteridae/*Miniopterus*. The Rhinolophidae, together with the Hipposideridae, also played an important role in the evolution of β-CoVs, being at the origin of most inter-family host switching events. Chinese horseshoe bats are characterized by a distinct and evolutionary divergent α- CoV diversity, while their β-CoV diversity is similar to that found in the Hipposideridae. The Rhinolophidae comprises a single genus, *Rhinolophus*, and is the most speciose bat family after the Vespertilionidae in China^51^, with 20 known species, just under a third of global *Rhinolophus* diversity, mostly in Southern China^35^. This family likely originated in Asia^52,53^, but some studies suggest an African origin^54,55^. Rhinolophid fossils from the middle Eocene (38 - 47.8 Mya) have been found in China, suggesting a westward dispersal of the group from eastern Asia to Europe^56^. The ancient likely origin of the Rhinolophidae in Asia and China in particular may explain the central role they played in the evolution and diversification of bat-CoVs in this region, including SARSr-CoVs, MERS-cluster CoVs, and SADSr-CoVs, which contain important human and livestock pathogens. Horseshoe bats are known to share roosts with genera from all other bat families in this study^57^, which may also favor CoV cross-species transmission from and to rhinolophids^34^. A global meta-analysis showing higher rates of viral sharing among co-roosting cave bats supports this finding^58^.

Vespertilionid and miniopterid bats (largely within the *Myotis* and *Miniopterus* genera) also appear to have been involved in several significant host switches during α-CoV evolution. However, no significant transition from vespertilionid bats was identified for β-CoVs and these bats exhibit a divergent β-CoV diversity compared to other bat families. Vespertilionid and miniopterid bats are characterized by strong basal phylogenetic clustering but high recent CoV diversification rates, indicating a more rapid evolutionary radiation of CoVs in these bat hosts. At the genus level, similar findings were observed for the genera *Myotis, Pipistrellus* and *Miniopterus*.

A significant correlation between geographic distance and genetic differentiation of both α- and β-CoVs has been detected, even if only a relatively small proportion of the variance is explained by geographic distance. We also revealed a significant effect of host phylogeny on β-CoV evolution while it had a minimal effect on α-CoV diversity. Contrary to the α-CoV phylogeny, the basal phylogenetic structure of β-CoVs mirrored the phylogeny of their bat hosts, with a clear distinction between the Yangochiroptera, encompassing the Vespertilionidae and Miniopteridae, and the Yinpterochiroptera, which includes the megabat family Pteropodidae and the microbat families Rhinolophidae and Hipposideridae, as evidenced in recent bat phylogenies^52,59^. These findings suggest a profound co-macroevolutionary process between β-CoVs and their bat hosts, even if host switches also occurred throughout their evolution as our study showed. The phylogenetic structure of α-CoVs, with numerous and closely related lineages identified in the Vespertilionidae and Miniopteridae, contrasts with the β-CoV macroevolutionary pattern and suggests α-CoVs have undergone an adaptive radiation in these two Yangochiroptera families. Our BSSVS procedure and Markov jump estimates revealed higher connectivity, both qualitatively and quantitatively, among bat families and genera in the α-CoV cross-species transmission history. Larger numbers of highly significant host transitions and higher rates of switching events along these pathways were inferred for α- than β-CoVs, especially at the host family level. These findings suggest that α-CoVs are able to switch hosts more frequently and between more distantly related taxa, and that phylogenetic distance among hosts represents a higher constraint on host switches for β- than α-CoVs. This is supported by more frequent dispersal events in the evolution of α- than β-CoVs in China.

Variation in the extent of host jumps between α and β-CoVs within the same hosts in the same environment may be due to virus-specific factors such as differences in receptor usage between α- and β-CoVs^60–62^. Coronaviruses use a large diversity of receptors, and their entry into host cells is mediated by the spike protein with an ectodomain consisting of a receptor-binding subunit S1 and a membrane-fusion subunit S2^63^. However, despite differences in the core structure of their S1 receptor binding domains (RBD), several α- and β-CoV species are able to recognize and bind to the same host receptors^64^. Other factors such as mutation rate, recombination potential, or replication rate might also be involved in differences in host switching potential between α- and β-CoVs. A better understanding of receptor usage and other biological characteristics of these bat-CoVs may help predict their cross-species transmission and zoonotic potential.

We also found that some bat genera were infected by a single CoV genus: *Miniopterus* (Miniopteridae) and *Murina* (Vespertilionidae) carried only α-CoVs, while *Cynopterus, Eonycteris, Megaerops* (Pteropodidae) and *Pipistrellus* (Vespertilionidae) hosted only β-CoVs. This was found despite using the same conserved pan-CoV PCR assays for all specimens screened and it can’t be explained by differences in sampling effort for these genera (Supplementary Table 1): for example, >250 α-CoV sequences but no β-CoV were discovered in *Miniopterus* bats in China during our recent fieldwork. These migratory bats, which seem to have played a key role in the evolution of α-CoVs, share roosts with several other bat genera hosting β-CoVs in China^57^, suggesting high likelihood of being exposed to β-CoVs. Biological or ecological properties of miniopterid bats may explain this observation and clearly warrant further investigation.

Our Bayesian ancestral reconstructions revealed the importance of South western and Southern China as centers of diversification for both α- and β-CoVs. These two regions are hotspots of CoV phylogenetic diversity, harboring evolutionarily old and phylogenetically diverse lineages of α- and β-CoVs. South western China acted as a refugium during Quaternary glaciation for numerous plant and animal species including several bat species, such as *Rhinolophus affinis*^65^, *Rhinolophus sinicus*^66^, *Myotis davidii*^67^, and *Cynopterus sphinx*^68^. The stable and long-term persistence of bats and other mammals throughout the Quaternary may explain the deep macroevolutionary diversity of bat-CoVs in these regions^69^. Several highly significant and ancient CoV dispersal routes from these two regions have been identified in this study. Other viruses, such as the Avian Influenza A viruses H5N6, H7N9 and H5N1, also likely originated in South western and Southern Chinese regions^70,71^.

Our findings suggest that bat host diversity is not the main driver of CoV diversity in China and that other ecological or biogeographic factors may influence this diversity. Overall, there were no significant correlations between CoV phylogenetic diversity and bat species diversity (total or sampled) for each province or biogeographic region, apart from a weak correlation between β-CoV phylogenetic diversity and the number of bat species sampled at the province level. Yet, we observed higher than expected phylogenetic diversity in several southern provinces (Hainan, Guangxi, Hunan). These results and main conclusions are consistent and robust even when we account for geographic biases in sampling effort by analyzing random subsets of the data.

Despite being the most exhaustive study of bat-CoVs in China, this study had several limitations that must be taken into consideration when interpreting our results. First, only partial *RdRp* sequences were generated in this study and used in our phylogenetic analysis as the non-invasive samples (rectal swabs/feces) collected in this study prevented us from generating longer sequences in many cases. The *RdRp* gene is a suitable marker for this kind of study as it reflects vertical ancestry and is less prone to recombination than other regions of the CoV genome such as the spike protein gene^16,72^. While using long sequences is always preferable, our phylogenetic trees are well supported and their topology consistent with trees obtained using longer sequences or whole genomes^30,73^. Second, most sequences in this study were obtained by consensus PCR using primers targeting highly conserved regions. Even if this broadly reactive PCR assay designed to detect widely variant CoVs has proven its ability to detect a large diversity of CoVs in a wide diversity of bats and mammals^30,74–77^, we may not rule out that some bat-CoV variants remained undetected. Using deep sequencing techniques would allow to detect this unknown and highly divergent diversity.

In this study, we identified the host taxa and geographic regions that together define hotspots of CoV phylogenetic diversity and centers of diversification in China. These findings may provide a strategy for targeted discovery of bat-borne CoVs of zoonotic or livestock infection potential, and for early detection of bat-CoV outbreaks in livestock and people, as proposed elsewhere^78^. Our results suggest that future sampling and viral discovery should target two hotspots of CoV diversification in Southern and South western China in particular, as well as neighboring countries where similar bat species live. These regions are characterized by a subtropical to tropical climate; dense, growing and rapidly urbanizing populations of people; a high degree of poultry and livestock production; and other factors which may promote cross-species transmission and disease emergence^78–80^. Additionally, faster rates of evolution in the tropics have been described for other RNA viruses which could favor cross-species transmission of RNA viruses in these regions^81^. Both SARS-CoV and SADS-CoV emerged in this region, and several bat SARSr-CoVs with high zoonotic potential have recently been reported from there, although the dynamics of their circulation in wild bat populations remain poorly understood^16,61^. Importantly, the closest known relative of SARS-CoV-2, a SARS-related virus, was found in a *Rhinolophus* sp. bat in this region^20^, although it is important to note that our survey was limited to China, and that the bat hosts of this virus also occur in nearby Myanmar and Lao PDR. The significant public health and food security implications of these outbreaks reinforces the need for enhanced, targeted sampling and discovery of novel CoVs. Because intensive sampling has not, to our knowledge, been undertaken in countries bordering southern China, these surveys should be extended to include Myanmar, Lao PDR, and Vietnam, and perhaps across southeast Asia. Our finding that *Rhinolophus* spp. are most likely to be involved in host-switching events makes them a key target for future longitudinal surveillance programs, but surveillance targeted the genera *Hipposideros* and *Aselliscus* may also be fruitful as they share numerous β-CoVs with *Rhinolophus* bats.

In the aftermath of the SARS-CoV and MERS-CoV outbreaks, β-CoVs have been the main focus of bat-CoV studies in China, Africa, and Europe^17,32,36,61,82^. However, we have shown that α-CoVs have a higher propensity to switch host within their natural bat reservoirs, and therefore also have a high cross-species transmission potential and risk of spillover. This is exemplified by the recent emergence of SADS-CoV in pigs in Guangdong province^17^. Two human α-CoVs, NL63 and 229E, also likely originated in bats^27,28^, reminding us that past spillover events from bat species can readily be established in the human population. Future work discovering and characterizing the biological properties of bat α-CoVs may therefore be of potential value for public and livestock health. Our study, and recent analysis of viral discovery rates^83^, suggest that a substantially wider sampling and discovery net will be required to capture the complete diversity of coronaviruses in their natural hosts and assess their potential for cross-species transmission. The bat genera *Rhinolophus*, *Hipposideros*, *Myotis* and *Miniopterus*, all involved in numerous naturally-occurring host switches throughout α-CoV evolution, should be a particular target for α-CoV discovery in China and across southeast Asia, with *in vitro* and experimental characterization to better understand their potential to infect people or livestock and cause disease.

## Methods

### Bat sampling

Bat oral and rectal swabs and fecal pellets were collected from 2010 to 2015 in numerous Chinese provinces (Anhui, Beijing, Guangdong, Guangxi, Guizhou, Hainan, Henan, Hubei, Hunan, Jiangxi, Macau, Shanxi, Sichuan, Yunnan, and Zhejiang). Fecal pellets were collected from tarps placed below bat colonies. Bats were captured using mist nets at their roost site or feeding areas. Each captured bat was stored into a cotton bag, all sampling was non-lethal and bats were released at the site of capture immediately after sample collection. A wing punch was also collected for barcoding purpose. Bat-handling methods were approved by Tufts University IACUC committee (proposal #G2017-32) and Wuhan Institute of Virology Chinese Academy of Sciences IACUC committee (proposal WIVA05201705). Samples were stored in viral transport medium at −80°C directly after collection.

### RNA extraction and PCR screening

RNA was extracted from 200 μl swab rectal samples or fecal pellets with the High Pure Viral RNA Kit (Roche) following the manufacturer’s instructions. RNA was eluted in 50 μl elution buffer and stored at −80°C. A one-step hemi-nested RT-PCR (Invitrogen) was used to detect coronavirus RNA using a set of primers targeting a 440-nt fragment of the *RdRp* gene and optimized for bat-CoV detection (CoV-FWD3: GGTTGGGAYTAYCCHAARTGTGA; CoV-RVS3: CCATCATCASWYRAATCATCATA; CoV-FWD4/Bat: GAYTAYCCHAARTGTGAYAGAGC)^84^. For the first round PCR, the amplification was performed as follows: 50°C for 30 min, 94°C for 2 min, followed by 40 cycles consisting of 94°C for 20 sec, 50°C for 30 sec, 68°C for 30 sec, and a final extension step at 68°C for 5 min. For the second round PCR, the amplification was performed as follows: 94°C for 2 min followed by 40 cycles consisting of 94°C for 20 sec, 59°C for 30 sec, 72°C for 30 sec, and a final extension step at 72°C for 7 min. PCR products were gel purified and sequenced with an ABI Prism 3730 DNA analyzer (Applied Biosystems, USA). PCR products with low concentration or bad sequencing quality were cloned into pGEM-T Easy Vector (Promega) for sequencing. Positive results detected in bat genera that were not known to harbor a specific CoV lineage previously were repeated a second time (PCR + sequencing) as a confirmation. Species identifications from the field were also confirmed and re-confirmed by cytochrome (cytb) DNA barcoding using DNA extracted from the feces or swabs^85^. Only viral detection and barcoding results confirmed at least twice were included in this study.

### Sequence data

We also added bat-CoV *RdRp* sequences from China available in GenBank to our dataset. All sequences for which sampling year and host or sampling location information was available either in GenBank metadata or in the original publication were included (as of March 15, 2018). Our final datasets include 630 sequences generated for this study and 616 sequences from GenBank or GISAID (list of GenBank and GISAID accession numbers available in Supplementary Note 1, and Supplementary Tables 34 and 35). Nucleotide sequences were aligned using MUSCLE and trimmed to 360 base pair length to reduce the proportion of missing data in the alignments. All phylogenetic analyses were performed on both the complete data and random subset, and for α- and β-CoVs separately.

### Defining zoogeographic regions in China

Hierachical clustering was used to define zoogeographic regions within China by clustering provinces with similar mammalian diversity^45^. Hierarchical cluster analysis classifies several objects into small groups based on similarities between them. To do this, we created a presence/absence matrix of all extant terrestrial mammals present in China using data from the IUCN spatial database^86^ and generated a cluster dendrogram using the function *hclust* with average method of the R package stats. Hong Kong and Macau were included within the neighboring Guangdong province. We then visually identified geographically contiguous clusters of provinces for which CoV sequences are available (Fig. 1 and Supplementary Fig. 1).

We identified six zoogeographic regions within China based on the similarity of the mammal community in these provinces: South western region (SW; Yunnan province), Northern region (NO; Xizang, Gansu, Jilin, Anhui, Henan, Shandong, Shaanxi, Hebei and Shanxi provinces and Beijing municipality), Central northern region (CN; Sichuan and Hubei provinces), Central region (CE; Guangxi, Guizhou, Hunan, Jiangxi and Zhejiang provinces), Southern region (SO; Guangdong and Fujian provinces, Hong Kong, Macau and Taiwan), and Hainan island (HI). Hunan and Jiangxi, clustering with the SO provinces in our dendrogram, were included within the central region to create a geographically contiguous Central cluster (Supplementary Fig. 1). These six zoogeographic regions are very similar to the biogeographic regions traditionally recognized in China^87^. The three β-CoV sequences from HI were included in the SO region to avoid creating a cluster with a very small number of sequences.

### Model selection and phylogenetic analysis

Bayesian phylogenetic analysis were performed in BEAST 1.8.4^46^. Sampling years were used as tip dates. Preliminary analysis were run to select the best fitting combination of substitution models (HKY/GTR), codon partition scheme, molecular clock (strict/lognormal uncorrelated relaxed clock) and coalescent models (constant population size/exponential growth/GMRF Bayesian Skyride). Model combinations were compared and the best fitting model was selected using a modified Akaike information criterion (AICM) implemented in Tracer 1.6^88^. We also used TEMPEST^89^ to assess the temporal structure within our α- and β-CoV datasets. TEMPEST showed that both datasets did not contain sufficient temporal information to accurately estimate substitution rates or time to the most recent common ancestor (TMRCA). Therefore we used a fixed substitution rate of 1.0 for all our BEAST analysis.

All subsequent BEAST analysis were performed under the best fitting model including a HKY substitution model with two codons partitions ((1+2), 3), a strict molecular clock and a constant population size coalescent model. Each analysis was run for 2.5 × 10^8^ generations, with sampling every 2 × 10^4^ steps. All BEAST computations were performed on the CIPRES Science Getaway Portal^90^. Convergence of the chain was assessed in Tracer so that the effective sample size (ESS) of all parameters was > 200 after removing at least 10% of the chain as burn-in.

### Ancestral state reconstruction and transition rates

A Bayesian discrete phylogeographic approach implemented in BEAST 1.8.4 was used to reconstruct the ancestral state of each node in the phylogenetic tree for three discrete traits: host family, host genus and zoogeographic region. An asymmetric trait substitution model was applied. These analyses were performed for each trait on the complete dataset and random subsets. Maximum clade credibility (MCC) tree annotated with discrete traits were generated in TreeAnnotator and visualized using the software SpreaD3^91^.

For each analysis, a Bayesian stochastic search variable selection (BSSVS) was applied to estimate the significance of pairwise switches between trait states using Bayesian Factor (BF) as a measure of statistical significance^47^. BF were computed in SpreaD3. BF support was interpreted according to Jeffreys 1961^92^ (BF > 3: substantial support, BF > 10: strong support, BF > 30: very strong support, BF > 100: decisive support) and only strongly supported transitions were presented in most figures, following a strategy used in other studies^93,94^. We also estimated the count of state switching events (Markov jumps)^48,49^ along the branches of the phylogenetic tree globally (for the three discrete traits) and for each strongly supported (BF > 10) transition between character states (for bat families and ecoregions only). Convergence of the MCMC runs was confirmed using Tracer. The rate of state switching events per unit of time was estimated for each CoV genus by dividing the total estimated number of state switching events by the total branch length of the MCC tree.

To assess the phylogenetic relationships among SARS-CoV-2 and other CoVs from the *Sarbecovirus* subgenus, we also reconstructed a MCC tree in BEAST 1.8.4 and median-joining network in Network 10.0^95^ including all *Sarbecovirus* sequences, two sequences of SARS-CoV-2 isolated in humans (GenBank accession numbers: MN908947 and MN975262), one sequence of SARS-CoV (GenBank accession number: NC_004718), eight sequences from Malayan pangolins (*Manis javanica*) (GISAID accession numbers: EPI_ISL_410538-410544, EPI_ISL_410721) and one from *Rhinolophus malayanus* (GISAID accession number: EPI_ISL_412977).

### Phylogenetic diversity

The Mean Phylogenetic Distance (MPD) and the Mean Nearest Taxon Distance (MNTD) statistics^50^ and their standardized effect size (SES) were calculated for each zoogeographic region, bat family and genus using the R package picante^96^. MPD measures the mean phylogenetic distance among all pairs of CoVs within a host or a region. It reflects phylogenetic structuring across the whole phylogenetic tree and assesses the overall divergence of CoV lineages in a community. MNTD is the mean distance between each CoV and its nearest phylogenetic neighbor in a host or region, and therefore it reflects the phylogenetic structuring closer to the tips and shows how locally clustered taxa are. SES MPD and SES MNTD values correspond to the difference between the phylogenetic distances in the observed communities versus null communities. Low and negative SES values denote phylogenetic clustering, high and positive values indicate phylogenetic over-dispersion while values close to 0 show random dispersion. The SES values were calculated by building null communities by randomly reshuffling tip labels 1000 times along the entire phylogeny. Phylogenetic diversity computations were performed on both the complete dataset and random subset for each trait. A linear regression analysis was performed in R to assess the correlation between CoV phylogenetic diversity (MPD) and bat species richness in China. Total species richness per province or region was estimated using data from the IUCN spatial database while sampled species richness corresponds to the number of bat species sampled and tested for CoV per province or region in our datasets.

The inter-region and inter-host values of MPD (equivalent to phylogenetic β diversity), corresponding to the mean phylogenetic distance among all pairs of CoVs from two distinct hosts or regions, and their SES were estimated using the function *comdist* of the R package phylocomr^97^. The matrices of inter-region and inter-host MPD were used to cluster zoogeographic regions and bat hosts in a dendrogram according to their evolutionary similarity (phylo-ordination) using the function *hclust* with complete linkage method of the R package stats (R core team). These computations were performed on both the complete dataset and random subset.

### Mantel tests and isolation by distance

Mantel tests performed in ARLEQUIN 3.5^98^ were used to compare the matrix of viral genetic differentiation (*F*_ST_) to matrices of host phylogenetic distance and geographic distance in order to evaluate the role of geographic isolation and host phylogeny in shaping CoV population structure. The correlation between these matrices was assessed using 10,000 permutations. To gain more resolution into the process of evolutionary diversification, these analyses were also performed at the host genus and province levels. To calculate phylogenetic distances among bat genera, we reconstructed a phylogenetic tree including a single sequence for all bat species included in our dataset. Pairwise patristic distances among tips were computed using the function *distTips* in the R package adephylo^99^. We then averaged all distances across genera to create a matrix of pairwise distances among bat genera. Pairwise Euclidian distances were measured between province centroids and log transformed. Mantel tests were performed with and without genera and provinces including less than four viral sequences to assess the impact of low sample size on our results.

## Data availability

GenBank accession numbers of sequences generated in this study and previously published sequences included in our analysis are available in the Supplementary Note 1 and Supplementary Tables 34 and 35.

## Acknowledgements

This study was funded by the National Institute of Allergy and Infectious Diseases of the National Institutes of Health (Award Number R01AI110964) and the United States Agency for International Development (USAID) Emerging Pandemic Threats PREDICT project (cooperative agreement number GHN-A-OO-09-00010-00), the strategic priority research program of the Chinese Academy of Sciences (XDB29010101), and National Natural Science Foundation of China (31770175, 31830096). Coronavirus research in L-FW’s group is funded by grants from Singapore National Research Foundation (NRF2012NRF-CRP001-056 and NRF2016NRF-NSFC002-013).

## Author contributions

K.J.O., H.E.F, J.H.E., L-F.W., Z.S. and P.D. created the study design, initiated field work and set up sample collection and testing protocols. B.H., G.Z., L.Z., H.L., A.A.C and Z.L. collected samples or provided data. B.H., B.L., and W.Z. performed laboratory work. A.L. carried out the analyses and drafted the manuscript with K.J.O, C.Z.-T. and P.D. All authors reviewed and edited the manuscript

