## Supplementary Information for "Origin and cross-species transmission of bat coronaviruses in China"

**Latinne *et al.***

### Supplementary Information

### Supplementary Note 1

#### GenBank accession numbers of sequences generated for this study

MN312240-MN312869

#### GenBank accession numbers of additional sequences included in this study

Bat-CoVs: AY594268, DQ022305, DQ071615, DQ084199-DQ084200, DQ412042-DQ412043, DQ648794-DQ648795, DQ648809, DQ648819, DQ648822-DQ648823, DQ648833-DQ648835, DQ648837, DQ648850, DQ648856-DQ648858, DQ666337-DQ666340, EF065506, EF065513-EF065516, EF203064-EF203067, EU420137-EU420138, FJ588686, GQ153539-GQ153548, HM211098-HM211101, JQ989266-JQ989273, JX993987-JX993988, KC522036-KC522048, KC522075-KC522089, KC881005-KC881006, KF294268-KF294282, KF294373-KF294378, KF294381-KF294383, KF294420-KF294457, KF367457, KF569973-KF569996, KF636752, KJ473795-KJ473804, KJ473806-KJ473816, KJ473820-KJ473822, KP876505-KP876510, KP876512-KP876515, KP876517-KP876528, KP876532-KP876534, KP876536-KP876537, KP876540-KP876542, KP876544-KP876546, KP886808-KP886809, KP895482-KP895494, KP895496, KP895498-KP895525, KT381902-KT381925, KT444582, KU182954-KU182968, KU182970-KU183003, KU343190-KU343196, KU343198, KU343200, KX285115-KX285179, KX285183, KX285185-KX285197, KX285199-KX285220, KX285223, KX442564, KX447544-KX447565, KY009612-KY009634, KY770850-KY770860, KY383882, KY417142-KY417152, KY783855-KY783881, KY783883-KY783903, MF760455, MF760515, MF769447-MF769451, MF769453, MF769466-MF769470, MF769475-MF769478, MF769485, MF769508-MF769511, MF769513, MG021452, MG762654-MG762656, MG762658

Human CoVs: MN908947, MN975262, NC_004718

#### GISAID accession numbers of additional sequences included in this study

Pangolin CoVs: EPI_ISL_410538-410544, EPI_ISL_410721

Bat-CoV: EPI_ISL_412977

### Supplementary Figures

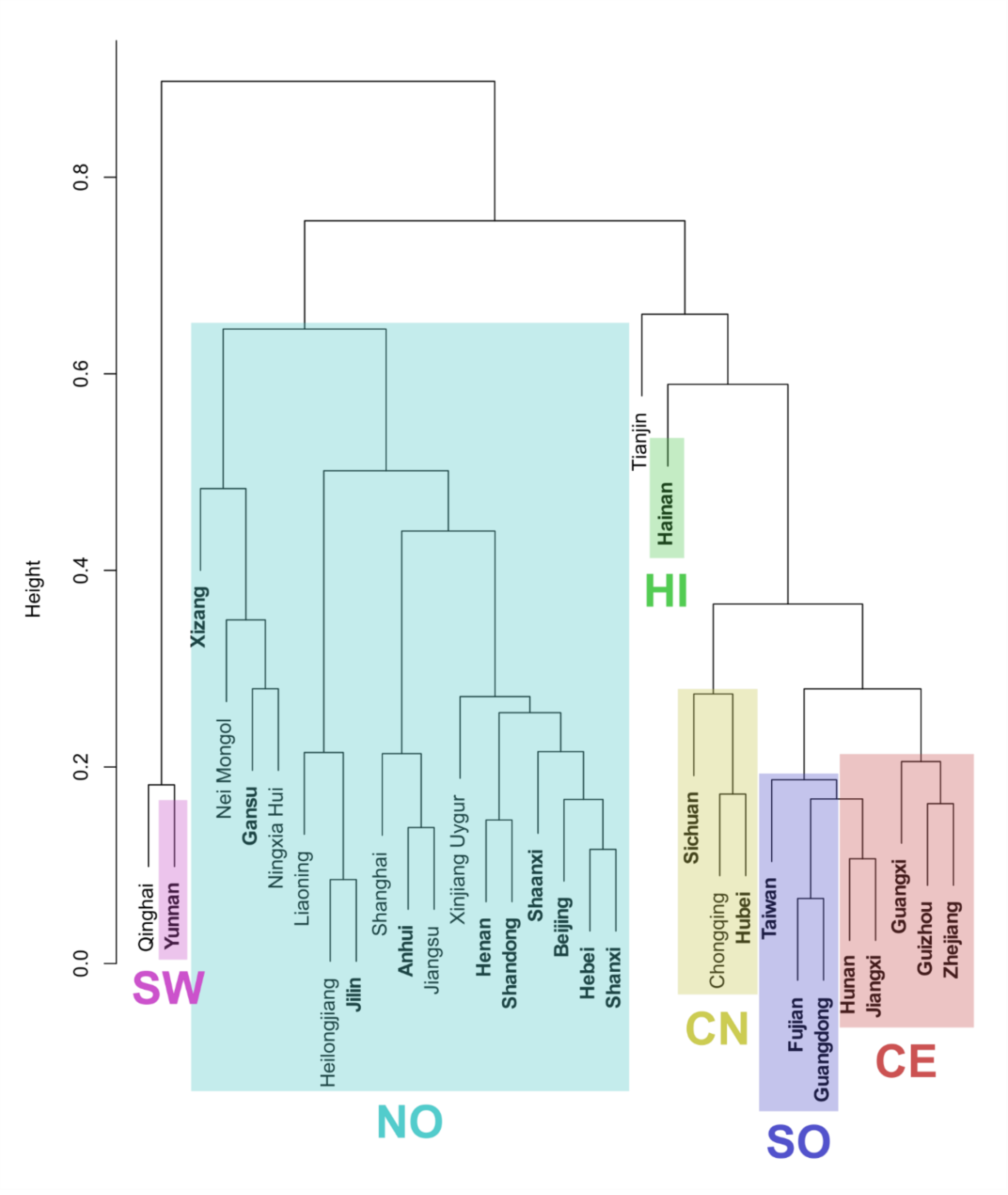

**Supplementary Figure 1**: Cluster dendrogram of Chinese provinces based on similarities between their mammalian diversity (hierarchical clustering). Provinces with CoV sequences available in this study are highlighted in bold.

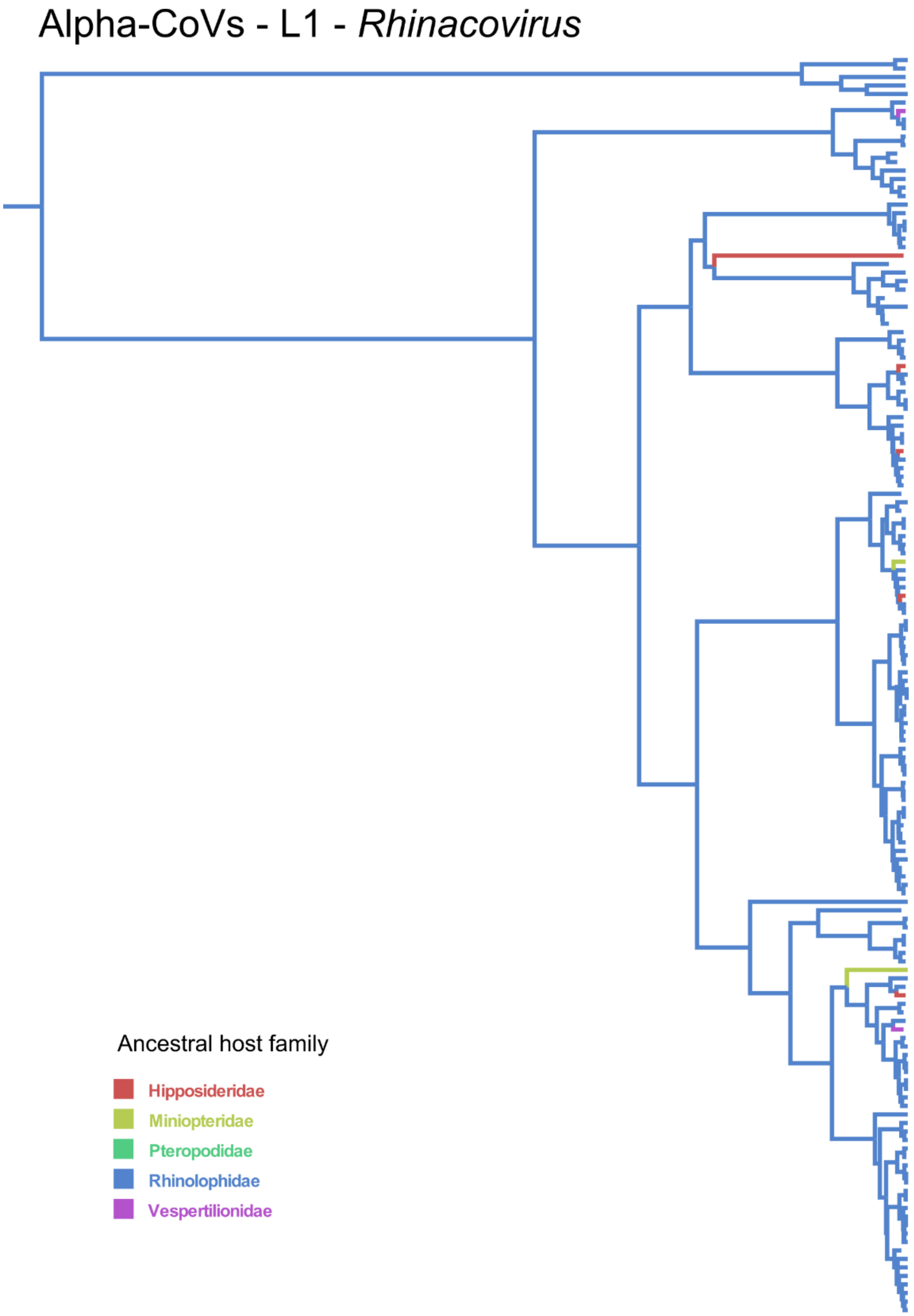

**Supplementary Figure 2**: Zoomed in alpha-CoV maximum clade credibility annotated tree focusing on Lineage L1 corresponding to *Rhinacovirus* and obtained using complete dataset of RdRp sequences and bat host family as discrete character state. Branch colors correspond to the inferred ancestral family with the highest probability.

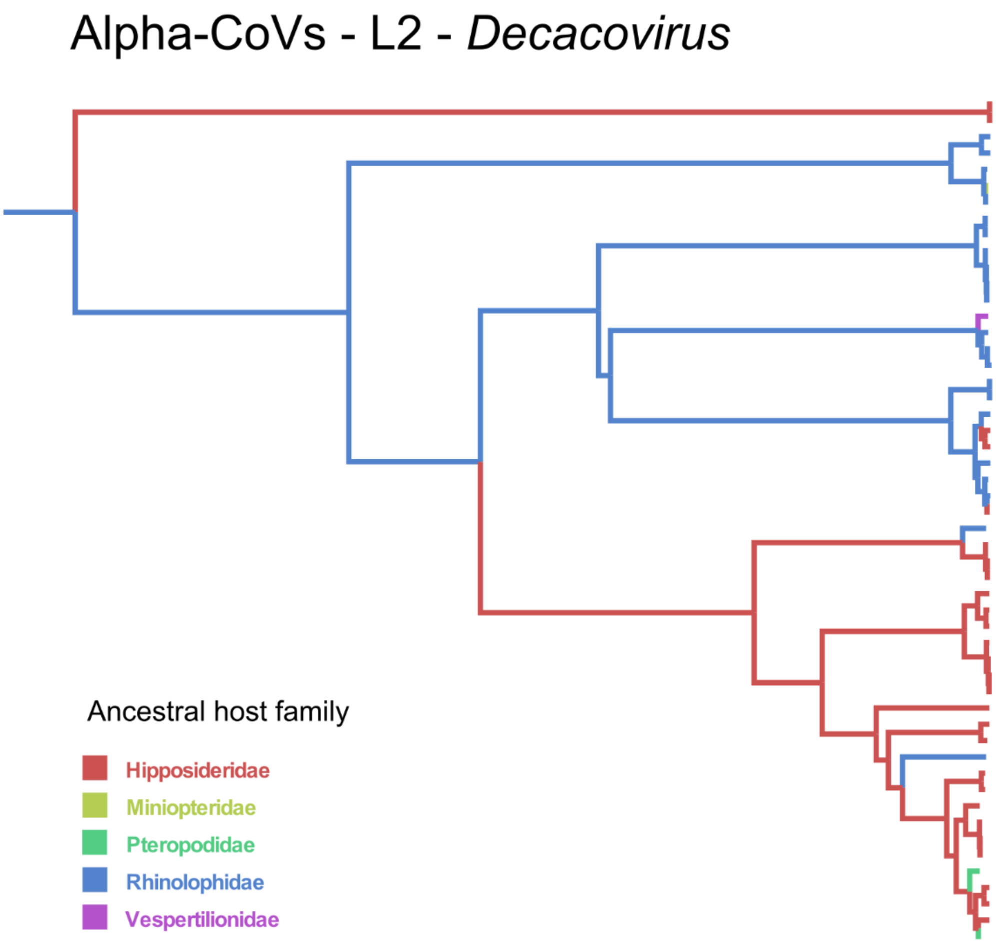

**Supplementary Figure 3:** Zoomed in alpha-CoV maximum clade credibility annotated tree focusing on Lineage L2 corresponding to *Decacovirus* and obtained using complete dataset of RdRp sequences and bat host family as discrete character state. Branch colors correspond to the inferred ancestral family with the highest probability.

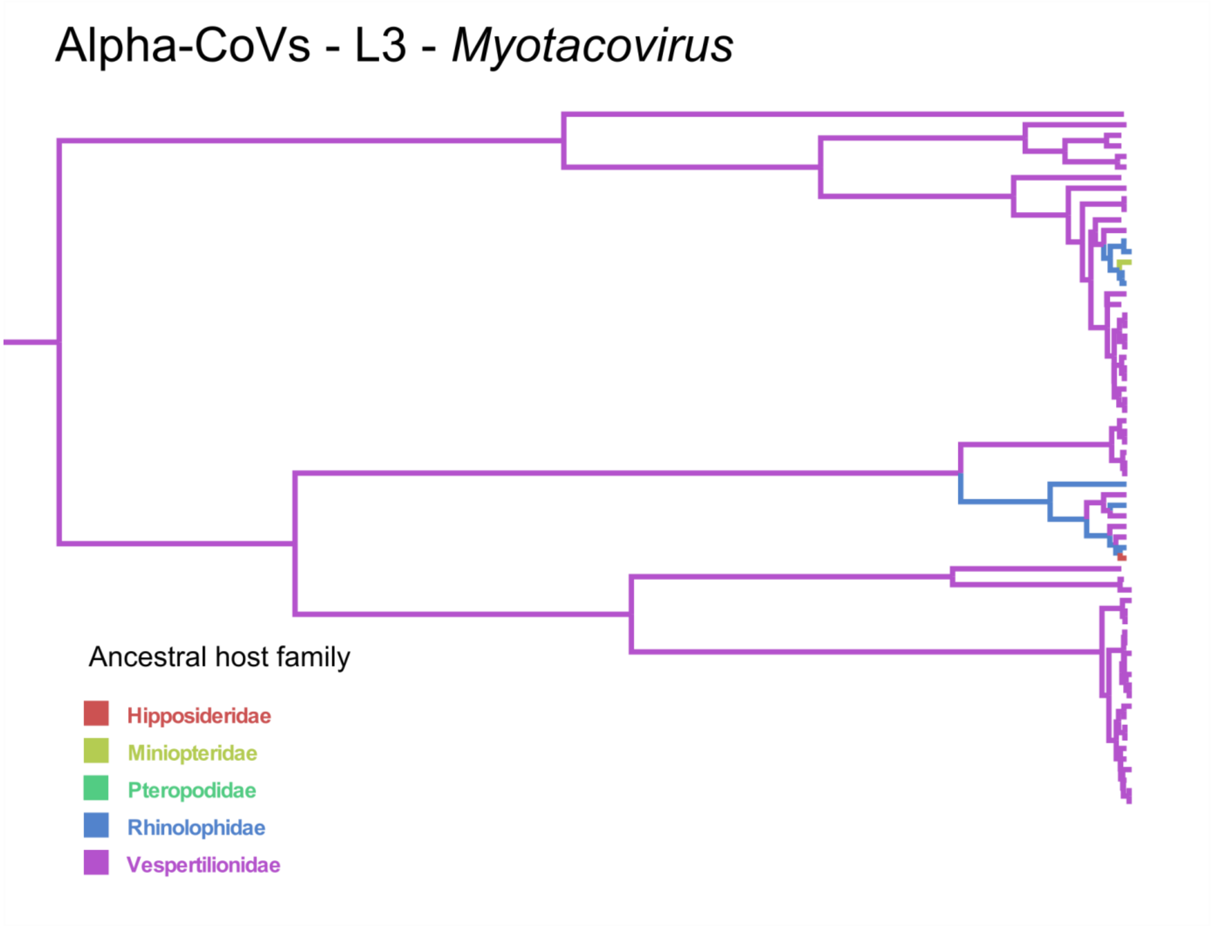

**Supplementary Figure 4:** Zoomed in alpha-CoV maximum clade credibility annotated tree focusing on Lineage L3 corresponding to *Myotacovirus* and obtained using complete dataset of RdRp sequences and bat host family as discrete character state. Branch colors correspond to the inferred ancestral family with the highest probability.

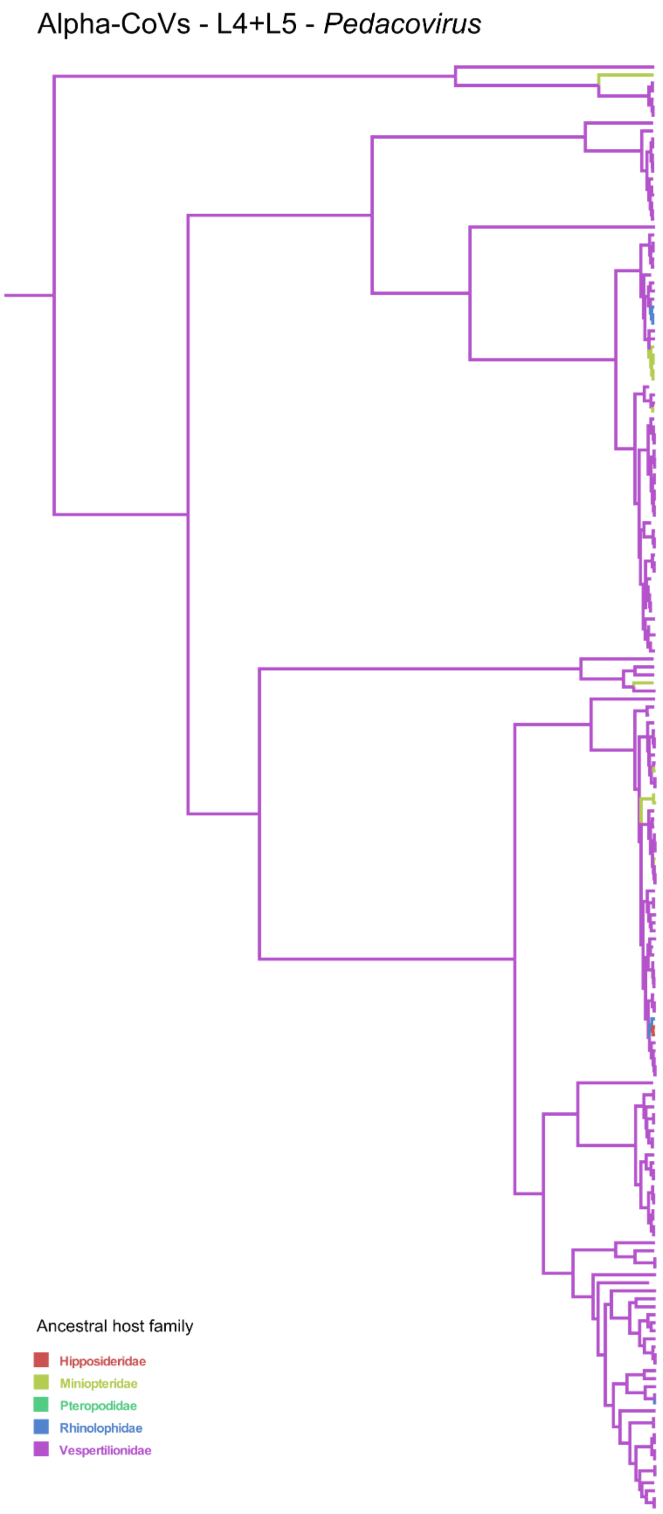

**Supplementary Figure 5:** Zoomed in alpha-CoV maximum clade credibility annotated tree focusing on Lineages L4 and L5 corresponding to *Pedacovirus* and obtained using complete dataset of RdRp sequences and bat host family as discrete character state. Branch colors correspond to the inferred ancestral family with the highest probability.

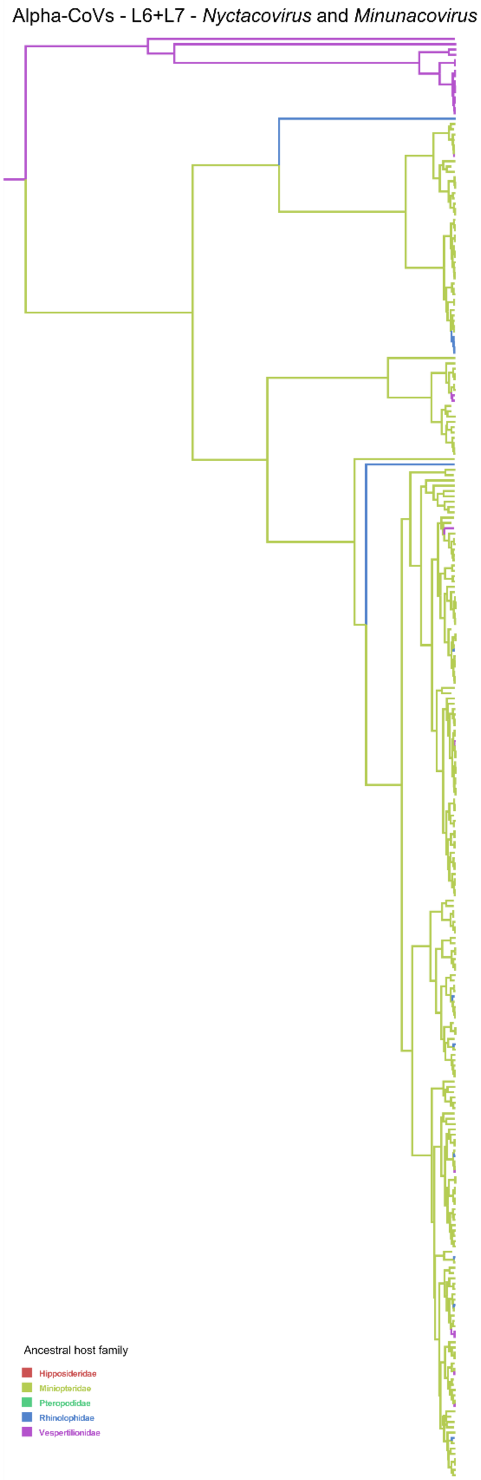

**Supplementary Figure 6:** Zoomed in alpha-CoV maximum clade credibility annotated tree focusing on Lineages L6 and L7 corresponding to *Nyctacovirus* and *Minunacovirus* and obtained using complete dataset of RdRp sequences and bat host family as discrete character state. Branch colors correspond to the inferred ancestral family with the highest probability.

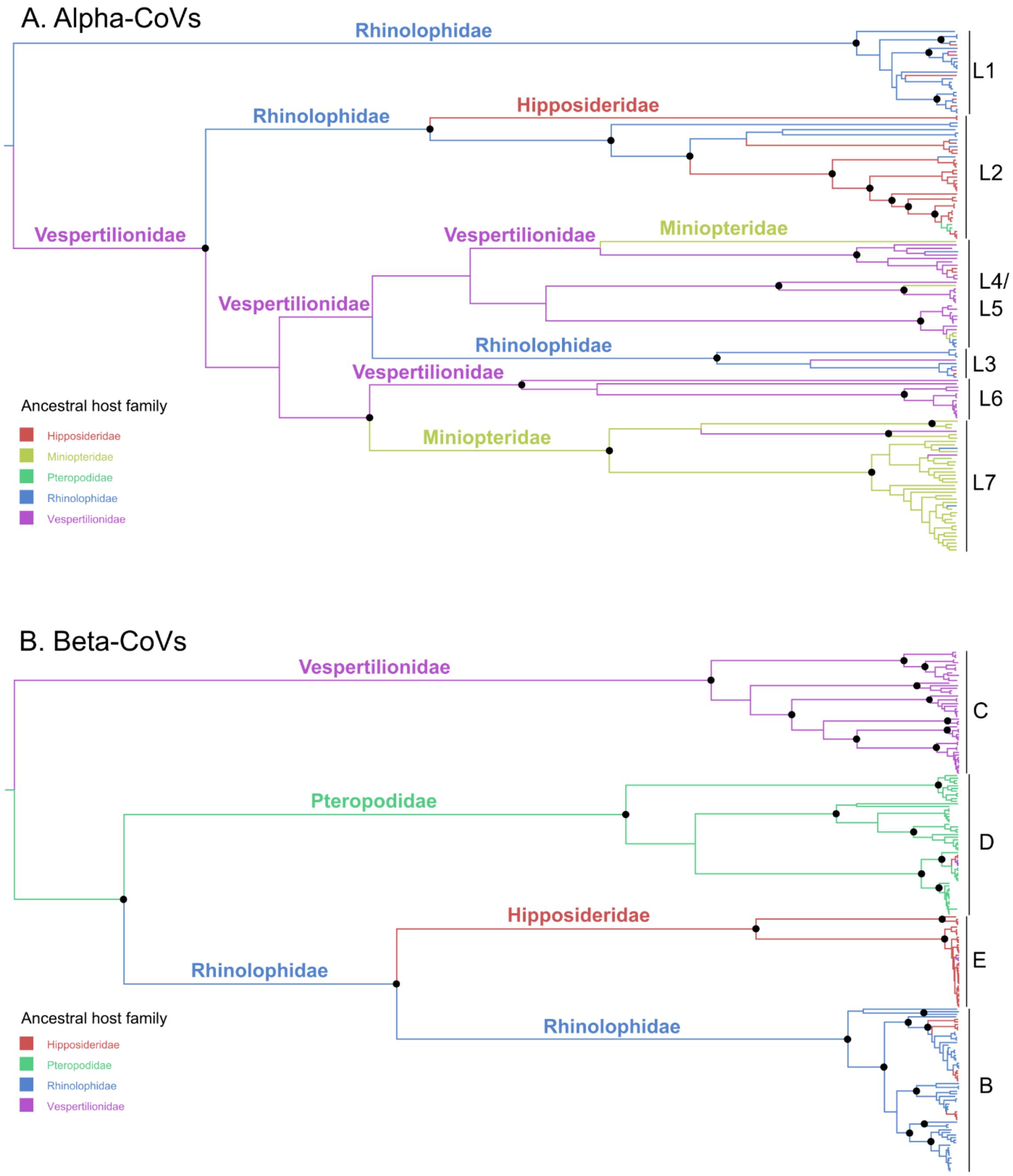

**Supplementary Figure 7**: Alpha-CoV (A) and beta-CoV (B) maximum clade credibility annotated trees using random subset of RdRp sequences to normalize sampling effort across bat families and bat host family as discrete character state. Branches colors correspond to the inferred ancestral family with the highest probability. Branch lengths are scaled according to time. Well-supported nodes (posterior probability > 0.95) are indicated with a black dot. The ICTV approved CoV subgenera were highlighted: Rhinacovirus (L1), Decacovirus (L2), Myotacovirus (L3), Pedacovirus (L5), Nyctacovirus (L6), Minunacovirus (L7) and an unidentified lineage (L4) for alpha-CoVs; and Merbecovirus (Lineage C), Nobecovirus (lineage D), Hibecovirus (lineage E) and Sarbecovirus (Lineage B) for beta-CoVs.

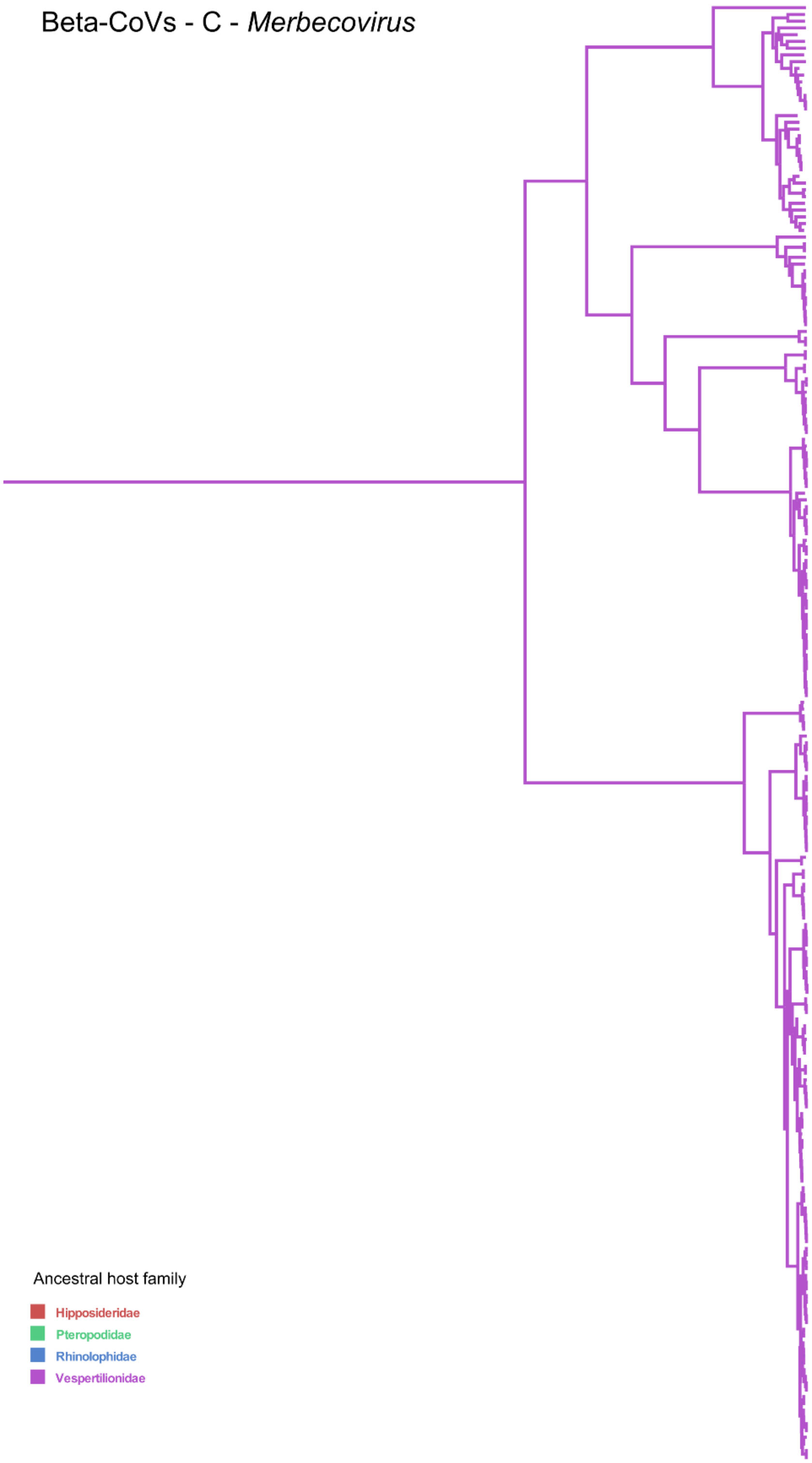

**Supplementary Figure 8:** Zoomed in beta-CoV maximum clade credibility annotated tree focusing on Lineage C corresponding to *Merbecovirus* and obtained using complete dataset of RdRp sequences and bat host family as discrete character state. Branch colors correspond to the inferred ancestral family with the highest probability.

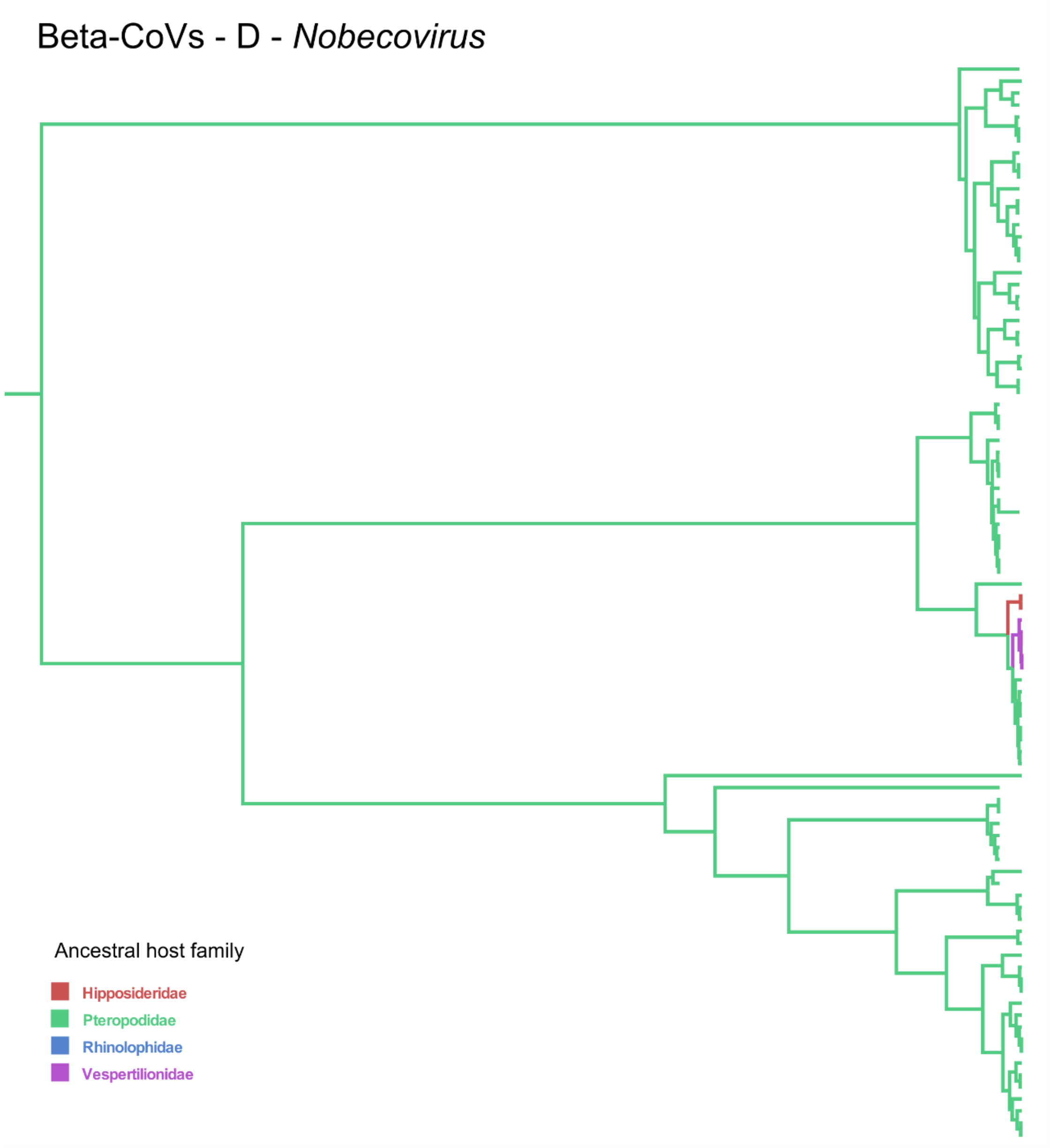

**Supplementary Figure 9:** Zoomed in beta-CoV maximum clade credibility annotated tree focusing on Lineage D corresponding to *Nobecovirus* and obtained using complete dataset of RdRp sequences and bat host family as discrete character state. Branch colors correspond to the inferred ancestral family with the highest probability.

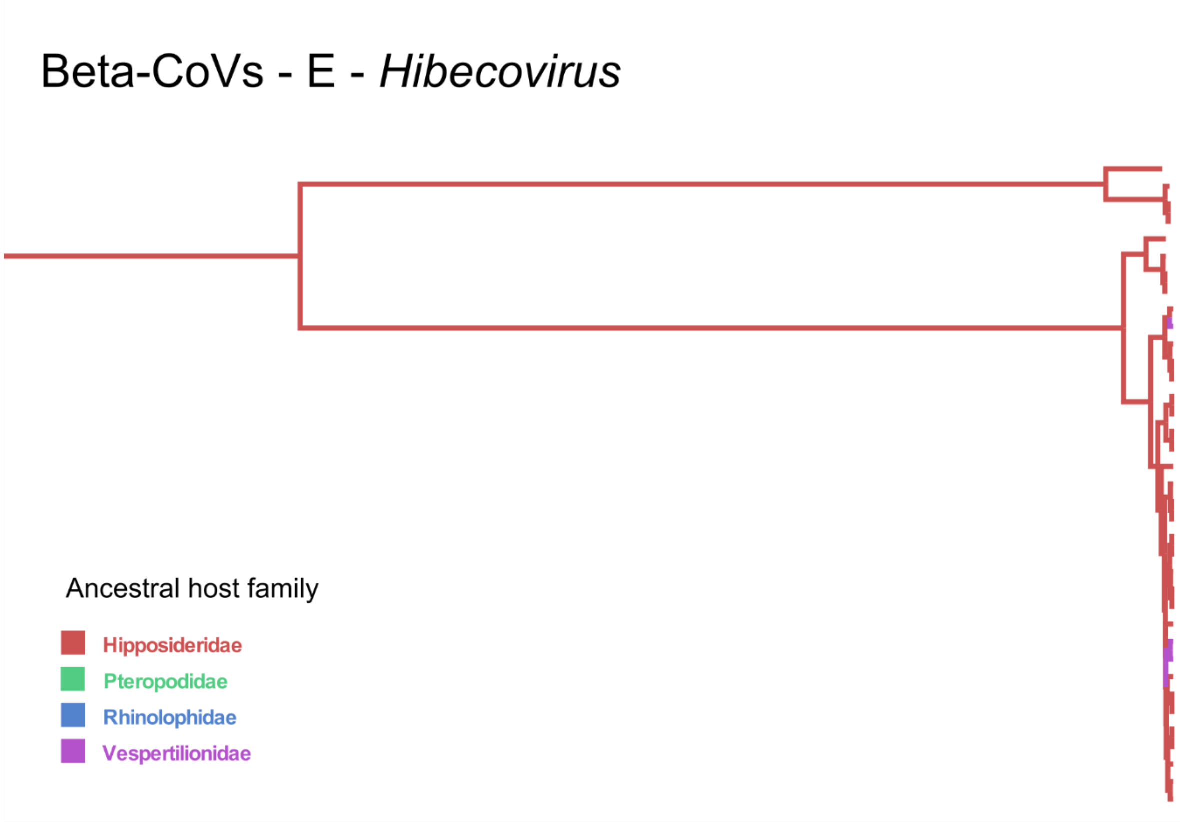

**Supplementary Figure 10:** Zoomed in beta-CoV maximum clade credibility annotated tree focusing on Lineage E corresponding to *Hibecovirus* and obtained using complete dataset of RdRp sequences and bat host family as discrete character state. Branch colors correspond to the inferred ancestral family with the highest probability.

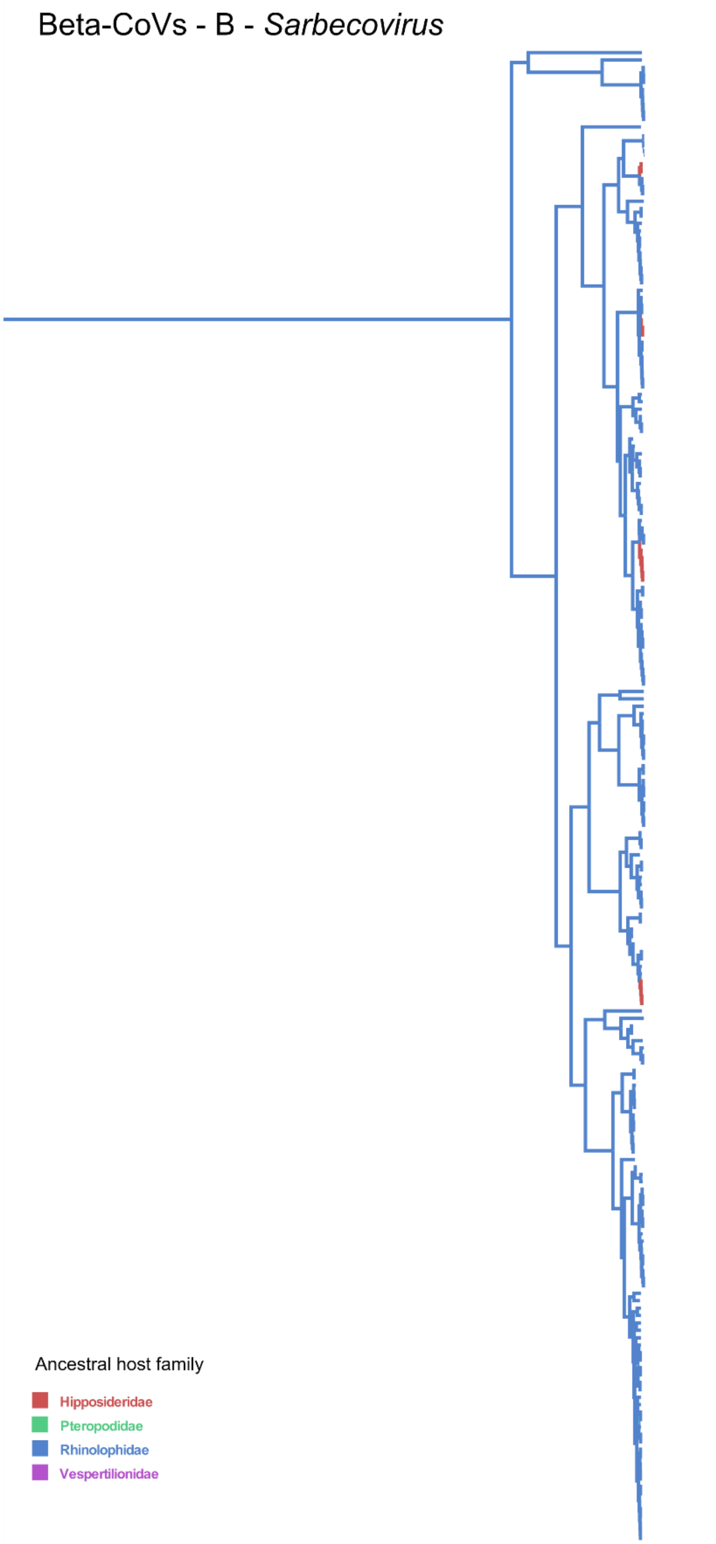

**Supplementary Figure 11:** Zoomed in beta-CoV maximum clade credibility annotated tree focusing on Lineage C corresponding to *Sarbecovirus* and obtained using complete dataset of RdRp sequences and bat host family as discrete character state. Branch colors correspond to the inferred ancestral family with the highest probability.

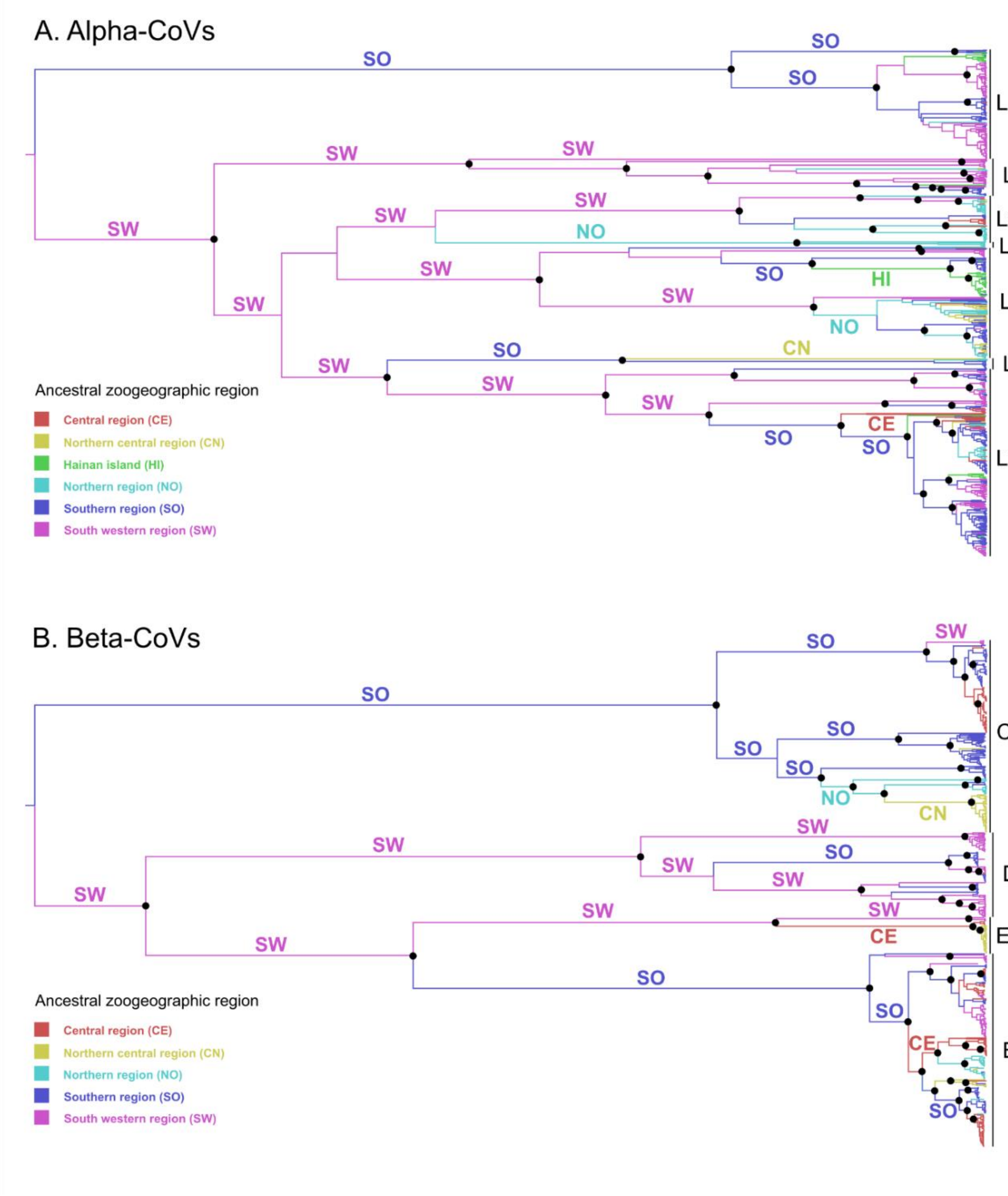

**Supplementary Figure 12**: Alpha-CoV (A) and beta-CoV (B) maximum clade credibility annotated trees using complete datasets of RdRp sequences and zoogeographic regions as discrete character state. Branches colors correspond to the inferred zoogeographic region with the highest probability. Branch lengths are scaled according to time. Well-supported nodes (posterior probability > 0.95) are indicated with a black dot. The ICTV approved CoV subgenera were highlighted: Rhinacovirus (L1), Decacovirus (L2), Myotacovirus (L3), Pedacovirus (L5), Nyctacovirus (L6), Minunacovirus (L7) and an unidentified lineage (L4) for alpha-CoVs; and Merbecovirus (Lineage C), Nobecovirus (lineage D), Hibecovirus (lineage E) and Sarbecovirus (Lineage B) for beta-CoVs. NO, Northern region; CN, Central northern region; SW, South western region; CE, Central region; SO, Southern region; HI, Hainan island

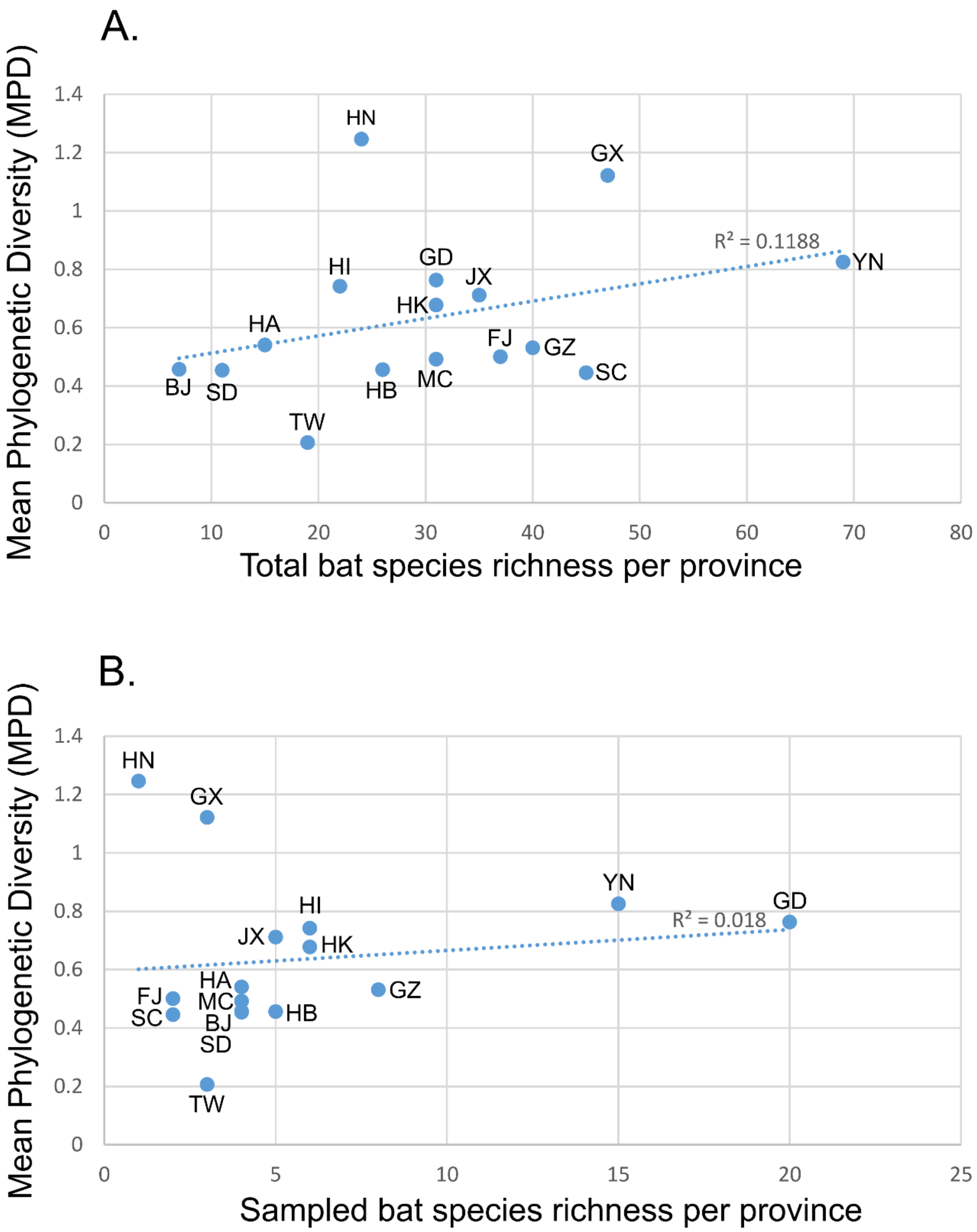

**Supplementary Figure 13:** Scatter plot and linear regression between alpha-CoV phylogenetic diversity (MPD) and total (A) or sampled (B) bat species richness per province using the complete dataset. BJ, Beijing; FJ, Fujian; GD, Guangdong; GX, Guangxi; GZ, Guizhou; HA, Henan; HB, Hubei; HI, Hainan; HK, Hong Kong; HN, Hunan; JX, Jiangxi; MC, Macau; SC, Sichuan; SD, Shandong; TW, Taiwan; YN, Yunnan.

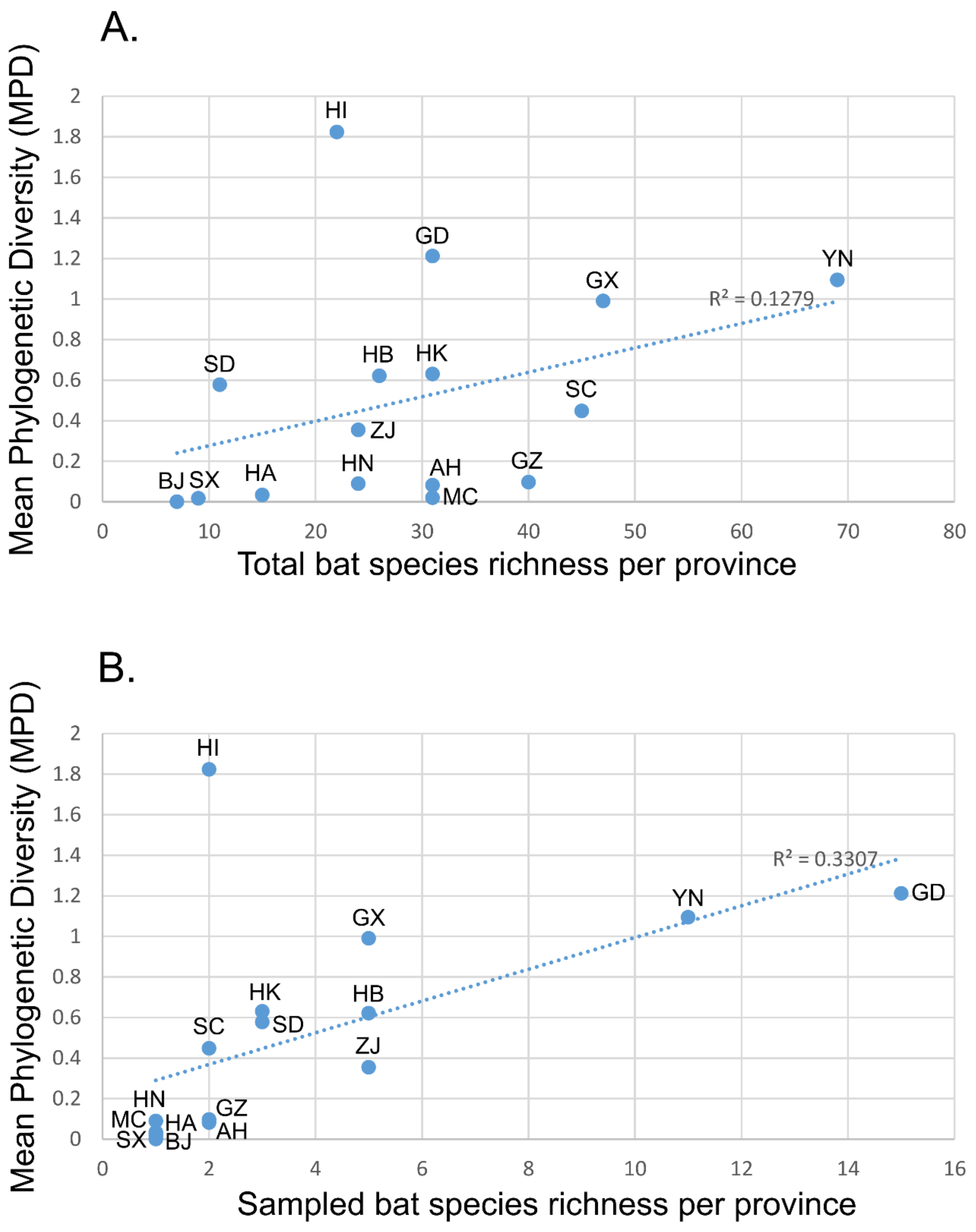

**Supplementary Figure 14:** Scatter plot and linear regression between beta-CoV phylogenetic diversity (MPD) and total (A) or sampled (B) bat species richness per province using the complete dataset. AH, Anhui; BJ, Beijing; GD, Guangdong; GX, Guangxi; GZ, Guizhou; HA, Henan; HB, Hubei; HI, Hainan; HK, Hong Kong; HN, Hunan; MC, Macau; SC, Sichuan; SD, Shandong; SX, Shanxi; YN, Yunnan; ZJ, Zhejiang.

### Supplementary Tables

**Supplementary Table 1**: Total numbers of RdRp sequences available for all CoV lineages from each bat species included in this study. Numbers in parentheses correspond to the number of sequences retrieved from GenBank.

| | Beta-CoVs | | | | Alpha-CoVs | | | | | | |
|---|---|---|---|---|---|---|---|---|---|---|---|
| | B | C | D | E | L1 | L2 | L3 | L4 | L5 | L6 | L7 |
| **Pteropodidae** | | | | | | | | | | | |
| *Cynopterus sphinx* | | | 23 (14) | | | | | | | | |
| *Eonycteris spelaea* | | | 11 | | | | | | | | |
| *Megaerops sp.* | | | 1 (1) | | | | | | | | |
| *Rousettus leschenaultii* | | | 35 (35) | | | 2 (2) | | | | | |
| *Rousettus* sp. | | | 13 | | | | | | | | |
| **Hipposideridae** | | | | | | | | | | | |
| *Aselliscus stoliczkanus* | 4 (1) | | | | | 2 | | | 2 | | |
| *Hipposideros armiger* | 4 (2) | | 2 | 1 | 2 | 2 | 1 (1) | | | | |
| *Hipposideros cineraceus* | | | | | 1 (1) | | | | | | |
| *Hipposideros pomona* | 3 | | | 4 (1) | 1 | 23 (10) | | | | | |
| *Hipposideros pratti* | 1 | | | 29 (1) | 1 (1) | | | | | | |
| **Rhinolophidae** | | | | | | | | | | | |
| *Rhinolophus affinis* | 5 (3) | | | | 59 (9) | 5 (1) | | | | | 1 (1) |
| *Rhinolophus ferrumequinum* | 20 (12) | | | | 1 (1) | 6 (5) | | | | | |
| *Rhinolophus hipposideros* | | | | | | 1 (1) | | | | | |
| *Rhinolophus macrotis* | 2 (2) | | | | 1 | | 1 (1) | | | | |
| *Rhinolophus monoceros* | | | | | | | | | 3 (3) | | |
| *Rhinolophus pearsonii* | 2 (2) | | | | | | 1 (1) | | | | 1 (1) |
| *Rhinolophus pusillus* | 37 (28) | | | | 6 (4) | | 1 (1) | | 1 (1) | | 2 (2) |
| *Rhinolophus rex* | 1 (1) | | | | 7 (7) | | | | | | |
| *Rhinolophus sinicus* | 107 (47) | | | | 27 (20) | 6 (6) | 2 (2) | | | | 8 (5) |
| *Rhinolophus* | | | | | 1 (1) | | | | | | |

| | | | | | | | | | | | |
|---|---|---|---|---|---|---|---|---|---|---|---|
| *stheno* | | | | | | | | | | | |
| *Rhinolophus thomasi* | 1 (1) | | | | | | | | | | |
| *Rhinolophus* sp. | 9 | | | | 37 | 3 | | | 1 | | 1 |
| **Miniopteridae** | | | | | | | | | | | |
| *Miniopterus schreibersii* | | | | | 2 | | 1 (1) | 1 (1) | 10 (9) | | 139 (80) |
| *Miniopterus fuscus* | | | | | | 1 (1) | | | | | 11 (2) |
| *Miniopterus magnater* | | | | | | | | | | | 4 (4) |
| *Miniopterus pusillus* | | | | | | | | | | | 34 (2) |
| *Miniopterus* sp. | | | | | | | | | 1 | | 38 (14) |
| **Vespertilionidae** | | | | | | | | | | | |
| *Eptesicus serotinus* | | 9 (9) | | | | | | | | 2 (2) | |
| *Hypsugo* sp. | | 13 (2) | | | | | | | | 11 (4) | |
| *Ia io* | | 3 (2) | | | | | 1 (1) | | | | |
| *Murina leucogaster* | | | | | | | | 6 (6) | | | |
| *Myotis chinensis* | | | | | | | 1 | | 6 | | |
| *Myotis daubentonii* | | 5 (5) | | | 1 (1) | | 5 (5) | | 28 (28) | | 3 (3) |
| *Myotis davidii* | | | | | | | 10 (10) | | 10 (10) | | 1 (1) |
| *Myotis fimbriatus* | | | | | | | | | 6 (6) | | |
| *Myotis horsfieldii* | | | 1 | | | | | | | | 1 |
| *Myotis myotis* | | | | | | | 1 (1) | | | | |
| *Myotis pequinius* | | 3 (3) | | | | | | | | | |
| *Myotis ricketti* | | 1 (1) | | | 1 (1) | | 24 (21) | | 56 (19) | | 4 (2) |
| *Myotis siligorensis* | | | | | | | 1 (1) | | | | |
| *Myotis* sp. | | | | | | 1 (1) | 6 (6) | | 3 (1) | | |
| *Nyctalus plancyi* | | | | | | | | | | 1 (1) | |
| *Pipistrellus abramus* | | 34 (21) | 3 | | | | | | | | |
| *Pipistrellus pipistrellus* | | 1 (1) | | | | | | | | | |
| *Pipistrellus* sp. | | | 1 | | | | | | | | |
| *Scotophilus kuhlii* | | 1 (1) | | | | | | | 44 (14) | | |

| | | | | | | | | | | | |
|---|---|---|---|---|---|---|---|---|---|---|---|
| *Tylonycteris pachypus* | | 96 (57) | | | | | | | | | 1 |
| *Tylonycteris robustula* | | | | | | | | | | 1 (1) | |
| *Vespertilio sinensis* | | 39 (2) | | 3 | | | | | 3 | | |
| Total | 196 | 205 | 90 | 37 | 148 | 52 | 56 | 7 | 174 | 15 | 249 |

**Supplementary Table 2**: Significant pairwise transitions between bat families for alpha-CoVs and their corresponding Bayes Factor (BF) and posterior probability (PP) using the complete dataset and random subset. Only transitions with BF >10 for at least one dataset are presented.

| Donor family | Recipient family | Complete dataset | | Random dataset | |
|---|---|---|---|---|---|
| | | BF | PP | BF | PP |
| Miniopteridae | Rhinolophidae | 23483.1 | 1 | 28.9 | 0.899 |
| Miniopteridae | Vespertilionidae | 23483.1 | 1 | 3.1 | 0.468 |
| Rhinolophidae | Hipposideridae | 23483.1 | 1 | 148.8 | 0.978 |
| Vespertilionidae | Miniopteridae | 23483.1 | 1 | 14.2 | 0.814 |
| Vespertilionidae | Rhinolophidae | 23483.1 | 1 | 2.9 | 0.469 |
| Rhinolophidae | Miniopteridae | 7825.5 | 0.999 | 0.6 | 0.168 |
| Rhinolophidae | Vespertilionidae | 3351.9 | 0.999 | 20.3 | 0.861 |
| Hipposideridae | Pteropodidae | 2606.3 | 0.999 | 14.5 | 0.816 |
| Hipposideridae | Rhinolophidae | 408.8 | 0.992 | 47.3 | 0.935 |

**Supplementary Table 3**: Significant pairwise transitions between bat families for beta-CoVs and their corresponding Bayes Factor (BF) and posterior probability (PP) using the complete dataset and random subset. Only transitions with BF >10 for at least one dataset are presented.

| Donor family | Recipient family | Complete dataset | | Random dataset | |
|---|---|---|---|---|---|
| | | BF | PP | BF | PP |
| Hipposideridae | Vespertilionidae | 16194.7 | 0.999 | 20243.3 | 0.999 |
| Rhinolophidae | Hipposideridae | 16194.7 | 1 | 20243.3 | 1 |
| Pteropodidae | Hipposideridae | 7.7 | 0.773 | 25.6 | 0.919 |

**Supplementary Table 4**: Number of state changes (Markov jumps) from/to each bat family (donor/receiver) along the significant inter-family transition rates for alpha-CoVs using the complete dataset and random subset.

| Family | Complete dataset | | | Random dataset | | |
|---|---|---|---|---|---|---|
| | Donor | Receiver | Total | Donor | Receiver | Total |
| Hipposideridae | 11 | 13.5 | 24.5 | 28.4 | 24.6 | 53 |
| Rhinolophidae | 28.4 | 33.3 | 61.7 | 35.2 | 31.9 | 67.1 |
| Vespertilionidae | 21.8 | 24.3 | 46.1 | 6.6 | 10.5 | 7.1 |
| Miniopteridae | 31.6 | 19.3 | 50.9 | 8.3 | 6.6 | 14.9 |
| Pteropodidae | 0 | 2.4 | 2.4 | 0 | 4.8 | 4.8 |

**Supplementary Table 5**: Number of state changes (Markov jumps) from/to each bat family (donor/receiver) along the significant inter-family transition rates for beta-CoVs using the complete dataset and random subset.

| Family | Complete dataset | | | Random dataset | | |
|---|---|---|---|---|---|---|
| | Donor | Receiver | Total | Donor | Receiver | Total |
| Hipposideridae | 3.4 | 7.9 | 11.3 | 2.4 | 6.9 | 9.3 |
| Rhinolophidae | 6.3 | 0 | 6.3 | 5.3 | 0 | 5.3 |
| Vespertilionidae | 0 | 3.4 | 3.4 | 0 | 2.4 | 2.4 |
| Pteropodidae | 1.6 | 0 | 1.6 | 1.6 | 0 | 1.6 |

**Supplementary Table 6**: Significant pairwise transitions between bat genera for alpha-CoVs and their corresponding Bayes Factor (BF) and posterior probability (PP) using the complete dataset and random subset. Only transitions with BF >10 for at least one dataset are presented.

| Donor genus | Recipient genus | Complete dataset | | Random dataset | |
|---|---|---|---|---|---|
| | | BF | PP | BF | PP |
| *Miniopterus* | *Myotis* | 88497.5 | 1 | 1410.2 | 0.991 |
| *Miniopterus* | *Rhinolophus* | 88497.5 | 1 | 440.3 | 0.973 |
| *Rhinolophus* | *Hipposideros* | 88497.5 | 1 | 99559.7 | 1 |
| *Myotis* | *Miniopterus* | 11051.4 | 0.999 | 5844.9 | 0.998 |
| *Scotophilus* | *Miniopterus* | 2669.8 | 0.996 | 21.1 | 0.632 |
| *Rhinolophus* | *Myotis* | 1540.5 | 0.993 | 771.7 | 0.984 |
| *Myotis* | *Vespertilio* | 1438.7 | 0.992 | 167.4 | 0.932 |
| *Rhinolophus* | *Miniopterus* | 1167.8 | 0.99 | 0.4 | 0.034 |
| *Hipposideros* | *Aselliscus* | 322.9 | 0.963 | 1734. 6 | 0.993 |
| *Myotis* | *Rhinolophus* | 242.1 | 0.952 | 11.4 | 0.481 |
| *Hipposideros* | *Rousettus* | 112.7 | 0.902 | 523.1 | 0.977 |
| *Miniopterus* | *Tylonycteris* | 102.4 | 0.893 | 1351.7 | 0.991 |
| *Scotophilus* | *Myotis* | 45.3 | 0.787 | 170.7 | 0.933 |
| *Myotis* | *Aselliscus* | 22.9 | 0.651 | 1.9 | 0.132 |
| *Rhinolophus* | *Ia* | 21.7 | 0.639 | 1.2 | 0.091 |
| *Aselliscus* | *Rhinolophus* | 17.8 | 0.591 | 1.8 | 0.128 |
| *Murina* | *Miniopterus* | 16.5 | 0.573 | 1.4 | 0.099 |
| *Hypsugo* | *Eptesicus* | 15.4 | 0.556 | 15.3 | 0.555 |
| *Hipposideros* | *Rhinolophus* | 10.7 | 0.465 | 134.6 | 0.916 |
| *Eptesicus* | *Hypsugo* | 10.2 | 0.455 | 10.8 | 0.467 |
| *Vespertilio* | *Aselliscus* | 3.9 | 0.242 | 68.8 | 0.848 |
| *Ia* | *Hipposideros* | 1.4 | 0.106 | 10.4 | 0.459 |

**Supplementary Table 7**: Significant pairwise transitions between bat genera for beta-CoVs and their corresponding Bayes Factor (BF) and posterior probability (PP) using the complete dataset and random subset. Only transitions with BF >10 for at least one dataset are presented.

| Donor genus | Recipient genus | Complete dataset | | Random dataset | |
|---|---|---|---|---|---|
| | | BF | PP | BF | PP |
| *Rousettus* | *Eonycteris* | 19130.4 | 0.999 | 95425.9 | 1 |
| *Rhinolophus* | *Hipposideros* | 4544.7 | 0.997 | 47706.3 | 0.999 |
| *Hipposideros* | *Vespertilio* | 2887.3 | 0.995 | 10591.1 | 0.999 |
| *Tylonycteris* | *Scotophilus* | 1354.1 | 0.990 | 10591.1 | 0.999 |
| *Eonycteris* | *Rousettus* | 380.6 | 0.966 | 13.4 | 0.502 |
| *Hypsugo* | *Myotis* | 211.9 | 0.941 | 862.3 | 0.985 |
| *Ia* | *Pipistrellus* | 68.9 | 0.838 | 160.5 | 0.923 |
| *Rousettus* | *Megaerops* | 63.5 | 0.827 | 166.8 | 0.926 |
| *Cynopterus* | *Myotis* | 31.9 | 0.706 | 28.8 | 0.684 |

| | | | | | |
|---|---|---|---|---|---|
| *Cynopterus* | *Rousettus* | 31.4 | 0.703 | 816.6 | 0.984 |
| *Cynopterus* | *Pipistrellus* | 22.9 | 0.633 | 70.5 | 0.841 |
| *Cynopterus* | *Hipposideros* | 20.8 | 0.611 | 29.3 | 0.688 |
| *Myotis* | *Eptesicus* | 18.4 | 0.580 | 15.1 | 0.533 |
| *Hipposideros* | *Aselliscus* | 14.3 | 0.519 | 5.1 | 0.279 |
| *Rhinolophus* | *Aselliscus* | 11.3 | 0.461 | 37.3 | 0.737 |
| *Eptesicus* | *Myotis* | 11.2 | 0.457 | 12.2 | 0.479 |

**Supplementary Table 8**: Number of state changes (Markov jumps) from/to each bat genus (donor/receiver) along the significant inter-family transition rates for alpha-CoVs using the complete dataset and random subset.

| Genus | Complete dataset | | | Random dataset | | |
|---|---|---|---|---|---|---|
| | Donor | Receiver | Total | Donor | Receiver | Total |
| *Aselliscus* | 1.3 | 5.6 | 6.9 | 0.7 | 2.7 | 3.4 |
| *Eptesicus* | 1 | 1.3 | 2.3 | 0.7 | 0.8 | 1.5 |
| *Hipposideros* | 10.9 | 17.9 | 28.8 | 6.6 | 10.5 | 17.1 |
| *Hypsugo* | 1.3 | 1 | 2.3 | 0.8 | 0.7 | 1.5 |
| *Ia* | 0 | 1.2 | 1.2 | 0 | 1.3 | 1.3 |
| *Miniopterus* | 37.3 | 23.4 | 60.7 | 5.8 | 4.2 | 10 |
| *Murina* | 1.7 | 0 | 1.7 | 1.2 | 0 | 1.2 |
| *Myotis* | 25.4 | 26.6 | 52 | 4.4 | 2.7 | 7.1 |
| *Nyctalus* | 0 | 0 | 0 | 0 | 0 | 0 |
| *Rhinolophus* | 36.8 | 37.7 | 74.5 | 13.8 | 9 | 22.8 |
| *Rousettus* | 0 | 2.3 | 2.3 | 0 | 1.7 | 1.7 |
| *Scotophilus* | 7.1 | 0 | 7.1 | 2.3 | 0 | 2.3 |
| *Tylonycteris* | 0 | 2 | 2 | 0 | 1.4 | 1.4 |
| *Vespertilio* | 0 | 3.9 | 3.9 | 0 | 1.3 | 1.3 |

**Supplementary Table 9**: Number of state changes (Markov jumps) from/to each bat genus (donor/receiver) along the significant inter-family transition rates for beta-CoVs using the complete dataset and random subset.

| Genus | Complete dataset | | | Random dataset | | |
|---|---|---|---|---|---|---|
| | Donor | Receiver | Total | Donor | Receiver | Total |
| *Aselliscus* | 0 | 2.9 | 2.9 | 0 | 1.4 | 1.4 |
| *Cynopterus* | 12.3 | 0 | 12.3 | 5.4 | 0 | 5.4 |
| *Eonycteris* | 1.2 | 20.2 | 21.4 | 4.1 | 2.8 | 6.9 |
| *Eptesicus* | 1.7 | 1.9 | 3.6 | 0.9 | 0.8 | 1.7 |
| *Hipposideros* | 5.8 | 14.5 | 20.3 | 1.8 | 5.3 | 7.1 |
| *Hypsugo* | 3.1 | 0 | 3.1 | 1.5 | 0 | 1.5 |
| *Ia* | 3.4 | 0 | 3.4 | 1.4 | 0 | 1.4 |
| *Megaerops* | 0 | 2.9 | 2.9 | 0 | 0.2 | 0.2 |
| *Myotis* | 1.9 | 7.5 | 9.4 | 0.8 | 3.5 | 4.3 |
| *Pipistrellus* | 0 | 6.3 | 6.3 | 0 | 2.8 | 2.8 |
| *Rhinolophus* | 14.2 | 0 | 14.2 | 5.1 | 0 | 5.1 |
| *Rousettus* | 23.1 | 5.1 | 28.2 | 3 | 5.7 | 8.7 |
| *Scotophilus* | 0 | 2.9 | 2.9 | 0 | 1.3 | 1.3 |
| *Tylonycteris* | 2.9 | 0 | 2.9 | 1.3 | 0 | 1.3 |
| *Vespertilio* | 0 | 5.5 | 5.5 | 0 | 1.6 | 1.6 |

**Supplementary Table 10**: Significant pairwise transitions between zoogeographic regions for alpha-CoVs and their corresponding Bayes Factor (BF) and posterior probability (PP) using the complete dataset and random subset. Only transitions with BF >10 for at least one dataset are presented. NO, Northern region; CN, Central northern region; SW, South western region; CE, Central region; SO, Southern region; HI, Hainan island.

| Donor region | Recipient region | Complete dataset | | Random dataset | |
|---|---|---|---|---|---|
| | | BF | PP | BF | PP |
| CN | SO | 30740.3 | 1 | 2.3 | 0.350 |
| HI | SO | 30740.3 | 1 | 2397.6 | 0.998 |
| NO | CN | 2790.7 | 0.998 | 6.2 | 0.592 |
| SO | CE | 649.9 | 0.993 | 13.4 | 0.759 |
| NO | CE | 441.3 | 0.990 | 122.9 | 0.966 |
| SW | SO | 267.8 | 0.984 | 13.3 | 0.757 |
| SW | NO | 136.1 | 0.969 | 0.5 | 0.112 |
| SO | SW | 83.6 | 0.951 | 19.7 | 0.822 |
| CE | SW | 60.4 | 0.934 | 2.4 | 0.362 |
| SO | HI | 58.3 | 0.932 | 5.2 | 0.550 |
| HI | NO | 32.1 | 0.882 | 0.3 | 0.063 |
| CN | NO | 2.9 | 0.405 | 14.2 | 0.769 |

**Supplementary Table 11**: Significant pairwise transitions between zoogeographic regions for beta-CoVs and their corresponding Bayes Factor (BF) and posterior probability (PP) using the complete dataset and random subset. Only transitions with BF >10 for at least one dataset are presented. NO, Northern region; CN, Central northern region; SW, South western region; CE, Central region; SO, Southern region.

| Donor region | Recipient region | Complete dataset | | Random dataset | |
|---|---|---|---|---|---|
| | | BF | PP | BF | PP |
| SO | CE | 835.5 | 0.996 | 122.7 | 0.974 |
| NO | SO | 94.2 | 0.966 | 0.6 | 0.157 |
| SW | SO | 66.0 | 0.953 | 95.2 | 0.967 |
| CE | SW | 45.5 | 0.933 | 132.6 | 0.976 |
| NO | CN | 27.7 | 0.895 | 197.8 | 0.984 |
| CE | NO | 14.3 | 0.814 | 8.8 | 0.730 |
| SO | SW | 10.7 | 0.766 | 1.4 | 0.305 |
| CE | SO | 5.8 | 0.6 | 22.9 | 0.875 |

**Supplementary Table 12**: Number of state changes (Markov jumps) from/to each zoogeographic region (donor/receiver) along the significant dispersal routes for alpha-CoVs using the complete dataset and random subset. NO, Northern region; CN, Central northern region; SW, South western region; CE, Central region; SO, Southern region; HI, Hainan island.

| Region | Complete dataset | | | Random dataset | | |
|---|---|---|---|---|---|---|
| | Donor | Receiver | Total | Donor | Receiver | Total |
| CE | 19.9 | 41.2 | 61.1 | 0 | 41.2 | 41.2 |
| CN | 42.9 | 22.4 | 65.3 | 23.6 | 0 | 23.6 |
| HI | 36.8 | 9.4 | 46.2 | 18.1 | 0 | 18.1 |
| NO | 46.5 | 20.1 | 66.5 | 25.1 | 23.6 | 48.7 |
| SO | 52 | 88.6 | 140.6 | 30.6 | 32.6 | 63.2 |
| SW | 29 | 45.5 | 74.5 | 14.6 | 14.5 | 29.1 |

**Supplementary Table 13**: Number of state changes (Markov jumps) from/to each zoogeographic region (donor/receiver) along the significant dispersal routes for beta-CoVs using the complete dataset and random subset. NO, Northern region; CN, Central northern region; SW, South western region; CE, Central region; SO, Southern region.

| Region | Complete dataset | | | Random dataset | | |
|---|---|---|---|---|---|---|
| | Donor | Receiver | Total | Donor | Receiver | Total |
| CE | 14.8 | 9.6 | 24.2 | 25.4 | 12.1 | 37.5 |
| CN | 0 | 7.5 | 7.5 | 0 | 7.6 | 7.6 |
| NO | 17.4 | 6.0 | 23.4 | 7.6 | 0 | 7.6 |
| SO | 17.1 | 17.4 | 34.5 | 12.1 | 21.4 | 33.5 |
| SW | 7.5 | 16.4 | 23.9 | 8.3 | 12.2 | 20.5 |

**Supplementary Table 14**: Mean Phylogenetic Distance (mpd.obs) and its standardized effect size (mpd.obs.z) within bat families for alpha-CoVs using the complete dataset and random subset. One-tailed p-values (quantiles) were calculated after randomly reshuffling tip labels 1000 times along the entire phylogeny (=mpd.obs.rank/runs+1). Significant p-value (mpd.obs.p) are highlighted in bold.

| | Complete dataset | | | | | | | |
|---|---|---|---|---|---|---|---|---|
| | n | mpd.obs | mpd.rand.mean | mpd.rand.sd | mpd.obs.rank | mpd.obs.z | mpd.obs.p | runs |
| Hipposideridae | 29 | 0.688 | 0.857 | 0.054 | 11 | -3.138 | **0.0110** | 1000 |
| Miniopteridae | 222 | 0.333 | 0.853 | 0.015 | 1 | -34.984 | **0.0010** | 1000 |
| Pteropodidae | 2 | 0.006 | 0.869 | 0.405 | 1 | -2.129 | **0.0010** | 1000 |
| Rhinolophidae | 123 | 0.713 | 0.854 | 0.023 | 1 | -6.167 | **0.0010** | 1000 |
| Vespertilionidae | 174 | 0.634 | 0.853 | 0.018 | 1 | -12.010 | **0.0010** | 1000 |
| | Random subset | | | | | | | |
| | n | mpd.obs | mpd.rand.mean | mpd.rand.sd | mpd.obs.rank | mpd.obs.z | mpd.obs.p | runs |
| Hipposideridae | 29 | 0.697 | 0.921 | 0.037 | 1 | -6.089 | **0.0010** | 1000 |
| Miniopteridae | 39 | 0.326 | 0.920 | 0.029 | 1 | -20.262 | **0.0010** | 1000 |
| Pteropodidae | 2 | 0.007 | 0.903 | 0.383 | 7 | -2.340 | **0.0070** | 1000 |
| Rhinolophidae | 34 | 0.921 | 0.919 | 0.034 | 464 | 0.040 | 0.4640 | 1000 |
| Vespertilionidae | 30 | 0.755 | 0.920 | 0.036 | 1 | -4.515 | **0.0010** | 1000 |

**Supplementary Table 15**: Mean Phylogenetic Distance (mpd.obs) and its standardized effect size (mpd.obs.z) within bat families for beta-CoVs using the complete dataset and random subset. One-tailed p-values (quantiles) were calculated after randomly reshuffling tip labels 1000 times along the entire phylogeny (=mpd.obs.rank/runs+1). Significant p-value (mpd.obs.p) are highlighted in bold.

| | Complete dataset | | | | | | | |
|---|---|---|---|---|---|---|---|---|
| | n | mpd.obs | mpd.rand.mean | mpd.rand.sd | mpd.obs.rank | mpd.obs.z | mpd.obs.p | runs |
| Hipposideridae | 21 | 0.7053 | 1.2273 | 0.0693 | 1 | -7.527 | **0.0010** | 1000 |
| Pteropodidae | 58 | 0.4290 | 1.2315 | 0.0315 | 1 | -25.481 | **0.0010** | 1000 |
| Rhinolophidae | 103 | 0.1187 | 1.2306 | 0.0203 | 1 | -54.789 | **0.0010** | 1000 |
| Vespertilionidae | 117 | 0.4542 | 1.2304 | 0.0184 | 1 | -42.097 | **0.0010** | 1000 |
| | Random subset | | | | | | | |
| | n | mpd.obs | mpd.rand.mean | mpd.rand.sd | mpd.obs.rank | mpd.obs.z | mpd.obs.p | runs |
| Hipposideridae | 21 | 0.724838 | 1.261965 | 0.059781 | 1 | -8.98499 | 0.000999 | 1000 |
| Pteropodidae | 33 | 0.438785 | 1.26173 | 0.041737 | 1 | -19.7172 | 0.000999 | 1000 |
| Rhinolophidae | 34 | 0.123321 | 1.262034 | 0.038973 | 1 | -29.2182 | 0.000999 | 1000 |
| Vespertilionidae | 39 | 0.545279 | 1.260986 | 0.037071 | 1 | -19.3065 | 0.000999 | 1000 |

**Supplementary Table 16**: Mean Nearest Taxon Distance (mntd.obs) and its standardized effect size (mntd.obs.z) within bat families for alpha-CoVs using the complete dataset and random subset. One-tailed p-values (quantiles) were calculated after randomly reshuffling tip labels 1000 times along the entire phylogeny (=mpd.obs.rank/runs+1). Significant p-value (mntd.obs.p) are highlighted in bold.

| | Complete dataset | | | | | | | |
|---|---|---|---|---|---|---|---|---|
| | n | mntd.obs | mntd.rand.mean | mntd.rand.sd | mntd.obs.rank | mntd.obs.z | mntd.obs.p | runs |
| Hipposideridae | 29 | 0.0484 | 0.1205 | 0.0300 | 8 | -2.4027 | **0.0080** | 1000 |
| Miniopteridae | 222 | 0.0239 | 0.0254 | 0.0036 | 343 | -0.4042 | 0.3427 | 1000 |
| Pteropodidae | 2 | 0.0057 | 0.8612 | 0.4040 | 4.5 | -2.1177 | **0.0045** | 1000 |
| Rhinolophidae | 123 | 0.0175 | 0.0414 | 0.0067 | 1 | -3.5612 | **0.0010** | 1000 |
| Vespertilionidae | 174 | 0.0310 | 0.0312 | 0.0049 | 468 | -0.0575 | 0.4675 | 1000 |
| | Random subset | | | | | | | |
| | n | mntd.obs | mntd.rand.mean | mntd.rand.sd | mntd.obs.rank | mntd.obs.z | mntd.obs.p | runs |
| Hipposideridae | 29 | 0.0474 | 0.1346 | 0.0313 | 2 | -2.7889 | **0.0020** | 1000 |
| Miniopteridae | 39 | 0.0545 | 0.1065 | 0.0230 | 11 | -2.2656 | **0.0110** | 1000 |
| Pteropodidae | 2 | 0.0075 | 0.9117 | 0.3732 | 4 | -2.4229 | **0.0040** | 1000 |
| Rhinolophidae | 34 | 0.0602 | 0.1183 | 0.0275 | 13 | -2.1108 | **0.0130** | 1000 |
| Vespertilionidae | 30 | 0.1483 | 0.1319 | 0.0304 | 711 | 0.5411 | 0.7103 | 1000 |

**Supplementary Table 17**: Mean Nearest Taxon Distance (mntd.obs) and its standardized effect size (mntd.obs.z) within bat families for beta-CoVs using the complete dataset and random subset. One-tailed p-values (quantiles) were calculated after randomly reshuffling tip labels 1000 times along the entire phylogeny (=mpd.obs.rank/runs+1). Significant p-value (mntd.obs.p) are highlighted in bold.

| | Complete dataset | | | | | | | |
|---|---|---|---|---|---|---|---|---|
| | n | mntd.obs | mntd.rand.mean | mntd.rand.sd | mntd.obs.rank | mntd.obs.z | mntd.obs.p | runs |
| Hipposideridae | 21 | 0.0826 | 0.1145 | 0.0376 | 222 | -0.847 | 0.222 | 1000 |
| Pteropodidae | 58 | 0.0156 | 0.0424 | 0.0102 | 1 | -2.610 | **0.001** | 1000 |
| Rhinolophidae | 103 | 0.0119 | 0.0262 | 0.0043 | 1 | -3.344 | **0.001** | 1000 |
| Vespertilionidae | 117 | 0.0247 | 0.0233 | 0.0035 | 658 | 0.388 | 0.657 | 1000 |
| | Random subset | | | | | | | |
| | n | mntd.obs | mntd.rand.mean | mntd.rand.sd | mntd.obs.rank | mntd.obs.z | mntd.obs.p | runs |
| Hipposideridae | 21 | 0.0922 | 0.1179 | 0.0342 | 255 | -0.7498 | 0.2547 | 1000 |
| Pteropodidae | 33 | 0.0234 | 0.0689 | 0.0175 | 2 | -2.5961 | **0.0020** | 1000 |
| Rhinolophidae | 34 | 0.0246 | 0.0666 | 0.0162 | 1 | -2.5910 | **0.0010** | 1000 |
| Vespertilionidae | 39 | 0.0599 | 0.0574 | 0.0132 | 604 | 0.1888 | 0.6034 | 1000 |

**Supplementary Table 18**: Mean Phylogenetic Distance (mpd.obs) and its standardized effect size (mpd.obs.z) within bat genera for alpha-CoVs using the complete dataset and random subset. One-tailed p-values (quantiles) were calculated after randomly reshuffling tip labels 1000 times along the entire phylogeny (=mpd.obs.rank/runs+1). Significant p-value (mpd.obs.p) are highlighted in bold.

| | Complete dataset | | | | | | | |
|---|---|---|---|---|---|---|---|---|
| | n | mpd.obs | mpd.rand.mean | mpd.rand.sd | mpd.obs.rank | mpd.obs.z | mpd.obs.p | runs |
| *Eptesicus* | 2 | 0.0155 | 0.8360 | 0.4188 | 8 | -1.9591 | **0.0080** | 1000 |
| *Hipposideros* | 25 | 0.6517 | 0.8556 | 0.0618 | 3 | -3.2997 | **0.0030** | 1000 |
| *Hypsugo* | 3 | 0.0192 | 0.8580 | 0.2572 | 2 | -3.2614 | **0.0020** | 1000 |
| *Miniopterus* | 222 | 0.3332 | 0.8534 | 0.0154 | 1 | -33.8196 | **0.0010** | 1000 |
| *Murina* | 4 | 0.1323 | 0.8527 | 0.2033 | 11 | -3.5438 | **0.0110** | 1000 |
| *Myotis* | 130 | 0.5830 | 0.8534 | 0.0222 | 1 | -12.1821 | **0.0010** | 1000 |
| *Rhinolophus* | 123 | 0.7127 | 0.8521 | 0.0239 | 1 | -5.8310 | **0.0010** | 1000 |
| *Rousettus* | 2 | 0.0057 | 0.8369 | 0.4131 | 3 | -2.0122 | **0.0030** | 1000 |
| *Scotophilus* | 29 | 0.0457 | 0.8518 | 0.0550 | 1 | -14.6628 | **0.0010** | 1000 |
| *Vespertilio* | 3 | 0.0101 | 0.8534 | 0.2666 | 1 | -3.1629 | **0.0010** | 1000 |
| *Aselliscus* | 4 | 0.7467 | 0.8573 | 0.2034 | 281 | -0.5441 | 0.2807 | 1000 |
| *Tylonycteris* | 2 | 0.8257 | 0.8670 | 0.3970 | 263 | -0.1040 | 0.2627 | 1000 |
| *Nyctalus* | 1 | NA | NaN | NA | NA | NA | NA | 1000 |
| *Ia* | 1 | NA | NaN | NA | NA | NA | NA | 1000 |
| | Random subset | | | | | | | |
| | n | mpd.obs | mpd.rand.mean | mpd.rand.sd | mpd.obs.rank | mpd.obs.z | mpd.obs.p | runs |
| *Eptesicus* | 2 | 0.0176 | 0.9298 | 0.3730 | 14 | -2.4458 | **0.0140** | 1000 |
| *Hipposideros* | 25 | 0.6612 | 0.9194 | 0.0416 | 1 | -6.2046 | **0.0010** | 1000 |
| *Hypsugo* | 3 | 0.0211 | 0.9064 | 0.2276 | 1 | -3.8894 | **0.0010** | 1000 |
| *Miniopterus* | 39 | 0.3257 | 0.9198 | 0.0292 | 1 | -20.3600 | **0.0010** | 1000 |
| *Murina* | 4 | 0.1285 | 0.9252 | 0.1703 | 5 | -4.6778 | **0.0050** | 1000 |
| *Myotis* | 8 | 0.7143 | 0.9234 | 0.0927 | 26 | -2.2564 | **0.0260** | 1000 |
| *Rhinolophus* | 34 | 0.9206 | 0.9175 | 0.0323 | 482 | 0.0960 | 0.4815 | 1000 |
| *Rousettus* | 2 | 0.0075 | 0.9064 | 0.3793 | 2 | -2.3701 | **0.0020** | 1000 |
| *Scotophilus* | 6 | 0.0316 | 0.9176 | 0.1175 | 1 | -7.5397 | **0.0010** | 1000 |
| *Vespertilio* | 3 | 0.0184 | 0.9273 | 0.2322 | 2 | -3.9145 | **0.0020** | 1000 |
| *Aselliscus* | 4 | 0.7599 | 0.9189 | 0.1710 | 177 | -0.9295 | 0.1768 | 1000 |
| *Tylonycteris* | 2 | 0.8217 | 0.9508 | 0.3537 | 231.5 | -0.3651 | 0.2313 | 1000 |
| *Nyctalus* | 1 | NA | NaN | NA | NA | NA | NA | 1000 |
| *Ia* | 1 | NA | NaN | NA | NA | NA | NA | 1000 |

**Supplementary Table 19**: Mean Phylogenetic Distance (mpd.obs) and its standardized effect size (mpd.obs.z) within bat genera for beta-CoVs using the complete dataset and random subset. One-tailed p-values (quantiles) were calculated after randomly reshuffling tip labels 1000 times along the entire phylogeny (=mpd.obs.rank/runs+1). Significant p-value (mpd.obs.p) are highlighted in bold.

| | Complete dataset | | | | | | | |
|---|---|---|---|---|---|---|---|---|
| | n | mpd.obs | mpd.rand.mean | mpd.rand.sd | mpd.obs.rank | mpd.obs.z | mpd.obs.p | runs |
| *Cynopterus* | 9 | 0.0387 | 1.2272 | 0.1376 | 1 | -8.6357 | **0.0010** | 1000 |
| *Eonycteris* | 10 | 0.3887 | 1.2222 | 0.1305 | 1 | -6.3859 | **0.0010** | 1000 |
| *Eptesicus* | 5 | 0.1658 | 1.2498 | 0.2304 | 6 | -4.7052 | **0.0060** | 1000 |
| *Hipposideros* | 20 | 0.6994 | 1.2297 | 0.0704 | 1 | -7.5356 | **0.0010** | 1000 |
| *Hypsugo* | 5 | 0.0422 | 1.2328 | 0.2353 | 1 | -5.0602 | **0.0010** | 1000 |
| *Ia* | 2 | 0.0415 | 1.2711 | 0.6764 | 39 | -1.8179 | **0.0390** | 1000 |
| *Myotis* | 8 | 0.7057 | 1.2244 | 0.1535 | 7 | -3.3787 | **0.0070** | 1000 |
| *Pipistrellus* | 33 | 0.1848 | 1.2329 | 0.0446 | 1 | -23.5152 | **0.0010** | 1000 |

| | | | | | | | | |
|---|---|---|---|---|---|---|---|---|
| *Rhinolophus* | 103 | 0.1187 | 1.2299 | 0.0217 | 1 | -51.1789 | **0.0010** | 1000 |
| *Rousettus* | 42 | 0.3904 | 1.2304 | 0.0401 | 1 | -20.9568 | **0.0010** | 1000 |
| *Tylonycteris* | 43 | 0.0393 | 1.2295 | 0.0411 | 1 | -28.9869 | **0.0010** | 1000 |
| *Vespertilio* | 24 | 0.4223 | 1.2301 | 0.0632 | 1 | -12.7811 | **0.0010** | 1000 |
| *Aselliscus* | 1 | NA | NaN | NA | NA | NA | NA | 1000 |
| *Megaerops* | 1 | NA | NaN | NA | NA | NA | NA | 1000 |
| *Scotophilus* | 1 | NA | NaN | NA | NA | NA | NA | 1000 |
| | Random subset | | | | | | | |
| | n | mpd.obs | mpd.rand.mean | mpd.rand.sd | mpd.obs.rank | mpd.obs.z | mpd.obs.p | runs |
| *Cynopterus* | 7 | 0.0444 | 1.2603 | 0.1643 | 1 | -7.4009 | **0.0010** | 1000 |
| *Eonycteris* | 10 | 0.3830 | 1.2599 | 0.1084 | 1 | -8.0918 | **0.0010** | 1000 |
| *Eptesicus* | 5 | 0.1666 | 1.2609 | 0.2150 | 3 | -5.0891 | **0.0030** | 1000 |
| *Hipposideros* | 20 | 0.7198 | 1.2636 | 0.0644 | 1 | -8.4454 | **0.0010** | 1000 |
| *Hypsugo* | 5 | 0.0429 | 1.2568 | 0.2157 | 1 | -5.6279 | **0.0010** | 1000 |
| *Ia* | 2 | 0.0423 | 1.2299 | 0.6504 | 44.5 | -1.8258 | **0.0445** | 1000 |
| *Myotis* | 7 | 0.7409 | 1.2604 | 0.1551 | 8 | -3.3502 | **0.0080** | 1000 |
| *Pipistrellus* | 7 | 0.6448 | 1.2594 | 0.1680 | 10 | -3.6586 | **0.0100** | 1000 |
| *Rhinolophus* | 34 | 0.1233 | 1.2603 | 0.0414 | 1 | -27.4860 | **0.0010** | 1000 |
| *Rousettus* | 17 | 0.3935 | 1.2628 | 0.0750 | 1 | -11.5884 | **0.0010** | 1000 |
| *Tylonycteris* | 8 | 0.0427 | 1.2642 | 0.1369 | 1 | -8.9241 | **0.0010** | 1000 |
| *Vespertilio* | 7 | 0.7478 | 1.2580 | 0.1613 | 12 | -3.1631 | **0.0120** | 1000 |
| *Aselliscus* | 1 | NA | NaN | NA | NA | NA | NA | 1000 |
| *Megaerops* | 1 | NA | NaN | NA | NA | NA | NA | 1000 |
| *Scotophilus* | 1 | NA | NaN | NA | NA | NA | NA | 1000 |

**Supplementary Table 20**: Mean Nearest Taxon Distance (mntd.obs) and its standardized effect size (mntd.obs.z) within bat genera for alpha-CoVs using the complete dataset and random subset. One-tailed p-values (quantiles) were calculated after randomly reshuffling tip labels 1000 times along the entire phylogeny (=mpd.obs.rank/runs+1). Significant p-value (mntd.obs.p) are highlighted in bold.

| | Complete dataset | | | | | | | |
|---|---|---|---|---|---|---|---|---|
| | n | mntd.obs | mntd.rand.mean | mntd.rand.sd | mntd.obs.rank | mntd.obs.z | mntd.obs.p | runs |
| *Eptesicus* | 2 | 0.0155 | 0.8361 | 0.4293 | 11 | -1.9114 | **0.0110** | 1000 |
| *Hipposideros* | 25 | 0.0613 | 0.1355 | 0.0356 | 13 | -2.0856 | **0.0130** | 1000 |
| *Hypsugo* | 3 | 0.0101 | 0.6834 | 0.3046 | 1 | -2.2107 | **0.0010** | 1000 |
| *Miniopterus* | 222 | 0.0239 | 0.0253 | 0.0036 | 354 | -0.3752 | 0.3536 | 1000 |
| *Murina* | 4 | 0.0675 | 0.5582 | 0.2503 | 29 | -1.9598 | **0.0290** | 1000 |
| *Myotis* | 130 | 0.0312 | 0.0393 | 0.0070 | 127 | -1.1619 | 0.1269 | 1000 |
| *Rhinolophus* | 123 | 0.0175 | 0.0416 | 0.0071 | 1 | -3.3978 | **0.0010** | 1000 |
| *Rousettus* | 2 | 0.0057 | 0.8577 | 0.4033 | 3 | -2.1128 | **0.0030** | 1000 |
| *Scotophilus* | 29 | 0.0123 | 0.1194 | 0.0307 | 1 | -3.4858 | **0.0010** | 1000 |
| *Vespertilio* | 3 | 0.0067 | 0.6734 | 0.3079 | 1 | -2.1656 | **0.0010** | 1000 |
| *Aselliscus* | 4 | 0.1922 | 0.5786 | 0.2391 | 53 | -1.6160 | 0.0529 | 1000 |
| *Tylonycteris* | 2 | 0.8257 | 0.8529 | 0.3976 | 280.5 | -0.0683 | 0.2802 | 1000 |
| *Nyctalus* | 1 | NA | NaN | NA | NA | NA | NA | 1000 |
| *Ia* | 1 | NA | NaN | NA | NA | NA | NA | 1000 |
| | Random subset | | | | | | | |
| | n | mntd.obs | mntd.rand.mean | mntd.rand.sd | mntd.obs.rank | mntd.obs.z | mntd.obs.p | runs |
| *Eptesicus* | 2 | 0.0176 | 0.9167 | 0.3761 | 12 | -2.3905 | **0.0120** | 1000 |
| *Hipposideros* | 25 | 0.0627 | 0.1514 | 0.0379 | 8 | -2.3394 | **0.0080** | 1000 |
| *Hypsugo* | 3 | 0.0113 | 0.7619 | 0.2925 | 1 | -2.5657 | **0.0010** | 1000 |
| *Miniopterus* | 39 | 0.0545 | 0.1060 | 0.0219 | 13 | -2.3561 | **0.0130** | 1000 |
| *Murina* | 4 | 0.0660 | 0.6334 | 0.2445 | 12 | -2.3205 | **0.0120** | 1000 |
| *Myotis* | 8 | 0.4345 | 0.3939 | 0.1392 | 620 | 0.2914 | 0.6194 | 1000 |
| *Rhinolophus* | 34 | 0.0602 | 0.1195 | 0.0278 | 14 | -2.1340 | **0.0140** | 1000 |
| *Rousettus* | 2 | 0.0075 | 0.9389 | 0.3615 | 2 | -2.5764 | **0.0020** | 1000 |
| *Scotophilus* | 6 | 0.0107 | 0.4958 | 0.1836 | 1 | -2.6423 | **0.0010** | 1000 |
| *Vespertilio* | 3 | 0.0115 | 0.7442 | 0.2942 | 1 | -2.4905 | **0.0010** | 1000 |
| *Aselliscus* | 4 | 0.1951 | 0.6353 | 0.2489 | 39 | -1.7686 | **0.0390** | 1000 |
| *Tylonycteris* | 2 | 0.8217 | 0.9268 | 0.3632 | 231 | -0.2894 | 0.2308 | 1000 |
| *Nyctalus* | 1 | NA | NaN | NA | NA | NA | NA | 1000 |
| *Ia* | 1 | NA | NaN | NA | NA | NA | NA | 1000 |

**Supplementary Table 21**: Mean Nearest Taxon Distance (mntd.obs) and its standardized effect size (mntd.obs.z) within bat genera for beta-CoVs using the complete dataset and random subset. One-tailed p-values (quantiles) were calculated after randomly reshuffling tip labels 1000 times along the entire phylogeny (=mpd.obs.rank/runs+1). Significant p-value (mntd.obs.p) are highlighted in bold.

| | Complete dataset | | | | | | | |
|---|---|---|---|---|---|---|---|---|
| | n | mntd.obs | mntd.rand.mean | mntd.rand.sd | mntd.obs.rank | mntd.obs.z | mntd.obs.p | runs |
| *Cynopterus* | 9 | 0.0068 | 0.2779 | 0.1044 | 1 | -2.5964 | **0.0010** | 1000 |
| *Eonycteris* | 10 | 0.0201 | 0.2470 | 0.0910 | 1 | -2.4935 | **0.0010** | 1000 |
| *Eptesicus* | 5 | 0.0066 | 0.5392 | 0.2396 | 1 | -2.2232 | **0.0010** | 1000 |
| *Hipposideros* | 20 | 0.0883 | 0.1218 | 0.0390 | 209 | -0.8599 | 0.2088 | 1000 |
| *Hypsugo* | 5 | 0.0342 | 0.5407 | 0.2477 | 6 | -2.0445 | **0.0060** | 1000 |
| *Ia* | 2 | 0.0415 | 1.2035 | 0.6902 | 41 | -1.6835 | **0.0410** | 1000 |
| *Myotis* | 8 | 0.2677 | 0.3179 | 0.1275 | 351 | -0.3938 | 0.3506 | 1000 |
| *Pipistrellus* | 33 | 0.0909 | 0.0723 | 0.0215 | 803 | 0.8643 | 0.8022 | 1000 |

| | | | | | | | | |
|---|---|---|---|---|---|---|---|---|
| *Rhinolophus* | 103 | 0.0119 | 0.0261 | 0.0042 | 1 | -3.3579 | **0.0010** | 1000 |
| *Rousettus* | 42 | 0.0325 | 0.0565 | 0.0154 | 40 | -1.5572 | **0.0400** | 1000 |
| *Tylonycteris* | 43 | 0.0028 | 0.0559 | 0.0153 | 1 | -3.4829 | **0.0010** | 1000 |
| *Vespertilio* | 24 | 0.0047 | 0.1001 | 0.0321 | 1 | -2.9686 | **0.0010** | 1000 |
| *Aselliscus* | 1 | NA | NaN | NA | NA | NA | NA | 1000 |
| *Megaerops* | 1 | NA | NaN | NA | NA | NA | NA | 1000 |
| *Scotophilus* | 1 | NA | NaN | NA | NA | NA | NA | 1000 |
| | Random subset | | | | | | | |
| | n | mntd.obs | mntd.rand.mean | mntd.rand.sd | mntd.obs.rank | mntd.obs.z | mntd.obs.p | runs |
| *Cynopterus* | 7 | 0.0093 | 0.4256 | 0.1495 | 1 | -2.7847 | **0.0010** | 1000 |
| *Eonycteris* | 10 | 0.0189 | 0.2808 | 0.0911 | 1 | -2.8739 | **0.0010** | 1000 |
| *Eptesicus* | 5 | 0.0084 | 0.6073 | 0.2377 | 2 | -2.5197 | **0.0020** | 1000 |
| *Hipposideros* | 20 | 0.0986 | 0.1244 | 0.0349 | 239 | -0.7397 | 0.2388 | 1000 |
| *Hypsugo* | 5 | 0.0356 | 0.6098 | 0.2502 | 1 | -2.2954 | **0.0010** | 1000 |
| *Ia* | 2 | 0.0423 | 1.3030 | 0.6178 | 33 | -2.0405 | **0.0330** | 1000 |
| *Myotis* | 7 | 0.3056 | 0.4231 | 0.1559 | 243 | -0.7536 | 0.2428 | 1000 |
| *Pipistrellus* | 7 | 0.3442 | 0.4248 | 0.1514 | 301 | -0.5324 | 0.3007 | 1000 |
| *Rhinolophus* | 34 | 0.0246 | 0.0669 | 0.0161 | 2 | -2.6303 | **0.0020** | 1000 |
| *Rousettus* | 17 | 0.0674 | 0.1504 | 0.0432 | 19 | -1.9231 | **0.0190** | 1000 |
| *Tylonycteris* | 8 | 0.0078 | 0.3623 | 0.1238 | 1 | -2.8652 | **0.0010** | 1000 |
| *Vespertilio* | 7 | 0.0081 | 0.4180 | 0.1494 | 1 | -2.7430 | **0.0010** | 1000 |
| *Aselliscus* | 1 | NA | NaN | NA | NA | NA | NA | 1000 |
| *Megaerops* | 1 | NA | NaN | NA | NA | NA | NA | 1000 |
| *Scotophilus* | 1 | NA | NaN | NA | NA | NA | NA | 1000 |

**Supplementary Table 22**: Mean Phylogenetic Distance (mpd.obs) and its standardized effect size (mpd.obs.z) within zoogeographic regions for alpha-CoVs using the complete dataset and random subset. One-tailed p-values (quantiles) were calculated after randomly reshuffling tip labels 1000 times along the entire phylogeny (=mpd.obs.rank/runs+1). Significant p-value (mpd.obs.p) are highlighted in bold. NO, Northern region; CN, Central northern region; SW, South western region; CE, Central region; SO, Southern region; HI, Hainan island.

| | Complete dataset | | | | | | | |
|---|---|---|---|---|---|---|---|---|
| | n | mpd.obs | mpd.rand.mean | mpd.rand.sd | mpd.obs.rank | mpd.obs.z | mpd.obs.p | runs |
| CE | 33 | 0.6571 | 0.8088 | 0.0531 | 13 | -2.8550 | **0.0130** | 1000 |
| CN | 26 | 0.4569 | 0.8073 | 0.0602 | 1 | -5.8252 | **0.0010** | 1000 |
| HI | 36 | 0.7413 | 0.8063 | 0.0515 | 125 | -1.2631 | 0.1249 | 1000 |
| NO | 61 | 0.7342 | 0.8051 | 0.0367 | 44 | -1.9351 | **0.0440** | 1000 |
| SO | 195 | 0.7418 | 0.8048 | 0.0169 | 1 | -3.7391 | **0.0010** | 1000 |
| SW | 161 | 0.8226 | 0.8047 | 0.0201 | 811 | 0.8898 | 0.8102 | 1000 |
| | Random subset | | | | | | | |
| | n | mpd.obs | mpd.rand.mean | mpd.rand.sd | mpd.obs.rank | mpd.obs.z | mpd.obs.p | runs |
| CE | 33 | 0.6753 | 0.8627 | 0.0444 | 1 | -4.2172 | **0.0010** | 1000 |
| CN | 26 | 0.4747 | 0.8603 | 0.0540 | 1 | -7.1421 | **0.0010** | 1000 |
| HI | 36 | 0.7807 | 0.8602 | 0.0428 | 38 | -1.8555 | **0.0380** | 1000 |
| NO | 42 | 0.7455 | 0.8625 | 0.0374 | 2 | -3.1302 | **0.0020** | 1000 |
| SO | 48 | 0.7953 | 0.8612 | 0.0360 | 46 | -1.8293 | **0.0460** | 1000 |
| SW | 47 | 0.9428 | 0.8622 | 0.0367 | 1000 | 2.1976 | **0.9990** | 1000 |

**Supplementary Table 23**: Mean Phylogenetic Distance (mpd.obs) and its standardized effect size (mpd.obs.z) within zoogeographic regions for beta-CoVs using the complete dataset and random subset. One-tailed p-values (quantiles) were calculated after randomly reshuffling tip labels 1000 times along the entire phylogeny (=mpd.obs.rank/runs+1). Significant p-value (mpd.obs.p) are highlighted in bold. NO, Northern region; CN, Central northern region; SW, South western region; CE, Central region; SO, Southern region.

| | Complete dataset | | | | | | | |
|---|---|---|---|---|---|---|---|---|
| | n | mpd.obs | mpd.rand.mean | mpd.rand.sd | mpd.obs.rank | mpd.obs.z | mpd.obs.p | runs |
| CE | 53 | 0.9032 | 1.2591 | 0.0372 | 1 | -9.5621 | **0.0010** | 1000 |
| CN | 40 | 1.0626 | 1.2571 | 0.0462 | 1 | -4.2134 | **0.0010** | 1000 |
| NO | 23 | 0.8428 | 1.2543 | 0.0688 | 1 | -5.9774 | **0.0010** | 1000 |
| SO | 103 | 1.1273 | 1.2571 | 0.0223 | 1 | -5.8184 | **0.0010** | 1000 |
| SW | 67 | 1.0388 | 1.2577 | 0.0330 | 1 | -6.6241 | **0.0010** | 1000 |
| | Random subset | | | | | | | |
| | n | mpd.obs | mpd.rand.mean | mpd.rand.sd | mpd.obs.rank | mpd.obs.z | mpd.obs.p | runs |
| CE | 30 | 0.6653 | 1.2250 | 0.0666 | 1 | -8.4043 | **0.0010** | 1000 |
| CN | 25 | 0.9666 | 1.2230 | 0.0760 | 5 | -3.3733 | **0.0050** | 1000 |
| NO | 23 | 0.7454 | 1.2213 | 0.0794 | 1 | -5.9950 | **0.0010** | 1000 |
| SO | 30 | 1.0708 | 1.2238 | 0.0673 | 25 | -2.2720 | **0.0250** | 1000 |
| SW | 37 | 1.0042 | 1.2179 | 0.0591 | 3 | -3.6137 | **0.0030** | 1000 |

**Supplementary Table 24**: Mean Nearest Taxon Distance (mntd.obs) and its standardized effect size (mntd.obs.z) within zoogeographic regions for alpha-CoVs using the complete dataset and random subset. One-tailed p-values (quantiles) were calculated after randomly reshuffling tip labels 1000 times along the entire phylogeny (=mpd.obs.rank/runs+1). Significant p-value (mntd.obs.p) are highlighted in bold. NO, Northern region; CN, Central northern region; SW, South western region; CE, Central region; SO, Southern region; HI, Hainan island.

| | Complete dataset | | | | | | | |
|---|---|---|---|---|---|---|---|---|
| | ntaxa | mntd.obs | mntd.rand.mean | mntd.rand.sd | mntd.obs.rank | mntd.obs.z | mntd.obs.p | runs |
| CE | 33 | 0.0583 | 0.1070 | 0.0274 | 19 | -1.7747 | **0.0190** | 1000 |
| CN | 26 | 0.1045 | 0.1274 | 0.0342 | 252 | -0.6718 | 0.2517 | 1000 |
| HI | 36 | 0.0415 | 0.1003 | 0.0246 | 5 | -2.3947 | **0.0050** | 1000 |
| NO | 61 | 0.0584 | 0.0691 | 0.0151 | 248 | -0.7085 | 0.2478 | 1000 |
| SO | 195 | 0.0119 | 0.0300 | 0.0050 | 1 | -3.6427 | **0.0010** | 1000 |
| SW | 161 | 0.0200 | 0.0349 | 0.0061 | 5 | -2.4495 | **0.0050** | 1000 |
| | Random subset | | | | | | | |
| | n | mntd.obs | mntd.rand.mean | mntd.rand.sd | mntd.obs.rank | mntd.obs.z | mntd.obs.p | runs |
| CE | 33 | 0.0601 | 0.1013 | 0.0256 | 43 | -1.6079 | **0.0430** | 1000 |
| CN | 26 | 0.1179 | 0.1249 | 0.0322 | 423 | -0.2185 | 0.4226 | 1000 |
| HI | 36 | 0.0482 | 0.0951 | 0.0225 | 15 | -2.0847 | **0.0150** | 1000 |
| NO | 42 | 0.0627 | 0.0843 | 0.0196 | 135 | -1.0965 | 0.1349 | 1000 |
| SO | 48 | 0.0451 | 0.0769 | 0.0168 | 21 | -1.8985 | **0.0210** | 1000 |
| SW | 47 | 0.0679 | 0.0775 | 0.0174 | 311 | -0.5539 | 0.3107 | 1000 |

**Supplementary Table 25**: Mean Nearest Taxon Distance (mntd.obs) and its standardized effect size (mntd.obs.z) within zoogeographic regions for beta-CoVs using the complete dataset and random subset. One-tailed p-values (quantiles) were calculated after randomly reshuffling tip labels 1000 times along the entire phylogeny (=mpd.obs.rank/runs+1). Significant p-value (mntd.obs.p) are highlighted in bold. NO, Northern region; CN, Central northern region; SW, South western region; CE, Central region; SO, Southern region.

| | Complete dataset | | | | | | | |
|---|---|---|---|---|---|---|---|---|
| | n | mntd.obs | mntd.rand.mean | mntd.rand.sd | mntd.obs.rank | mntd.obs.z | mntd.obs.p | runs |
| CE | 53 | 0.0413 | 0.0460 | 0.0119 | 385 | -0.3900 | 0.3846 | 1000 |
| CN | 40 | 0.0162 | 0.0615 | 0.0175 | 1 | -2.5923 | **0.0010** | 1000 |
| NO | 23 | 0.0097 | 0.1081 | 0.0347 | 1 | -2.8372 | **0.0010** | 1000 |
| SO | 103 | 0.0198 | 0.0258 | 0.0041 | 73 | -1.4606 | 0.0729 | 1000 |
| SW | 67 | 0.0151 | 0.0368 | 0.0083 | 1 | -2.6244 | **0.0010** | 1000 |
| | Random subset | | | | | | | |
| | n | mntd.obs | mntd.rand.mean | mntd.rand.sd | mntd.obs.rank | mntd.obs.z | mntd.obs.p | runs |
| CE | 30 | 0.0803 | 0.0752 | 0.0239 | 606 | 0.2105 | 0.6054 | 1000 |
| CN | 25 | 0.0077 | 0.0899 | 0.0290 | 1 | -2.8348 | **0.0010** | 1000 |
| NO | 23 | 0.0094 | 0.0983 | 0.0310 | 1 | -2.8703 | **0.0010** | 1000 |
| SO | 30 | 0.0360 | 0.0748 | 0.0229 | 35 | -1.6969 | **0.0350** | 1000 |
| SW | 37 | 0.0658 | 0.0592 | 0.0177 | 662 | 0.3720 | 0.6613 | 1000 |

**Supplementary Table 26**: R square and p-value of linear regression analysis between Alpha-CoV phylogenetic diversity (MPD) and bat species richness (total or sampled) per zoogeographic region or province in China

| | Complete dataset | |
|---|---|---|
| Linear regression | R square | p-value |
| Total bat richness per zoogeographic region | 0.033 | 0.728 |
| Sampled bat richness per zoogeographic region | 0.199 | 0.374 |
| Total bat richness per province | 0.119 | 0.191 |
| Sampled bat richness per province | 0.018 | 0.621 |
| | Random subset | |
| Linear regression | R square | p-value |
| Total bat richness per zoogeographic region | 0.075 | 0.599 |
| Sampled bat richness per zoogeographic region | 0.209 | 0.362 |
| Total bat richness per province | 0.203 | 0.080 |
| Sampled bat richness per province | 0.073 | 0.311 |

**Supplementary Table 27**: R square and p-value of linear regression analysis between Beta-CoV phylogenetic diversity (MPD) and bat species richness (total or sampled) per zoogeographic region or province in China

| | Complete dataset | |
|---|---|---|
| Linear regression | R square | p-value |
| Total bat richness per zoogeographic region | 0.090 | 0.624 |
| Sampled bat richness per zoogeographic region | 0.440 | 0.222 |
| Total bat richness per province | 0.128 | 0.174 |
| Sampled bat richness per province | 0.331 | **0.020** |
| | Random subset | |
| Linear regression | R square | p-value |
| Total bat richness per zoogeographic region | 0.011 | 0.866 |
| Sampled bat richness per zoogeographic region | 0.352 | 0.292 |
| Total bat richness per province | 0.101 | 0.231 |
| Sampled bat richness per province | 0.352 | **0.015** |

**Supplementary Table 28**: Phylogenetic β-diversity (standardized effect size (SES) of the Mean Phylogenetic Distance (MPD)) among bat families for alpha-CoVs using the complete dataset (below the diagonal) and random subset (above the diagonal).

| | Hipposideridae | Rhinolophidae | Vespertilionidae | Miniopteridae | Pteropodidae |
|---|---|---|---|---|---|
| Hipposideridae | | 20.1333 | -4.5575 | 1.1073 | 4.0884 |
| Rhinolophidae | 7.1505 | | 7.8328 | 10.0433 | 0.0297 |
| Vespertilionidae | 8.4421 | 7.0519 | | 3.0433 | 3.6763 |
| Miniopteridae | 15.5949 | 20.3070 | 13.8885 | | -3.4863 |
| Pteropodidae | -2.6837 | 4.4666 | 4.0478 | 5.3109 | |

**Supplementary Table 29**: Phylogenetic β-diversity (standardized effect size (SES) of the Mean Phylogenetic Distance (MPD)) among bat families for Beta-CoVs using the complete dataset (below the diagonal) and random subset (above the diagonal).

| | Hipposideridae | Rhinolophidae | Vespertilionidae | Pteropodidae |
|---|---|---|---|---|
| Hipposideridae | | 7.6102 | -12.614 | 3.0319 |
| Rhinolophidae | -11.9738 | | 15.5243 | 15.8607 |
| Vespertilionidae | 5.2859 | 24.4489 | | 8.8575 |
| Pteropodidae | 5.9384 | 18.2126 | 27.8056 | |

**Supplementary Table 30**: Phylogenetic β-diversity (standardized effect size (SES) of the Mean Phylogenetic Distance (MPD)) among bat genera for alpha-CoVs using the complete dataset (below the diagonal) and random subset (above the diagonal).

| | *Aselliscus* | *Hipposideros* | *Rhinolophus* | *Eptesicus* | *Hypsugo* | *Ia* | *Murina* | *Myotis* | *Nyctalus* | *Scotophilus* | *Tylonycteris* | *Vespertilio* | *Miniopterus* | *Rousettus* |
|---|---|---|---|---|---|---|---|---|---|---|---|---|---|---|
| *Aselliscus* | | -2.800 | -0.509 | 0.288 | 0.948 | -0.775 | -1.288 | -2.951 | -0.565 | -1.171 | 0.736 | -3.34 | 4.102 | -1.171 |
| *Hipposideros* | -1.101 | | 1.446 | 2.755 | 4.408 | -0.044 | 0.723 | 1.609 | 0.526 | 2.528 | 3.645 | 0.634 | 20.23 | -4.814 |
| *Rhinolophus* | 1.638 | 6.981 | | 0.843 | 2.235 | -1.99 | -2.113 | -3.783 | -0.959 | -1.778 | 1.322 | -2.145 | 10.110 | 3.036 |
| *Eptesicus* | 0.479 | 2.554 | 1.655 | | -4.702 | -0.592 | -1.018 | -1.041 | -1.739 | -0.581 | -0.4 | -0.876 | 1.185 | 1.877 |
| *Hypsugo* | 1.046 | 3.799 | 2.946 | -3.818 | | -0.418 | -0.611 | -0.421 | -1.784 | 0.030 | -0.027 | -0.529 | 2.872 | 2.769 |
| *Ia* | -0.396 | 0.299 | 0.062 | -0.258 | -0.039 | | -1.765 | -2.244 | -0.890 | -1.63 | -0.388 | -1.518 | -0.268 | 0.634 |
| *Murina* | -0.364 | 2.212 | -0.705 | -0.238 | 0.238 | -0.623 | | -3.554 | -1.573 | -2.870 | -0.552 | -2.75 | -0.145 | 1.414 |
| *Myotis* | -2.075 | 8.289 | 4.289 | 0.012 | 0.99 | -1.727 | -2.202 | | -1.857 | -3.894 | -0.602 | -6.332 | -0.242 | 2.339 |
| *Nyctalus* | -0.296 | 0.826 | -0.009 | -1.554 | -1.554 | -0.603 | -0.858 | -1.205 | | -1.401 | -0.912 | -1.375 | -0.910 | 0.819 |
| *Scotophilus* | 0.957 | 10.902 | 7.918 | 1.590 | 2.728 | 0.445 | -0.401 | -3.158 | 0.088 | | -0.020 | -3.355 | 1.144 | 2.285 |
| *Tylonycteris* | 0.769 | 2.967 | 1.724 | -0.408 | -0.131 | -0.099 | 0.086 | 0.248 | -0.872 | 2.056 | | -0.476 | -5.056 | 2.298 |
| *Vespertilio* | -2.293 | 1.328 | -1.047 | -0.593 | -0.252 | -0.852 | -2.276 | -6.619 | -1.068 | -3.216 | -0.334 | | -0.131 | 1.259 |
| *Miniopterus* | 3.069 | 15.72 | 20.662 | 0.728 | 1.778 | 0.43 | 1.312 | 7.774 | -0.622 | 10.820 | -3.077 | 0.07 | | 7.978 |
| *Rousettus* | -0.601 | -2.944 | 4.751 | 1.869 | 2.565 | 0.549 | 2.215 | 4.124 | 0.917 | 5.396 | 2.155 | 1.532 | 5.732 | |

**Supplementary Table 31**: Phylogenetic β-diversity (standardized effect size (SES) of the Mean Phylogenetic Distance (MPD)) among bat genera for beta-CoVs using the complete dataset (below the diagonal) and random subset (above the diagonal).

| | *Aselliscus* | *Hipposideros* | *Rhinolophus* | *Eptesicus* | *Hypsugo* | *Ia* | *Myotis* | *Pipistrellus* | *Scotophilus* | *Tylonycteris* | *Vespertilio* | *Cynopterus* | *Eonycteris* | *Megaerops* | *Rousettus* |
|---|---|---|---|---|---|---|---|---|---|---|---|---|---|---|---|
| *Aselliscus* | | -2.172 | -5.361 | 0.576 | 0.666 | 0.254 | 0.917 | 1.073 | 0.868 | 2.011 | 0.001 | 0.851 | 2.489 | 1.393 | 2.371 |
| *Hipposideros* | -2.064 | | -11.697 | 0.524 | 0.855 | -0.001 | 1.376 | 1.937 | 1.792 | 5.119 | -2.577 | 0.376 | 5.859 | 2.969 | 5.995 |
| *Rhinolophus* | -4.736 | -11.642 | | 1.918 | 2.334 | 0.823 | 3.401 | 4.119 | 2.745 | 8.087 | -0.253 | 2.805 | 10.023 | 4.514 | 11.32 |
| *Eptesicus* | 0.702 | 1.176 | 2.491 | | -5.272 | -4.151 | -4.692 | -4.486 | -1.645 | -3.921 | -4.900 | 0.993 | 4.437 | 2.401 | 4.356 |
| *Hypsugo* | 0.831 | 1.588 | 3.092 | -4.464 | | -3.526 | -4.943 | -4.795 | -1.987 | -4.795 | -4.552 | 1.211 | 4.618 | 2.488 | 4.671 |
| *Ia* | 0.401 | 0.54 | 1.314 | -3.586 | -2.922 | | -3.677 | -2.756 | -1.109 | -2.566 | -3.512 | 0.387 | 2.622 | 1.572 | 2.502 |
| *Myotis* | 1.033 | 2.206 | 4.361 | -3.906 | -4.137 | -3.037 | | -4.043 | -1.914 | -4.639 | -4.041 | 0.355 | 4.371 | 2.506 | 4.313 |
| *Pipistrellus* | 1.597 | 5.021 | 11.489 | -5.82 | -6.718 | -3.602 | -5.913 | | -1.803 | -4.393 | -3.617 | 0.754 | 4.898 | 2.712 | 4.860 |
| *Scotophilus* | 0.93 | 1.966 | 2.852 | -1.372 | -1.669 | -0.920 | -1.744 | -2.495 | | -4.247 | -1.064 | 1.785 | 3.555 | 1.836 | 3.695 |
| *Tylonycteris* | 2.593 | 9.336 | 20.573 | -5.027 | -6.129 | -3.055 | -7.00 | -14.206 | -4.906 | | -2.563 | 4.477 | 9.210 | 4.307 | 10.176 |
| *Vespertilio* | -0.364 | -3.713 | -2.526 | -6.447 | -5.95 | -4.477 | -5.889 | -9.562 | -1.434 | -7.181 | | 0.280 | 4.144 | 2.386 | 3.944 |
| *Cynopterus* | 0.378 | -0.659 | 1.227 | 1.250 | 1.562 | 0.631 | 1.004 | 3.47 | 1.821 | 6.808 | -1.505 | | -4.091 | -0.688 | -6.017 |
| *Eonycteris* | 1.838 | 4.353 | 7.791 | 4.368 | 4.646 | 2.724 | 4.938 | 9.092 | 3.355 | 13.055 | 3.582 | -3.698 | | -3.456 | -10.8 |
| *Megaerops* | 1.119 | 2.386 | 3.624 | 2.3388 | 2.431 | 1.557 | 2.565 | 3.991 | 1.782 | 5.519 | 2.201 | -0.654 | -2.913 | | -3.464 |
| *Rousettus* | 2.172 | 6.571 | 16.731 | 5.735 | 6.327 | 3.449 | 6.756 | 15.928 | 4.298 | 23.882 | 5.498 | -5.729 | -10.765 | -3.676 | |

**Supplementary Table 32**: Phylogenetic β-diversity (standardized effect size (SES) of the Mean Phylogenetic Distance (MPD)) among zoogeographic regions for Alpha-CoVs using the complete dataset (below the diagonal) and random subset (above the diagonal). NO, Northern region; CN, Central northern region; SW, South western region; CE, Central region; SO, Southern region; HI, Hainan island.

| | CE | CN | NO | SO | SW | HI |
|---|---|---|---|---|---|---|
| CE | | -0.4238 | 2.9153 | -4.8786 | -2.7423 | 4.4078 |
| CN | 0.9041 | | -1.2683 | -5.5677 | -1.4736 | 0.4568 |
| NO | -1.7163 | -2.8428 | | 1.7891 | 0.943 | 4.8338 |
| SO | -2.8927 | -1.2139 | -1.7968 | | -2.0649 | 3.5021 |
| SW | 0.6287 | 2.4100 | 2.0902 | 0.6691 | | 4.7082 |
| HI | 3.7697 | 1.2204 | 3.5429 | 2.6766 | 6.1801 | |

**Supplementary Table 33**: Phylogenetic β-diversity (standardized effect size (SES) of the Mean Phylogenetic Distance (MPD)) among zoogeographic regions for Beta-CoVs using the complete dataset (below the diagonal) and random subset (above the diagonal). NO, Northern region; CN, Central northern region; SW, South western region; CE, Central region; SO, Southern region.

| | CE | CN | NO | SO | SW |
|---|---|---|---|---|---|
| CE | | -2.3751 | -9.0517 | 0.9142 | 6.3531 |
| CN | -5.2248 | | -3.6026 | -1.3420 | 6.8906 |
| NO | -6.7161 | -5.686 | | 0.0850 | 6.2733 |
| SO | -1.6547 | -5.4274 | -1.2255 | | 8.6052 |
| SW | 7.0965 | 3.6185 | 3.4593 | 16.3362 | |

**Supplementary Table 34**: GenBank accession numbers of Alpha-CoV sequences

| | Lineages | | | | | | |
|---|---|---|---|---|---|---|---|
| Hosts | L1 | L2 | L3 | L4 | L5 | L6 | L7 |
| **Pteropodidae** | | | | | | | |
| *Rousettus leschenaultii* | | JQ989270-271 | | | | | |
| **Hipposideridae** | | | | | | | |
| *Aselliscus stoliczkanus* | | MN312307<br>MN312477 | | | MN312268-269 | | |
| *Hipposideros armiger* | MN312240<br>MN312251 | MN312473<br>MN312475 | KY770852 | | | | |
| *Hipposideros cineraceus* | KU182955 | | | | | | |
| *Hipposideros pomona* | MF769513 | MN312250<br>MN312290<br>MN312308<br>MN312310-312<br>MN312314-315<br>MN312406-408<br>MN312455<br>MN312581<br>JQ989266-269<br>JQ989272-273<br>KP895522-523<br>KU343195-196 | | | | | |
| *Hipposideros pratti* | KX285186 | | | | | | |
| **Rhinolophidae** | | | | | | | |
| *Rhinolophus affinis* | KX285119-120<br>MF769453<br>MF769468-470<br>MF769485<br>MN312247<br>MN312261<br>MN312267<br>MN312273<br>MN312276<br>MN312289<br>MN312303<br>MN312318<br>MN312320-321<br>MN312323<br>MN312325-329<br>MN312331-334<br>MN312336<br>MN312338<br>MN312341<br>MN312344-349<br>MN312351-352<br>MN312354-355<br>MN312359<br>MN312454<br>MN312456<br>MN312458<br>MN312465<br>MN312476<br>MN312479<br>MN312561 | MN312356<br>MN312474<br>MN312481-482<br>KP876527 | | | | | KP876544 |

| | | | | | | | |
|---|---|---|---|---|---|---|---|
| | MN312576-580<br>MN312582-583<br>MN312587-588<br>KP876528<br>KU343198 | | | | | | |
| *Rhinolophus ferrumequinum* | KJ473808 | MN312417<br>KF569979-981<br>KJ473807<br>KX285155 | | | | | |
| *Rhinolophus hipposideros* | | KF569982 | | | | | |
| *Rhinolophus macrotis* | MN312411 | | KY770854 | | | | |
| *Rhinolophus monoceros* | | | | | KT381913<br>KT381915-916 | | |
| *Rhinolophus pearsonii* | | | KY770853 | | | | KF294272 |
| *Rhinolophus pusillus* | MF760455<br>MF760515<br>MF769475-476<br>MN312516<br>MN312586 | | KY770855 | | KY009617 | | KY783865<br>KY783896 |
| *Rhinolophus rex* | KX285187-193 | | | | | | |
| *Rhinolophus sinicus* | MF769447-451<br>MF769466-467<br>MF769477-478<br>MF769508-509<br>MF769511<br>MN312291<br>MN312293-294<br>MN312449-450<br>MN312478<br>MN312515<br>EF203064-067<br>KP876537<br>KP876540-541<br>KU182954 | KP876532<br>KP876536<br>KU182967-968<br>KU182970-971 | KX285168<br>KX285169 | | | | MN312405<br>MN312451<br>MN312590<br>KP876509<br>KP876525-526<br>KP876533<br>KP876545 |
| *Rhinolophus stheno* | KP895525 | | | | | | |
| *Rhinolophus* sp. | MN312241-242<br>MN312246<br>MN312249<br>MN312252<br>MN312254<br>MN312256-257<br>MN312260<br>MN312263<br>MN312265<br>MN312270-272<br>MN312274-275<br>MN312277-278<br>MN312301<br>MN312319<br>MN312322<br>MN312324<br>MN312337<br>MN312339-340<br>MN312350<br>MN312357<br>MN312457<br>MN312461-462 | MN312330<br>MN312463<br>MN312472 | | | MN312258 | | MN312253 |

| | | | | | | | |
|---|---|---|---|---|---|---|---|
| | MN312480<br>MN312243<br>MN312245<br>MN312264<br>MN312292<br>MN312299<br>MN312335 | | | | | | |
| **Miniopteridae** | | | | | | | |
| *Miniopterus schreibersii* | MN312286<br>MN312466 | | KX285179 | KF294378 | KX285171<br>KX285176<br>MN312445<br>KT381919-923<br>KX285176<br>KF294383 | | KX285115-117<br>KX285121-124<br>KX285170<br>KX285172-175<br>KX285177-178<br>MN312248<br>MN312279-280<br>MN312283-284<br>MN312287-288<br>MN312302<br>MN312372-373<br>MN312378<br>MN312380<br>MN312382-387<br>MN312389<br>MN312394<br>MN312397-400<br>MN312402<br>MN312404<br>MN312421-422<br>MN312507<br>MN312592<br>KY783870-872<br>KY783874<br>KY783876-878<br>KY783880<br>MN312452-453<br>MN312483-484<br>MN312486-487<br>MN312491<br>MN312493-498<br>MN312504<br>MN312511-512<br>MN312537<br>MN312547<br>MN312550-551<br>MN312559-560<br>MN312564-565<br>KJ473795-804<br>KP876505-506<br>KP876512-515<br>KT381917-918<br>KU343190<br>KX285118<br>KX285171-175<br>DQ648835<br>DQ648850<br>KF294268-271<br>KF294273-282<br>KP876507-508<br>KP876510<br>KP876517-518<br>KP876521<br>KU343191<br>KU343194<br>KY783887-888<br>KY783890-894 |

| | | | | | | | |
|---|---|---|---|---|---|---|---|
| | | | | | | | KY783897-901<br>KY783903 |
| *Miniopterus fuscus* | | KP876534 | | | | | MN312295-297<br>MN312300<br>MN312305<br>MN312313<br>MN312566-568<br>KU343192-193 |
| *Miniopterus magnater* | | | | | | | DQ666337<br>DQ666339-340<br>EU420138 |
| *Miniopterus pusillus* | | | | | | | MN312365-366<br>MN312377<br>MN312379<br>MN312381<br>MN312388<br>MN312390-392<br>MN312401<br>MN312403<br>MN312435-439<br>MN312485<br>MN312488-490<br>MN312492<br>MN312499-502<br>MN312505-506<br>MN312508-510<br>MN312513<br>MN312548<br>DQ666338<br>EU420137 |
| *Miniopterus* sp. | | | | | MN312446 | | KX285129-133<br>MN312262<br>MN312281-282<br>MN312285<br>MN312298<br>MN312306<br>MN312317<br>MN312342-343<br>MN312353<br>MN312358<br>MN312360-362<br>MN312460<br>MN312468<br>MN312470<br>KY783873<br>KY783875<br>KY783879<br>MN312304<br>MN312309<br>MN312467<br>MN312469<br>MN312503<br>MN312562-563<br>KP876519-520<br>KP876522-524<br>KP876542 |
| **Vespertilionidae** | | | | | | | |
| *Eptesicus serotinus* | | | | | | KY009627<br>KY009631 | |
| *Hypsugo* sp. | | | | | | KX285210-213<br>MN312569-575 | |
| *Ia io* | | | KY770857 | | | | |
| *Murina* | | | | KF294373-377<br>KU182966 | | | |

| | | | | | | | |
|---|---|---|---|---|---|---|---|
| *leucogaster* | | | | | | | |
| *Myotis chinensis* | | | MN312412 | | MN312375<br>MN312523-524<br>MN312530<br>MN312532-533 | | |
| *Myotis daubentonii* | KP895492 | | KF569975<br>KF569995<br>KP895499<br>KP895508<br>KP895510 | | KF569974<br>KF569976-978<br>KF569994<br>KP895494<br>KP895496<br>KP895498<br>KP895500-507<br>KP895509<br>KP895511-521 | | DQ648855<br>KP895491<br>KP895493 |
| *Myotis davidii* | | | KX285201-206<br>KF294381<br>KF569991<br>KY770851<br>KY770856 | | KF294382<br>KF569983-988<br>KF569990<br>KF569992-993 | | KF569989 |
| *Myotis fimbriatus* | | | | | KY009612<br>KY009620<br>KY009623<br>KY009629<br>KY009630<br>KY009633 | | |
| *Myotis horsfieldii* | | | | | | | MN312433 |
| *Myotis myotis* | | | KX285183 | | | | |
| *Myotis ricketti* | MF769510 | | KX285137<br>KX285141-142<br>KX285126<br>KX285128<br>KX285147-149<br>KX285151-152<br>KX285154<br>MN312415<br>KJ473806<br>KX285137-140<br>KX285142-154 | | KX285215-216<br>MN312367<br>MN312370<br>MN312410<br>MN312416<br>KX285127<br>KX285214<br>KX285217-218<br>KY383882<br>KY783866-869<br>KY783881<br>KY783883-885<br>MN312363<br>MN312368-369<br>MN312371<br>MN312374<br>MN312376<br>MN312393<br>MN312395-396<br>MN312409<br>MN312414<br>MN312418<br>MN312440<br>MN312442-444<br>MN312448<br>MN312520-522<br>MN312525-528<br>MN312531<br>MN312534-536<br>MN312543-546<br>MN312549<br>DQ648833<br>KP895490<br>KY009616<br>KY009625 | | KX285143<br>MN312364<br>MN312447<br>KX285141 |

| | | | | | | | |
|---|---|---|---|---|---|---|---|
| *Myotis siligorensis* | | | KY770850 | | | | |
| *Myotis* sp. | | KJ473810 | KX285138-139<br>KX285144-146<br>KX285153 | | KX285194<br>MN312413<br>MN312514 | | |
| *Nyctalus plancyi* | | | | | | KJ473809 | |
| *Scotophilus kuhlii* | | | | | MN312419-420<br>MN312423-432<br>MN312434<br>MN312538-542<br>MN312552-558<br>MN312584-585<br>MN312589<br>MN312593-594<br>DQ648822-823<br>DQ648858<br>KT381902-912 | | |
| *Tylonycteris pachypus* | | | | | | | MN312591 |
| *Tylonycteris robustula* | | | | | | KX447559 | |
| *Vespertilio sinensis* | | | | | MN312517-519 | | |

**Supplementary Table 35:** GenBank accession numbers of Beta-CoV sequences

| | Lineages | | | |
|---|---|---|---|---|
| Hosts | B | C | D | E |
| **Pteropodidae** | | | | |
| *Cynopterus sphinx* | | | MN312674<br>MN312827-828<br>MN312836-841<br>KU182961-962<br>KU182992-003 | |
| *Eonycteris spelaea* | | | MN312612-613<br>MN312616<br>MN312619<br>MN312622-623<br>MN312666-670 | |
| *Megaerops sp.* | | | KU182986 | |
| *Rousettus leschenaultii* | | | MG762654-656<br>MG762658<br>EF065513-514<br>KP895482-489<br>KP895524<br>KU182958-960<br>KU182974-985<br>KU182987-991 | |
| *Rousettus* sp. | | | MN312610<br>MN312614-615<br>MN312617-618<br>MN312620-621<br>MN312624<br>MN312627-631 | |
| **Hipposideridae** | | | | |
| *Aselliscus stoliczkanus* | KY417142<br>MN312601<br>MN312603<br>MN312607 | | | |
| *Hipposideros armiger* | KX285135-136<br>MN312598<br>MN312856 | | MN312860-861 | MN312817 |
| *Hipposideros pomona* | MN312658-659<br>MN312661 | | | MN312608-609<br>MN312625<br>KU343200 |
| *Hipposideros pratti* | MN312855 | | | MN312789-813<br>MN312863-865<br>KF636752 |
| **Rhinolophidae** | | | | |
| *Rhinolophus affinis* | MN312602<br>MN312634<br>KF569973<br>KF569996<br>KP876546 | | | |
| *Rhinolophus ferrumequinum* | KX285156<br>MN312832-835<br>MN312857<br>MN312867-869<br>DQ412042<br>DQ648856<br>KF294456<br>KJ473811-813<br>KP886808-809<br>KU182964<br>KY417145<br>KY770860 | | | |

| | |
|---|---|
| *Rhinolophus macrotis* | DQ412043<br>DQ648857 |
| *Rhinolophus pearsonii* | KF294442-443 |
| *Rhinolophus pusillus* | MN312652<br>MN312656<br>MN312660<br>MN312662-664<br>MN312742<br>MN312843-844<br>JX993987<br>KF294420<br>KF294422-424<br>KF294426-430<br>KF294434-438<br>KF294444-454<br>KF294457<br>KT381914 |
| *Rhinolophus rex* | KF294455 |
| *Rhinolophus sinicus* | KX285125<br>KX285134<br>KX285220<br>KY417151-152<br>MN312595-596<br>MN312600<br>MN312604-606<br>MN312637-645<br>MN312650<br>MN312653-654<br>MN312657<br>MN312688<br>MN312691-711<br>MN312738-741<br>MN312814-816<br>MN312818-821<br>MN312826<br>MN312829-831<br>MN312845-846<br>MN312858-859<br>DQ022305<br>DQ071615<br>DQ084199-200<br>DQ648795<br>FJ588686<br>GQ153539-548<br>KC881005-006<br>KF294421<br>KF294431-433<br>KF294440-441<br>KF367457<br>KJ473814-816<br>KT444582<br>KU182963<br>KX447563-565<br>KY417143-144<br>KY417146-152<br>KY770858-859 |
| *Rhinolophus thomasi* | KF294425 |
| *Rhinolophus* sp. | MN312632-633<br>MN312635-636<br>MN312671-673<br>MN312651 |

| | | | |
|---|---|---|---|
| | MN312655 | | |
| **Vespertilionidae** | | | |
| *Eptesicus serotinus* | | KY009613<br>KY009618-619<br>KY009621<br>KY009624<br>KY009626<br>KY009628<br>KY009632<br>KY009634 | |
| *Hypsugo* sp. | | KX285208<br>MN312842<br>MN312848-849<br>MN312852-854<br>KX285207<br>KX285209<br>MN312647<br>MN312850-851<br>KX442564 | |
| *Ia io* | | KX285195-196<br>KX285196<br>MN312847 | |
| *Myotis daubentonii* | | KU182956-957<br>KU182965<br>KU182972-973 | |
| *Myotis horsfieldii* | | | MN312680 |
| *Myotis pequinius* | | KY009614-615<br>KY009622 | |
| *Myotis ricketti* | | KX285219 | |
| *Pipistrellus abramus* | | KX285197<br>KX285199-200<br>MG021452<br>MN312646<br>MN312648-649<br>MN312665<br>MN312678-679<br>MN312681-683<br>MN312685-687<br>MN312866<br>DQ648809<br>KC522075-089<br>KJ473820 | MN312675<br>MN312677<br>MN312684 |
| *Pipistrellus pipistrellus* | | DQ648819 | |
| *Pipistrellus* sp. | | | MN312676 |
| *Scotophilus kuhlii* | | KX285160 | |
| *Tylonycteris pachypus* | | KX285157-159<br>KX285161-167<br>KY783855-864<br>KY783886<br>MN312689-690<br>MN312712-737<br>MN312743-748<br>MN312822-825<br>MN312862<br>DQ648794<br>KC522036-048<br>KJ473822<br>KX285160 | |

| | | | | |
|---|---|---|---|---|
| | | KX447544-558<br>KX447560-562<br>KY783889<br>KY783895<br>KY783902 | | |
| *Vespertilio sinensis* | | KX285223<br>MN312749-753<br>MN312753<br>MN312757-788<br>KJ473821 | | MN312754<br>MN312755<br>MN312756 |
